# Gradient matching accelerates mixed-effects inference for biochemical networks

**DOI:** 10.1101/2024.06.11.598320

**Authors:** Yulan B. van Oppen, Andreas Milias-Argeitis

## Abstract

Single-cell time series data frequently display considerable variability across a cell population. The current gold standard for inferring parameter distributions across cell populations is the Global Two Stage (GTS) approach for nonlinear mixed-effects (NLME) models. However, this method is computationally intensive, as it makes repeated use of non-convex optimization that in turn requires numerical integration of the underlying system. Here, we propose the Gradient Matching GTS (GMGTS) method as an efficient alternative to GTS. Gradient matching offers an integration-free approach to parameter estimation that is particularly powerful for dynamical systems that are linear in the unknown parameters, such as biochemical networks modeled by mass action kinetics. Here, we harness the power of gradient matching by integrating it into the GTS framework. To this end, we significantly expand the capabilities of gradient matching via uncertainty propagation calculations and the development of an iterative estimation scheme for partially observed systems. Through comparisons of GMGTS with GTS in different inference setups, we demonstrate that our method provides a significant computational advantage, thereby facilitating the use of complex NLME models in systems biology applications.

## INTRODUCTION

The development of experimental methods capable of tracking individual cellular responses in time [Spiller et al., 2010, Locke and Elowitz, 2009, Muzzey and Van Oudenaarden, 2009] has revealed extensive variability in cellular behaviors within isogenic cell populations [Levien et al., 2021, Pinheiro et al., 2022, Battich et al., 2015, Foreman and Wollman, 2020]. Variability within a biochemical system of interest can arise from the inherent stochasticity of chemical reactions that involve small numbers of molecules [Raj and Van Oudenaarden, 2008, Raser and O’shea, 2005, Paulsson, 2005], giving rise to fluctuations in the concentrations of the different interacting species. Variability extrinsic to the system of interest, such as cell-to-cell differences in growth rate, cell cycle stage, biosynthetic capacity, the cellular microenvironment, epigenetic states, and the abundance of species that are not explicitly accounted for, can also affect the observed dynamics [Huang, 2009, Kramer et al., 2022, Spencer et al., 2009, Keren et al., 2015, Guantes et al., 2015, Sherman et al., 2015, Zopf et al., 2013]. When the system of interest involves highly abundant species, intrinsic stochasticity is typically ignored, and the system is modeled deterministically via nonlinear ordinary differential equations (ODEs) [Aldridge et al., 2006, Albeck et al., 2008, Llamosi et al., 2016]. While individual cells may behave in a deterministic manner, extrinsic sources of variability may still produce heterogeneity at the population level. When these extrinsic sources vary slowly compared to the system dynamics, the ODE-based individual model can be augmented with randomly distributed parameters and/or initial conditions to model the collective dynamics of a heterogeneous cell population [Spencer et al., 2009, Albeck et al., 2008, Waldherr et al., 2009, Loos and Hasenauer, 2019].

Such *Nonlinear Mixed Effects (NLME) models* are routinely used to describe the responses of individuals in pharmacokinetics/pharmacodynamics studies [Davidian and Giltinan, 2003, Lindstrom and Bates, 1990]. However, their use in systems biology [Karlsson et al., 2015, Almquist et al., 2015, Fröhlich et al., 2018, Dharmarajan et al., 2019] has remained relatively limited due to the fact that inference of population parameters quickly becomes computationally intensive as both the size of the underlying ODE system (in terms of states and parameters) and the number of measured cells increase. For this reason, a common practice in the field has been to use averages of population measurements to obtain a point estimate of the model parameters. While it is often thought that this point estimate reflects the average of the population parameters, this is not the case, especially for biochemical models whose responses are highly nonlinear functions of their parameters [Bijman et al., 2021, Llamosi et al., 2016, Dharmarajan et al., 2019].

A number of inference methods for NLME models already exist in the pharmacometric literature [Kuhn and Lavielle, 2005, Davidian and Giltinan, 2003], and have recently started being applied to biochemical network inference [Dharmarajan et al., 2019, Llamosi et al., 2016]. The current gold standard for NLME inference is the Global Two Stage (GTS) [Davidian and Giltinan, 2017, Dharmarajan et al., 2019] method. This approach is applicable when a fair number of individual trajectories are available, and each trajectory is sampled sufficiently densely to reveal the underlying system dynamics. The GTS method begins by calculating cell-specific parameter estimates in the first stage and combines these estimates (and their associated uncertainties) to infer the population parameters via Expectation Maximization in the second stage. The second stage of GTS improves upon the “naïve” estimator that directly computes statistics of the population distribution from the individual estimates, leading to inflated covariance estimates. Although the GTS approach is highly reliable in general, the standard implementation of the first stage requires the use of non-convex optimization to identify cell-specific parameter estimates by minimizing the discrepancy between the measurements and the ODE solutions. This approach, which we will call *trajectory matching*, requires repeated numerical integration of the underlying ODE system to estimate the parameters of each cell. Moreover, to avoid local optima, the optimization algorithm needs to be started from multiple initial points, which further raises the computational cost of the method, especially as the size of the ODE system increases. Despite these bottlenecks, the GTS method remains quite popular, thanks to its straightforward implementation.

To avoid numerical integration of the system ODEs during parameter inference, an alternative class of methods, called *gradient matching* has been developed [Macdonald and Husmeier, 2015]. Instead of minimizing the discrepancy between measurements and ODE predictions, gradient matching minimizes the discrepancy between time derivatives predicted by the ODEs and those computed by smoothing the measurements. In this way, these methods can thought of as minimizing the difference between the two sides of the system ODEs. Several variants of gradient matching have been proposed over time [Varah, 1982, Ramsay et al., 2007, Calderhead et al., 2009, Dondelinger et al., 2013], with [Varah, 1982] being one of the first systematic treatments of the method.

Gradient matching is a good candidate for replacing the costly trajectory matching in the first stage of the GTS method, yet no gradient matching method has been used for the inference of NLME dynamical models so far. Here, we present the *Gradient Matching Global-Two Stage* (GMGTS) method for NLME dynamical systems, which constitutes the first implementation of gradient matching within the GTS framework. Besides providing point estimates of individual cell parameters, the first stage of GMGTS computes all the necessary uncertainty estimates for these parameters, which are subsequently fed into the second GTS stage. Although our method is applicable to general nonlinear ODE systems, it is particularly powerful when the ODE right-hand side is linear in the unknown parameters, as is the case for nonlinear models based on mass-action kinetics. For such systems, parameter estimation via gradient matching turns into a generalized least squares problem that can be solved analytically via regression methods. Moving beyond the standard gradient matching framework that requires full state information, GMGTS is also applicable to partially observed systems thanks to an iterative estimation scheme that retains computational efficiency compared to trajectory matching. Overall, GMGTS combines the strengths of gradient matching and the GTS method to provide sufficiently accurate population parameter estimates at a fraction of the time required by GTS. In this way, GMGTS increases the applicability of NLME parameter inference to computationally demanding problems comprising a large number of single-cell measurements and complex dynamical systems.

In the following, we start with a description of the GTS and GMGTS methods. We then provide a number of tests using *in silico* data to evaluate the applicability, efficiency, and accuracy of the GMGTS method in different settings and for different examples of biochemical systems. We show that the computational cost of GMGTS increases much more gradually with the number of free parameters within a fixed model structure compared to GTS. Gradient matching benefits greatly when the underlying system is linear in its parameters, and we therefore demonstrate how certain biochemical systems with nonlinear dependence on the parameters can be reformulated to satisfy this requirement. Finally, we show GMGTS is considerably more economical than the simulation-based GTS approach while providing sufficiently accurate estimates of population parameters within a realistic range of time step sizes and measurement noise levels. Having validated the GMGTS method in a simulation-based environment, we utilize it to infer parameter variability in a system describing fluorescent protein maturation in budding yeast, using single-cell experimental data for twelve commonly used fluorescent proteins [Guerra et al., 2022]. Our results suggest that maturation rates of fluorescent proteins display considerable variability across an isogenic yeast population, reflecting recent findings in mammalian cells [Wu et al., 2020].

As the development of advanced experimental methods increases the amount and complexity of single-cell data, we expect that NLME dynamical systems will become a “standard element” of the systems biology modeling toolbox in the future, similar to what ODE models have been in the past. Through the introduction of gradient matching into the NLME parameter estimation problem, we expect that our work will further stimulate and facilitate the use of this powerful class of models in systems biology applications.

## RESULTS

### GMGTS: seamless integration of gradient matching into the GTS framework

To capture the collective dynamic behavior of a heterogeneous cell population, we use an ODE-based *mixed effects model*. In this model, a set of ODEs describes the dynamics of the biochemical network of interest, while cell-to-cell variability is introduced by assuming that each cell “samples” a particular set of kinetic parameters from a multivariate probability distribution. More concretely, the dynamics of the *i*-th cell in a population of *N* cells is described by a *K*-dimensional ODE system (the *individual model*) of the form

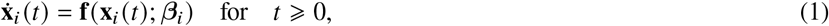

with initial condition 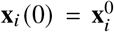. We further assume that the cell-specific vector of the kinetic parameters *β*_*i*_ is drawn from a multivariate normal distribution with mean **b** and covariance **D**; that is, *β*_*i*_ ∼ N (**b, D**) where **b** is a vector of *fixed effects* and **D** is the random effects covariance matrix. Note that **f** may depend on additional parameters that are fixed prior to inference. Therefore, *β*_*i*_ denotes only the set of kinetic parameters that are unknown.

We assume that each cell is observed at a discrete set of time points *t*_1_, … , *t*_*T*_ . The number and locations of the observation points may differ among cells, but we will assume that they are the same to simplify the notation since this assumption does not lead to any loss of generality. The experimental observations of the *i*-th cell are then related to the solution 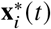 of (1) given *β*_*i*_ through the measurement model

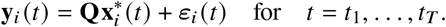

In this model, **Q** is a binary matrix that is used to select the measured components of 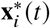. For instance, when all states are observed, **Q** is equal to the *K*-dimensional identity matrix. The measurement errors *ε*_*i*_ (*t*) are a combination of uncorrelated additive and multiplicative zero-mean Gaussian noise. Therefore, the *k*-th component of *ε*_*i*_ (*t*_*j*_), denoted *ε*_*ik*_ (*t* _*j*_ ), has mean 0 and variance 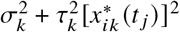. In this expression, *σ*_*k*_ and *τ*_*k*_ are some positive constants (the same for all cells), and 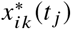 denotes the *k*-th component of the solution 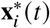 at time *t* _*j*_ .

Given the mixed-effects model presented above, the goal of the Global Two-Stage method (GTS) [Davidian and Giltinan, 2017] is to infer the *population parameters* **b** and **D** that give rise to the observed variability in single-cell trajectories. To achieve this, GTS proceeds in two steps: in the first stage, an individual kinetic parameter estimate and its associated uncertainty are obtained for each cell using generalized least squares (GLS) estimation. GLS also provides estimates of the measurement model parameters (*σ*_*k*_ and *τ*_*k*_) using the pooled residuals from all cells. In the second stage, estimates of the cell-specific parameters and their precisions are used to estimate the population parameters via an expectation-maximization scheme. Individual precision estimates are a critical input to the second stage since neglecting them leads to an inflation of the inferred random effects covariance [Davidian and Giltinan, 2017, Llamosi et al., 2016, Dharmarajan et al., 2019]. Besides providing population parameter estimates, the second stage of GTS refines the cell-specific parameter estimates based on the estimated population parameters. A detailed exposition of the GTS method implementation is provided in Methods.

The main computational bottleneck of the GTS method arises in the first stage. There, the estimation of cell-specific parameters is generally carried out by minimizing the discrepancy between single-cell measurements and model predictions with respect to the vector of kinetic parameters *β*_*i*_ (that is, by *trajectory matching*). This approach relies on non-convex optimization methods that often require multiple, randomly sampled starting points to avoid local minima in the parameter space. Moreover, the calculation of the objective function at each step of the optimization requires the numerical integration of (1). Non-convex optimization and ODE integration make the classical GTS method scale poorly with *N* (the number of observed cells), *K* (the dimension of (1)), and the size of the parameter vector *β*_*i*_.

To circumvent these limitations, the individual kinetics parameters in the first step of GTS can be obtained via *gradient matching*, an alternative approach to parameter estimation for ODE-based systems [Macdonald and Husmeier, 2015]. Instead of minimizing the discrepancy between the measured and predicted trajectories, gradient matching proceeds in two steps: first, the observed time series data are smoothed and the slopes of the smoothed time series are estimated at the measurement time points. Then, the discrepancy between the estimated slopes (representing the left-hand side of (1)) and the ODE-based time derivative (right-hand side of (1) evaluated at the measured states) is minimized with respect to the unknown parameters *β*_*i*_. By directly matching the two sides of (1), this parameter estimation approach does not require numerical solutions of ODEs.

Gradient matching is particularly powerful when the individual model (1) is linear in its parameters; that is, when the right-hand side of (1) can be decomposed as

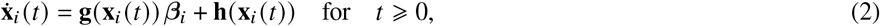

where **g**(·) is a matrix-valued function multiplying the parameter vector *β*_*i*_, **h**(·) is a vector-valued function, and both functions are independent of *β*_*i*_. In such a case, matching the two sides of (2) results in a linear regression problem that can be solved algebraically by means of (generalized) least-squares methods [Varah, 1982]. Linearity in parameters arises naturally in models based on mass action kinetics and, more generally, when the right-hand side of (1) is a polynomial function of **x**. As we will demonstrate below, systems with Michaelis-Menten or Hill kinetics can also be recast into a form that is linear in parameters by writing the elementary (mass-action) equations from which the Michaelis-Menten or Hill approximations were derived.

Combining gradient matching with the GTS method, this work presents the *Gradient Matching Global-Two Stage* (GMGTS) method for models of the form (2). In GMGTS, the time-consuming trajectory matching step in the first stage of the GTS method is replaced with gradient matching. In this way, the estimation of cell-specific parameters *β*_*i*_ is carried out with linear regression instead of non-convex optimization. By propagating the uncertainty of the measurement data to the smooth trajectory estimates and then to the cell-specific parameter estimates, the second stage of the GTS method is subsequently applied to obtain the population parameters (the vector of fixed effects **b** and the covariance of the random effects **D**) and refine the cell-specific parameter estimates.

Similarly to gradient matching, GMGTS starts by generating smooth estimates of the measured single-cell time series with the help of penalized B-splines (SI, § S4). We use the splines to approximate the system states and their time derivatives at the measurement time points of each cell. We then substitute **x**_*i*_ and 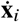 in (2) with the smoothed trajectory and derivative estimates, and formulate a generalized least-squares (GLS) problem to minimize the norm of the residuals Δ_*i*_ representing the difference between the two sides of (2) computed with the spline data. The covariance of the GLS residuals has an unknown structure that needs to be estimated iteratively to avoid biasing the parameter estimates. Approximating this covariance is not straightforward, since inserting the spline data (which carry uncertainty from the measurements) into (2) leads to a model with uncertain factors on both sides. A detailed treatment of the covariance matrix calculation is provided in SI, § S5. Upon convergence of the GLS, the precision of each estimate 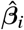 is approximated using the delta method [Asmussen and Glynn, 2007] (SI, § S5) and subsequently used in the second stage of the GTS method. Figure 1 shows a graphical overview of the GMGTS approach, and Methods provides a detailed treatment of the methodology.

**Figure 1:**
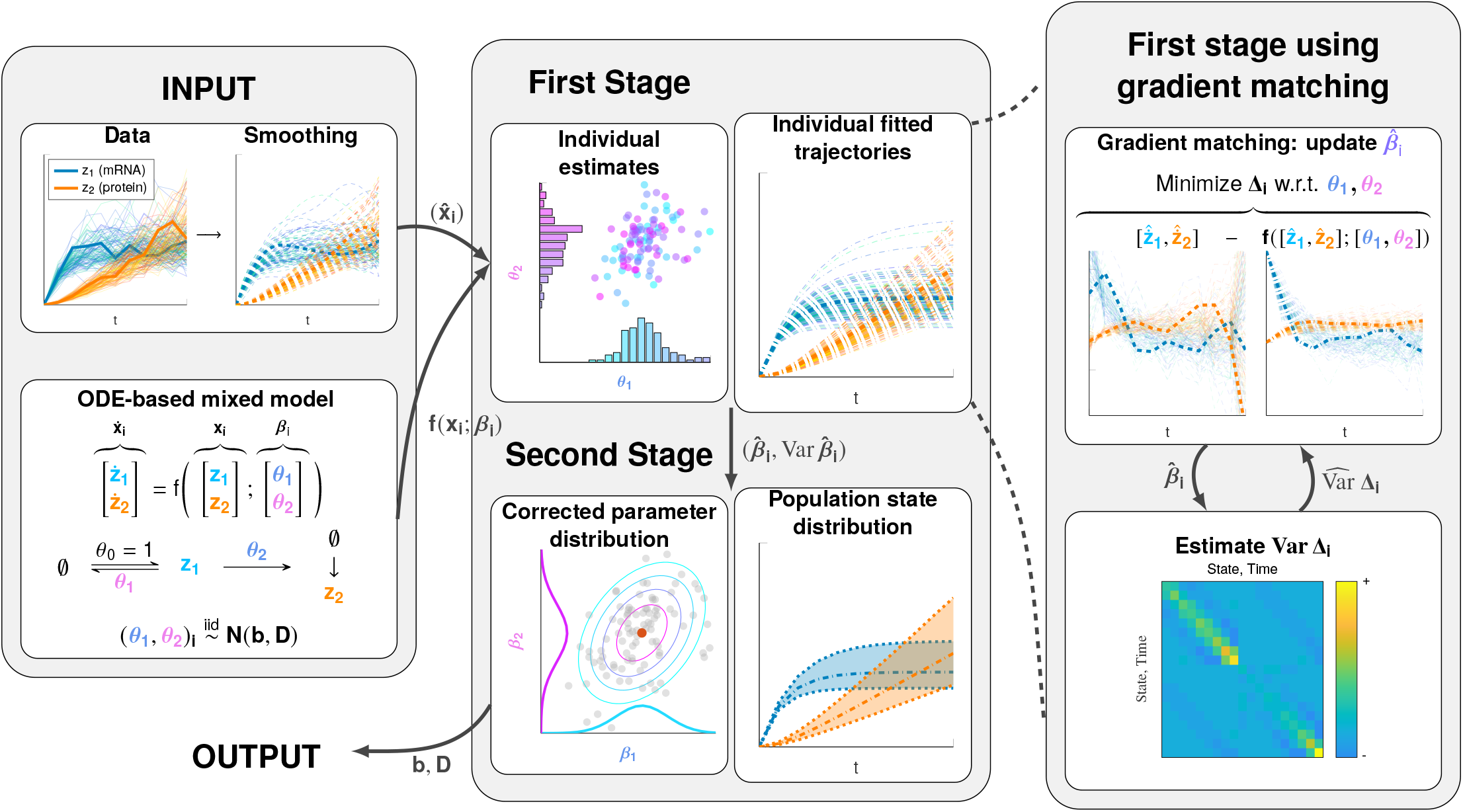
Graphical overview of the GMGTS method for fully observed systems. During initialization, the data are smoothed using penalized cubic B-splines (*left panel, top*) and are provided as input to the method along with an ODE-based model with randomly distributed parameters (*left panel, bottom*). The smoothed trajectories 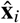 are used in the first stage (*middle panel, top*) to obtain individual parameter estimates 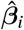 through gradient matching (*right panel, top*). Because of the linearity in parameters of the ODE system, gradient matching is carried out by solving a generalized least squares regression problem. This regression problem has a vector of residuals Δ_*i*_ with an unknown covariance matrix, which needs to be iteratively estimated (*right panel, bottom*). Upon convergence, uncertainties of the individual parameter estimates 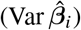 are computed by propagating the uncertainty from the single-cell data through the spline estimates. The parameter estimates 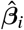 and their corresponding uncertainties are used in the second stage (*middle panel, bottom*), where the population parameter estimates 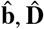 are obtained.

### GMGTS can utilize partial state information

By construction, the original gradient matching method requires measurements of the entire state vector to evaluate the model gradients. However, it is not always feasible to observe all states of a given biochemical system. For these cases, we propose a modification to the gradient matching method which estimates the hidden states and their gradients and correctly handles the uncertainties that enter the Generalized Least Squares (GLS) problem for individual parameter estimation. Iterative numerical integration of the individual model (2) is inevitable in that case, but the number of integration runs is massively reduced compared to trajectory matching.

To numerically integrate the individual model at the start of the iterative procedure, an initial guess for the individual parameter values is required. This guess can be based on prior knowledge, but we obtain it by fitting the individual model (2) to the average population trajectories using maximum likelihood estimation. An iterative optimization scheme is subsequently applied to estimate the individual parameter vector *β*_*i*_ for each cell. More concretely, for every cell we perform the following operations:

#### [0. Initialization]

Smooth the observed data **y**_*i*_ and differentiate the smoothed data to obtain estimates of the observed states and gradients. Integrate the system using the initial parameter guess to obtain estimates of the hidden states. Substitute the numerically integrated solution into the right-hand side of the ODE system to obtain hidden gradient predictions, and initialize the random-effects covariance matrix.

#### [1. Gradient matching]

With full state and gradient information, optimize the parameter vector estimate 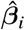 through gradient matching using GLS.

#### [2. Estimation of hidden states]

With the updated individual parameter estimate 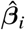 , integrate the system to update the hidden state and gradient predictions.

#### [3. GLS residual covariance]

Approximate the uncertainty of the individual parameter estimate and use it to approximate the covariance matrix of the residuals Δ_*i*_ of the next GLS iteration. Repeat steps 1–3 until convergence.

Figure 2 provides a graphical overview of the estimation procedure for partially observed systems. To compute the GLS estimate of individual parameters at the gradient matching step, we need an estimate of the residual covariance matrix. The approximation of this matrix is significantly more elaborate in the partially observed case compared to the case with full state information since the uncertainty also needs to be propagated via the hidden states. The details of these calculations are presented in Methods and the SI, § S6.

**Figure 2:**
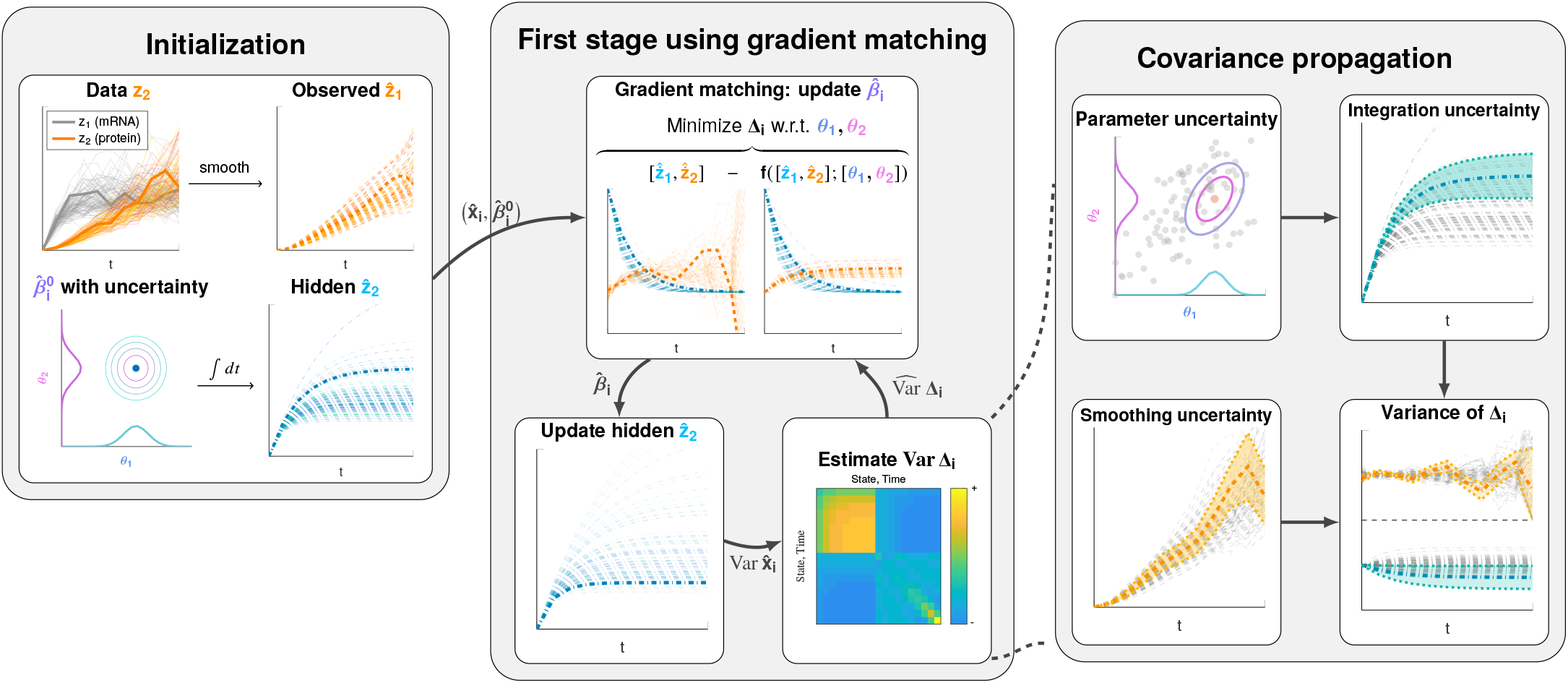
Graphical overview of the GMGTS method modifications to the first stage for partially observed systems. Besides smoothing the measured states, an initial parameter guess 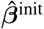 is required to initialize the hidden state estimates by numerically integrating the ODE system (*left panel*). With full state information now available, an extended iterative scheme comprises the modified first stage of GMGTS. In this scheme, an individual parameter estimate 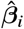 is obtained via gradient matching using generalized linear regression (*middle panel, top*). Numerical integration of the ODE system yields updated hidden state estimates (*middle panel, bottom left*), whose uncertainty needs to be considered together with the smoothing uncertainty to approximate the residual covariance matrix Var Δ_*i*_ (*middle panel, bottom right*). The smoothing uncertainty affects the residual covariances directly but also propagates through the updated parameter estimates to the hidden states via the numerical ODE solutions (*right panel*).

### GMGTS achieves high accuracy across a range of sampling intervals and noise levels

Given that GMGTS relies on time series smoothing, we first tested how the accuracy of the estimated random effects distribution depends on measurement noise levels and the number of measurement time points. For these tests, we used two example systems under different scenarios of data sparsity and noisiness.

Our first example system is a model of a bifunctional two-component signaling cascade [Kurdyaeva and Milias-Argeitis, 2021] comprising two proteins: a histidine kinase *H* and a response regulator *R*. In response to an external input, the histidine kinase autophosphorylates (*H*_*p*_) and transfers its phosphate group to the response regulator (*R*_*p*_). At the same time, the non-phosphorylated form of the bifunctional kinase (*H*) acts as a phosphatase towards *R*_*p*_. The system comprises six nonlinear ODEs with polynomial right-hand sides that are linear in parameters. Full model details are provided in Methods. Besides the fully observed case, we also treat the case where only the total phosphorylated kinase (*H*_*p*_) [Duvall and Childers, 2020] and the response regulator (*R*_*p*_) [Butcher and Tabor, 2022] are observed, while the other states are hidden. To avoid loss of structural identifiability, we fixed one rate constant in each pair of reverse reactions (*k*_1_ = *k*_2_ = *k*_6_ = 0.5 and *k*_4_ = 1.5), and let *k*_3_, *k*_5_, *k*_7_, *k*_8_ vary across a population according to normal distributions with means 0.5, 0.1, 0.8, 0.2 and a common coefficient of variation equal to 0.25.

Our second example is a model of fluorescent protein synthesis and maturation in budding yeast [Guerra et al., 2022]. In this system, transcription of the protein is activated in a step-wise manner at *t* = 0, and immature (“dark”) protein (*D*) is translated from mRNA (*M*) with time delay *d*. The dark protein matures into fluorescent protein (*F*) at a rate that depends on the maturation rate constant *k*_*m*_. Finally, all system components are subjected to dilution due to cell growth. The dynamics of this system is described by a system of delay differential equations provided in Methods.

We considered both fully and partially observed cases, with measurements of *F* being available in the latter. To avoid loss of structural identifiability [Guerra et al., 2022], the dilution rate was fixed to *k*_dil_ = 0.004 min^−1^ and the mRNA synthesis rate was set to *k*_*r*_ = 0.1 transcripts per minute. According to measurements presented in [Guerra et al., 2022], the time delay *d* was fixed at 4 min. Since our tests indicated that *k*_dr_ is weakly identifiable when it is held equal for all cells, we assumed that it is fixed at a *k*_dr_ = 0.07 min^−1^. This rate corresponds to an mRNA half-life of around 10 minutes which is in line with previous estimates for budding yeast [Neymotin et al., 2016]. We further assumed that *k*_p_, *k*_m_ follow normal distributions across a cell population, with respective means 0.025, 0.05 and coefficients of variation equal to 0.25.

To understand how the inferred random effects distributions depend on the quality and quantity of measurement data in both the GTS and GMGTS methods, we generated noisy measurement time series (between *t* = 0 and *t* = 100 min for the two-component system and between *t* = 0 and *t* = 200 min for the protein maturation model) while varying the number of time points *T* and the multiplicative error factor *τ*. For each combination of settings (number of time points and noise magnitude), we generated 10 independent samples of individual parameters assuming a population of *N* = 100 measured cells each repetition and monitored how the random effects distribution is recovered by the GMGTS and GTS methods. Figure 3 summarizes the resulting mismatch between ground truth (data-generating) and estimated distributions in terms of the Wasserstein distance [Villani, 2009]. Overall, with the exception of overly sparse measurements, the accuracy of the GMGTS distribution estimates is comparable to that of GTS and displays good robustness to measurement noise. For a visual accuracy comparison, Figure S1 and Figure S2 show representative examples of the inferred random effects distributions and the predicted state distributions for the two-component system and the maturation model. Although the additional loss in accuracy incurred by the GMGTS method is small, it is significantly more computationally efficient than GTS, which takes 15–30 times longer for the two-component system and 10–15 times longer for the maturation model. See Panel A of Figure 4 for a complete computing time comparison of the GMGTS versus the GTS method for both systems.

**Figure 3:**
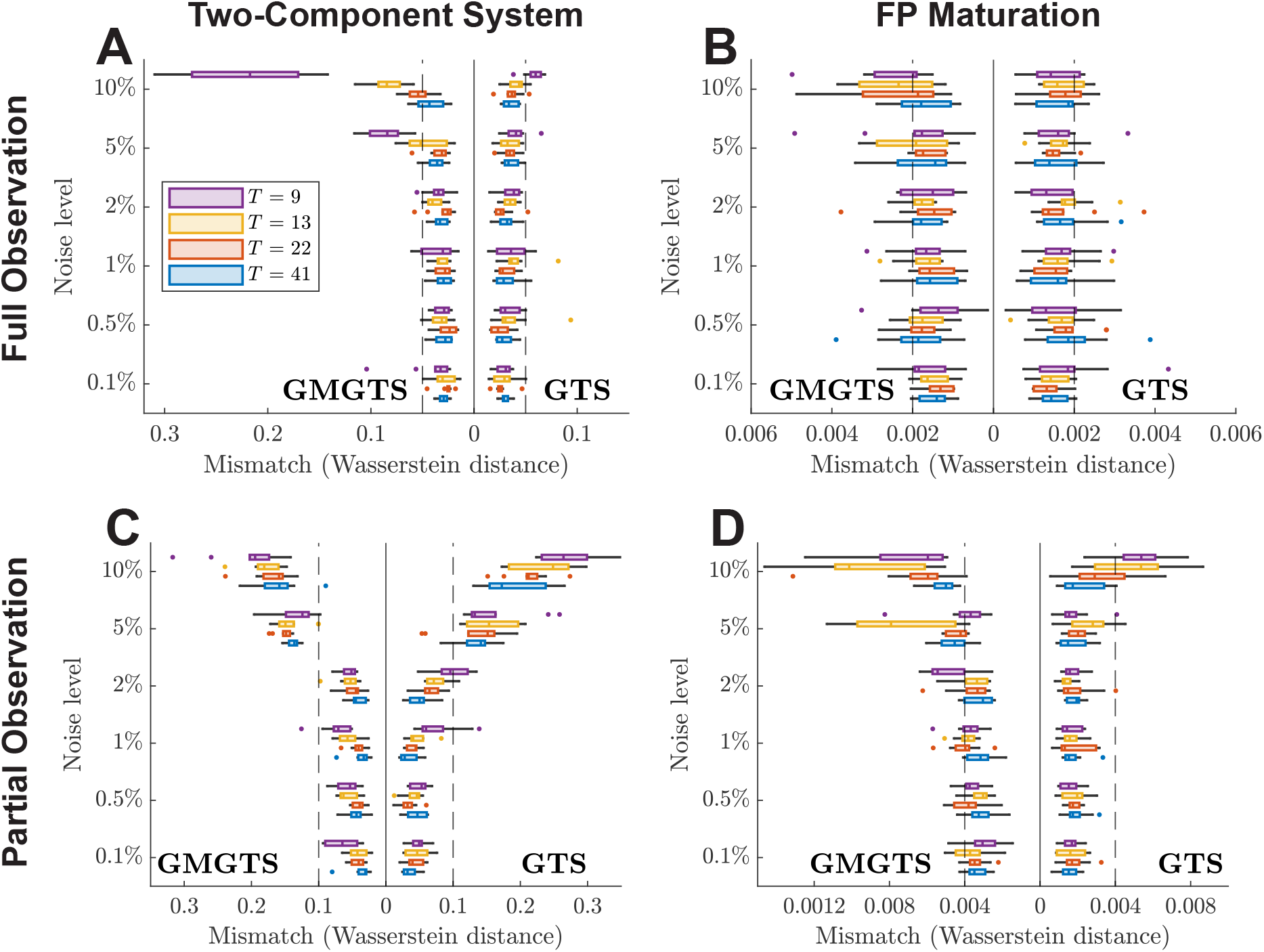
Accuracy comparison between GTS and GMGTS for two-component and FP maturation systems. The multiplicative noise level (*τ, y*-axis) and the number of simulated measurement time points (*T* , color labels) were varied to test the accuracy of each method. For each setup, 10 independent sets of individual parameters and corresponding measurements were sampled and used for inference of the random effects distributions. The boxplots summarize the mismatch between the inferred and the data-generating random effects distributions in terms of Wasserstein distances. Points lying further than 1.5 times the interquartile range of each box are considered outliers and are displayed as dots. **A** and **C**. Results for the bifunctional two-component signaling cascade. **B** and **D**. Results for the one-step fluorescent protein expression and maturation model. For all but the noisiest and sparsest measurements, the additional accuracy loss of GMGTS is small compared to that of GTS.

**Figure 4:**
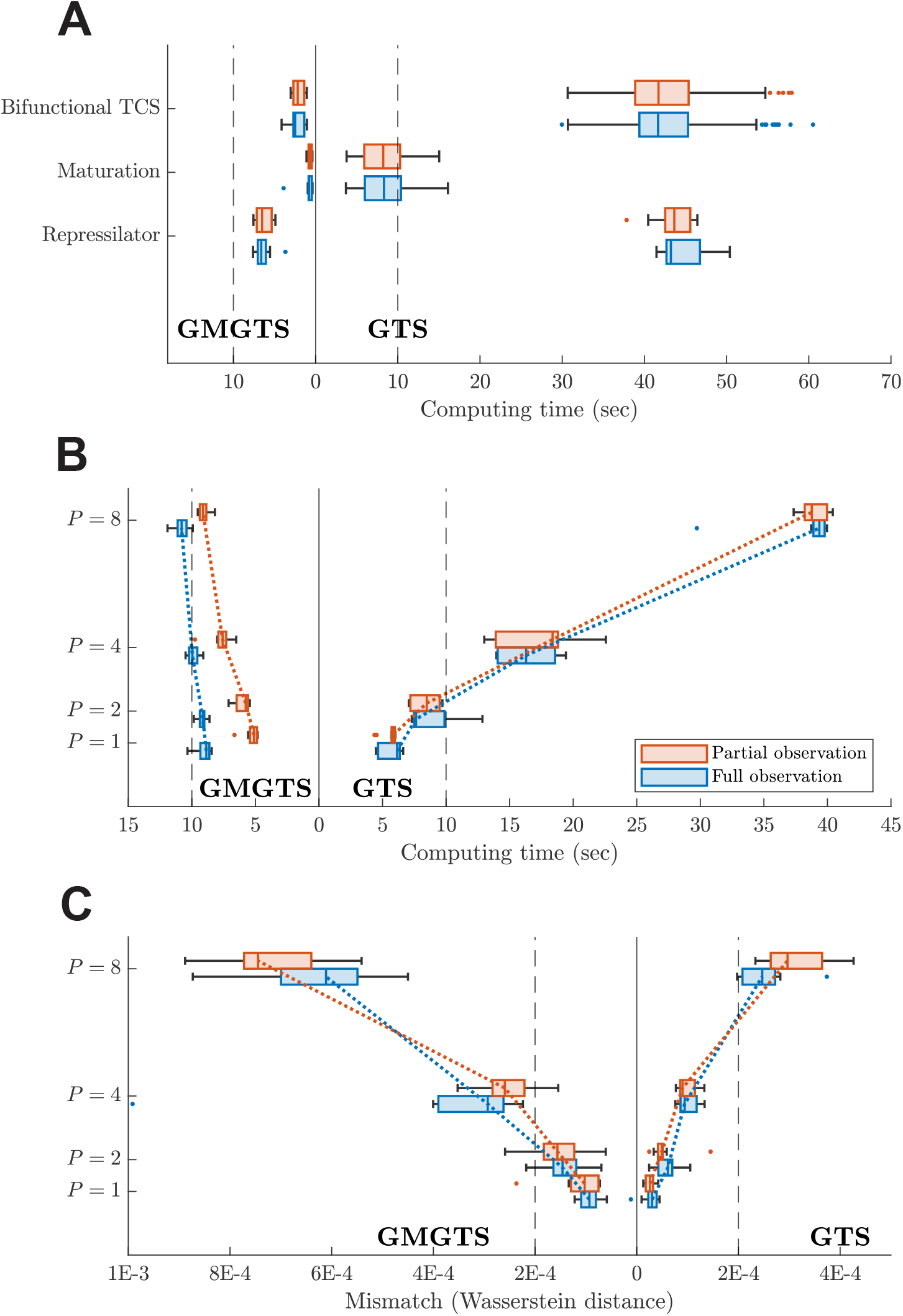
Computational efficiency comparison between GMGTS and GTS in different simulation studies. Computing times and inaccuracies of the inferred population distributions were obtained using simulated data as ground truth, and were summarized using boxplots. In each test setup, both GMGTS and GTS were applied on fully and partially observed systems. Points lying further than 1.5 times the interquartile range of each box are considered outliers and are displayed as dots. **A**. Summary of computing times for multiple data generation and inference runs (240 for the two-component system and the maturation model, 10 for the repressilator) for the mechanistic models presented above. The GMGTS method provides a 6- to 30-fold speedup compared to GTS, both with full and partial state information. Note that for the fully observed repressilator, the additional protein-DNA complexes are taken as observed states as well. **B**. Computational efficiency comparison between the two methods with respect to the number of free parameters (*P*) in a fixed Lotka-Volterra model structure. For each value of *P*, 10 independent data samples and corresponding parameter distributions were generated and inferred. In the partially observed case, the first 4 states were assumed to be hidden. **C**. Accuracy comparison between the two methods for the simulation setup considered in Panel B.

### Casting systems into mass action form enables parameter inference via gradient matching using linear regression

When the individual ODE model is linear in the unknown parameters, the gradient matching step of GMGTS is carried out via linear regression, which is considerably faster than numerical optimization. Biochemical models derived from the law of mass action automatically fulfill this requirement. On the other hand, models with nonlinear dependence on parameters are often based on quasi-steady-state approximations of mass action kinetics. Therefore, it is sometimes possible to expand these models so that they become linear in parameters. In such cases, GMGTS can still be used to infer the population parameters in a computationally efficient manner.

To illustrate this point, we consider the mathematical model of the repressilator, a synthetic gene network that exhibits self-sustaining oscillations via delayed negative feedback [Elowitz and Leibler, 2000]. The system consists of three transcriptional repressors arranged in a ring topology so that each protein represses the expression of the next and is repressed by the previous one.

Its dynamics is described by six coupled ODEs

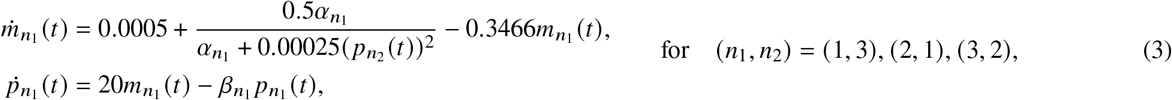

where *m*_*n*_ and *p*_*n*_ denote the respective mRNA and protein concentrations (µM) of the *n*-th repressor (*n* = 1, 2, 3). In this example, we assume that parameters *α*_*n*_, *β*_*n*_ (*n* = 1, 2, 3) vary across a cell population according to a normal distribution with means 0.16 µM and 0.0693 min^−1^ respectively. Moreover, each parameter has a coefficient of variation equal to 0.1, and each pair (*α*_*n*_, *β*_*n*_) has a correlation coefficient *ρ* = 0.5. With these settings, the system displays sustained oscillations in protein and mRNA concentrations across a large range of random effect values.

As can be seen in (3), the model equations contain Hill functions in which unknown parameters can also appear in the denominators of fractions. Since these Hill functions have been derived from quasi-steady-state approximations of protein-DNA interactions, (3) can be expanded with additional states (protein-DNA complexes) to produce a system that describes mass-action kinetics, and is therefore linear in parameters. The derivation of the expanded system can be found in Methods. With this expansion, we naturally assume that the additional protein-DNA complexes are hidden states.

To compare the performance of the GMGTS and GTS methods on the inference of the random effects distribution for this example system, we sampled *N* = 100 individual parameter vectors (corresponding to 100 cells), and generated measurements for each *m*_*n*_ and *p*_*n*_, assuming a 5-min sampling interval over a period of 100 minutes and multiplicative measurement noise (*τ* = 5%). The whole inference process was repeated ten times with independently sampled datasets, using the expanded model (Methods) for both GTS and GMGTS. Figure S3 shows the parameter and state distributions for the first repressor fitted by the GMGTS and the GTS methods in a representative inference run. Both approaches provide adequate population parameter estimates and predicted population trajectories, but the time required by the GTS method is 6 to 8 times longer compared to the GMGTS method (cf. Panel A of Figure 4).

### The computational cost of GMGTS scales modestly with the number free parameters

Parameter estimates obtained via gradient matching may be less precise than those obtained by trajectory matching, but the former method can be considerably faster when the individual model is linear in the unknown parameters. Similarly, the main idea behind GMGTS is that a significant gain in speed can be achieved at a relatively small possible loss in accuracy compared to GTS. The goal of this section is to explore this trade-off more systematically.

To begin with, Panel A of Figure 4 summarizes the computation times for the simulation studies presented so far. As can be seen, the GMGTS method offers a 5- to 20-fold speedup over GTS. To better understand this behavior, we tested how the computational cost and accuracy of GMGTS and GTS scale with the number of free parameters in a given model structure. To this end, we considered a generalized Lotka-Volterra system with 16 species (*x*_*n*_, *n* = 1, … 16) governed by the following set of differential equations

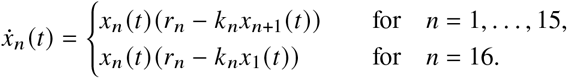

In these equations, the *n* th birth rate *r*_*n*_ was fixed at *n* 80 for all species, and a subset of the *k*_*n*_ parameters were assumed to be variable with mean 0.02 and standard deviation 0.005. The number *m* of free parameters was fixed at *m* = 1, 2, 4, 8 by setting 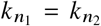 when *n*_1_ ≡ *n*_2_ mod *m* (e.g., *k*_1_ = *k*_3_ = · · · , when *m* = 2). We considered both fully observed and partially observed systems with the first four states assumed to be hidden in the latter case. For each simulation setting, we sampled *N* = 100 vectors of individual random effects (*k*_1_, … , *k*_*m*_) from the random effects distribution described above, and generated the corresponding measured time series assuming multiplicative measurement noise (*τ* = 5%) and fixed measurement time points *t* = 0, 1, 2, … , 20. For both GMGTS and GTS, we recorded the computation time and the Wasserstein distance of the estimated random effects distribution from the ground truth. Panel B of Figure 4 depicts the comparison of computing times, showing that the cost of GTS scales quite steeply with the number of free parameters. On the other hand, the cost of GMGTS shows only a modest increase with the number of free of free parameters. At the same time, Panel C of Figure 4 shows that the inaccuracy of GMGTS increases faster compared with the inaccuracy of GTS, though the inaccuracies of both methods are very small in absolute terms.

### Fluorescent protein maturation rates show considerable variability across budding yeast populations

The maturation rate of a fluorescent protein (FP) limits the temporal resolution at which dynamical intracellular processes can be monitored and is a major determinant of *in vivo* FP brightness in fast-dividing cells [Balleza et al., 2018, Guerra et al., 2022]. For example, in order to monitor transient gene expression changes during the cell cycle of fast-dividing cells, FPs need to have a sufficiently short maturation half-times compared to the typical cell cycle duration of those cells [Guerra et al., 2022]. Comprehensive studies of FP maturation are still scarce and are challenged by the fact that maturation rates often differ across different organisms and growth conditions, and also between *in vivo* and *in vitro* experiments. Moreover, whereas maturation rates are typically reported using population-average measurements, recent work in mammalian cells [Wu et al., 2020] has uncovered significant cell-to-cell variability in FP maturation rates across isogenic mammalian cells. The origins of this variability are unknown, but it has been conjectured that they are caused by cell-to-cell differences in the intracellular redox state [Wu et al., 2020].

Moving beyond the simulation studies presented above, we asked to what extent cell-to-cell variability affects FP maturation kinetics in *Saccharomyces cerevisiae* (budding yeast). In our previous work [Guerra et al., 2022], we presented a systematic *in vivo* characterization of maturation for a set of twelve FPs that are commonly used in this organism. Our experimental data comprised single-cell time course measurements of FP accumulation upon step-wise induction of protein expression with an optogenetic system. FP maturation rates were then inferred by fitting a mathematical model to the *averages* of the single-cell traces. While the use of averaged data allowed us to align and compare our maturation rate estimates with previous work, the datasets of [Guerra et al., 2022] contained single-cell information that could provide insights into the variability of maturation rates across cell populations. We therefore applied our GMGTS method to infer mixed-effects models of FP expression and maturation for the same set of twelve FPs. Similarly to [Guerra et al., 2022], we considered models with one- and two-step FP maturation kinetics, to account for proteins with one or two rate-limiting maturation steps. In either model, immature (“dark”) protein (*D* or *D*_1_) is synthesized from mRNA (*M*) with time delay *d*, and mRNA is produced and degraded at respective rates *k*_r_ and *k*_dr_. In the one-step model, *D* matures into mature fluorescent protein (*F*) with a maturation rate *k*_m_. In the two-step model, *D*_1_ first transitions into an intermediate (non-fluorescent) form (*D*_2_) at rate *k*_m1_ before maturing into *F* at rate *k*_m2_. *k*_m1_ and *k*_m2_ cannot be inferred individually given only fluorescence data, but replacing them with a single parameter (i.e. assuming *k*_*m*_ := *k*_m1_ = *k*_m2_) introduces negligible error [Guerra et al., 2022], so the same approach is followed here. Furthermore, all system components are diluted due to cell growth. Panels A and B of Figure 5 contain schematics of the reactions of the two models. The delay differential equations describing the dynamics of each model can be found in the Methods.

**Figure 5:**
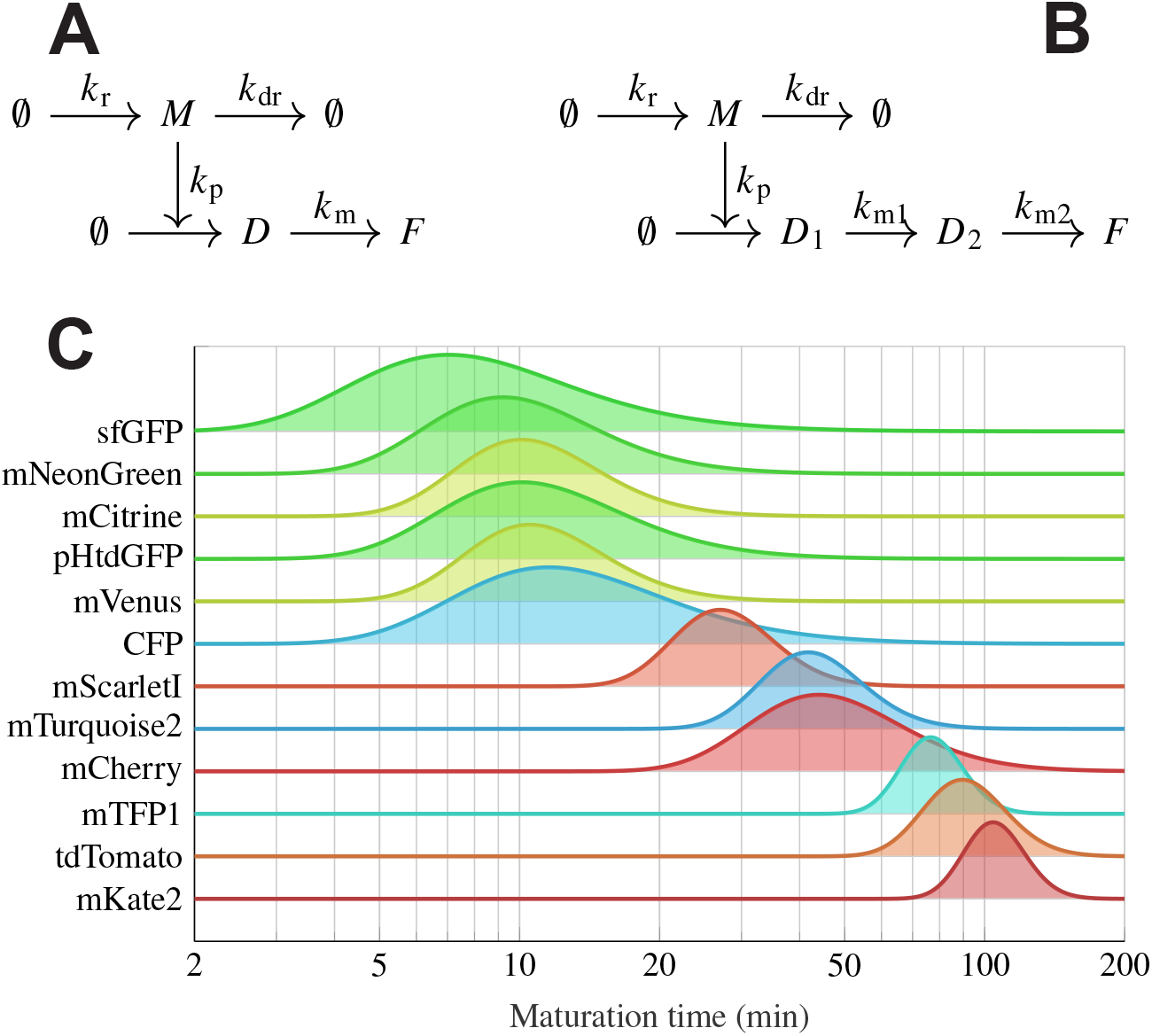
Schematic representation of the mathematical models describing FP maturation, and inferred distributions of maturation half-times. **A–B** Schematics of the one-step (left) and two-step (right) models. Dark protein (*D* or *D*_1_) is synthesized from mRNA (*M*) and matures into *F* (via *D*_2_ in the two-step model). Each state is diluted at a common rate due to cell growth (omitted here for clarity). **C** Estimated maturation half-time distributions across the different tested FPs. Half-time distributions were computed from the corresponding distributions of maturation rate estimates (Methods), and are ordered by their respective means. For two-step proteins, an approximate maturation half-time was derived analytically (Methods). Single-step maturing FPs were estimated to be considerably faster, and their maturation rates displayed larger variability compared to proteins undergoing two-step maturation (cf. Table T3).

Given that only the concentration of *F* can be measured by fluorescence microscopy, we assumed that states *M* and *D* (*D*_1_ and *D*_2_ in the two-step model) are unobserved. To ensure structural identifiability of the models, the average dilution rate due to cell growth was estimated by tracking cell volume measurements over time [Guerra et al., 2022], and was fixed to an experiment-specific value for each protein (*k*_dil_ ≈ 0.0045 min^−1^). Given the fact that the product *k*_r_*k*_p_ is identifiable but the individual synthesis rates are not [Guerra et al., 2022], we fixed *k*_r_ and let *k*_p_ be variable. Finally, given the poor practical identifiability of *k*_dr_ (cf. simulation study above), we fixed it at 0.07 min^−1^, which is within the range of previously obtained mRNA degradation rate estimates for GFP in budding yeast [Neymotin et al., 2016]. With these choices, the rate of protein synthesis *k*_p_ (a measurement scaling factor) and the maturation rate(s) *k*_m_ were left to be estimated from the data.

To understand how our simplifying modeling assumption of a common dilution rate for all cells could affect parameter inference, we generated *in silico* measurement data, assuming that the dilution rate has a known mean (as in our model), but is also affected by cell-to-cell variability with a coefficient of variation of 0.05, a value that was determined experimentally in [Guerra et al., 2022]. We then used a model with a fixed (i.e. non-variable) dilution rate to infer the random effects distribution of *k* _*p*_ and *k*_m_, assuming a coefficient of variation of 0.25 for the maturation rate. Table T1 summarizes the inferred maturation rate medians and quartiles from ten independent runs of data generation and inference. These results suggest that neglecting the variability of the dilution rate during inference does not significantly inflate the variability of maturation rate estimates, as long as the maturation rate variability is considerably larger than the variability in dilution rates.

After some simple preprocessing steps of the model (described in Methods), we estimated the population distributions of (*k*_p_, *k*_m_) for each of the 12 FPs quantified in [Guerra et al., 2022] using the GMGTS approach. Panel C of Figure 5 summarizes the estimated maturation half-time distributions, ordered with respect to their means. A description of how these distributions are obtained from maturation rates is provided in Methods, and an approximation of the maturation half-time for two-step maturation models is derived in Methods. Figure S5 to Figure S16 show the full random effects distributions and the measurements overlayed with the predicted state distributions, as inferred by the GMGTS method for each FP.

As can be seen from Figure 5, our estimates confirm the empirical observation that proteins that mature in a single step are faster than proteins with two-step maturation. The means of the population distributions are largely in line with those obtained in [Guerra et al., 2022] (cf. Table T3). Similarly to [Wu et al., 2020], several FPs displayed considerable intercellular variability in terms of their maturation half-times. In particular, sfGFP and the sfGFP-based pH-tdGFP show increased variability compared to the other FPs. Moreover, Table T3 indicates a tendency of fast-maturing proteins to vary more in terms of single-cell maturation rates compared to slow-maturing proteins.

To further verify whether a variable maturation rate is supported by the data, we performed a statistical comparison of a model with a fixed maturation rate against the model with a variable maturation rate considered above. To obtain the former model, we fixed the maturation rate of each FP to its average (cf. Table T3) and inferred *k* _*p*_ from the single-cell data. We then calculated and compared the marginal likelihoods of the two models, listed in Table T2. For every FP, the model with fixed maturation rate had a substantially lower log-likelihood compared to the model with variable maturation, suggesting that the data do not support fixed maturation rates. Considering the potential complications arising from variability in maturation rates in single-cell fluorescence studies, these results warrant additional experimental validation.

## DISCUSSION

Dynamic NLME models offer a computationally efficient alternative to the fully stochastic models of intracellular dynamics, especially when the main sources of cell-to-cell variability are extrinsic to the system of interest and vary slowly compared to the system dynamics. However, inference of population parameters for this class of models remains a computationally intensive task that had received relatively little interest in the biological literature until recently. While an increasing number of inference methods are now becoming available, these methods do not take advantage of the fact that the nonlinear ODEs modeling biochemical reaction networks often depend linearly on the unknown parameters. Exploiting this linearity can lead to computationally efficient inference methods, which are particularly useful when alternative model structures need to be tested during the model-building process or in a model-selection setting.

To speed up the inference of dynamic NLME models, we presented the Gradient Matching Global Two-Stage (GMGTS) method, an adaptation of the well-established Global Two-Stage (GTS) approach [Davidian and Giltinan, 2017]. GMGTS relies on gradient matching to obtain first-stage individual parameter estimates which are subsequently used in the second stage to estimate the population parameters via Expectation-Maximization. When the underlying dynamical system is linear in parameters, the first stage of GMGTS solves a linear regression problem (generalized least squares) that replaces the non-convex optimization required by the more commonly used trajectory matching approach. Thanks to the speed of GMGTS, tuning of the method is quite straightforward since the user gets near-instant feedback.

We demonstrated the computational efficiency of GMGTS over GTS in various *in silico* tests and showed that GMGTS maintains a satisfactory level of accuracy in all cases. Having established the performance of the method on simulated data, we applied it to recover the maturation times of fluorescent proteins from experimentally determined fluorescence time series of single budding yeast cells. Our findings are largely in line with current literature in terms of the mean maturation times [Guerra et al., 2022], but we additionally observed that the maturation rates of certain fluorescent proteins show considerable variability across isogenic populations. Due to their implications for single-cell fluorescence microscopy experiments, these results warrant further investigation, potentially via an orthogonal experimental method.

Besides enabling the use of gradient matching for mixed-effects model inference, GMGTS addresses two common shortcomings of gradient matching methods. The first is the fact most gradient matching methods require full state information in order to calculate the time derivative of each state since standard gradient matching approaches do not make use of numerical ODE integration. However, this reliance can be a limitation, given that current experimental methods typically do not permit the simultaneous quantification of many readouts within the same cell and across time. To address this issue, we developed an algorithm that treats unobserved states as latent variables and estimates them iteratively together with the system parameters. This scheme relies on a small number of numerical integration runs ( ∼5-10 per iteration per cell) that is insignificant compared to the hundreds of runs that trajectory matching generates for each initial point of the optimization. A second weak point of gradient matching approaches is that they provide point parameter estimates and no parameter uncertainties. However, these uncertainties are required for inferring the population parameters in the second stage of GMGTS. We have addressed this issue for both fully and partially observed systems, by propagating the measurement variability to the individual parameter estimates via the estimated states and their gradients.

Overall, our gradient matching approach is based on and extends that of [Varah, 1982]. Alternative solutions to the gradient matching problem have been proposed more recently, such as gradient matching using Gaussian processes (GPs) [Calderhead et al., 2009, Dondelinger et al., 2013, Barber and Wang, 2014] and the generalized profiling method [Ramsay et al., 2007]. While these methods are also able to handle unobserved states, to the best of our knowledge, no parameter uncertainty computations have been provided for them. Moreover, GP-based approaches require the estimation of posterior distributions whose evaluation is not analytically tractable and instead necessitates computationally intensive sampling schemes. While generalized profiling does not suffer from this limitation, it needs to solve a non-convex optimization problem over a large number of parameters. This is because the method optimizes the parameters of the smoothed state estimates and the ODEs simultaneously, and the right-hand sides of ODE systems are typically nonlinear when considered as joint functions of states and parameters. Smoothing the experimental data using penalized cubic B-splines prior to gradient matching (as we do here) means that the smooth estimates are not constrained by the ODE structure. However, our approach provides a significant computational edge to GMGTS at the cost of a manageable loss in estimation accuracy and allows us to properly account for the uncertainty in parameter estimates.

Since smoothing precedes parameter estimation in GMGTS, careful smoothing of the measurements is important for obtaining accurate estimates of population distributions. Unfortunately, a fully automated method for time series smoothing cannot exist, and some trial and error is still required. To facilitate and speed up this process, we have implemented an interactive smoothing app [van Oppen, 2023] to provide immediate visual feedback on the choice of the B-spline knot locations, the interval of curvature penalization, and the penalty multiplier on the smooth trajectory estimates. The smooth estimates obtained via the app are directly fed into GMGTS, making it easier to assess individual parameter estimates obtained with different smoothing settings. Since gradient matching only requires the smoothed trajectories, their gradients, and the covariances of these estimates, any model other than penalized B-splines can be interchangeably used at the smoothing step. In this way, the GMGTS approach can be seamlessly adapted to other types of (penalized) splines, Gaussian process-based smoothing, or support vector machine regression.

Besides uncertainty in parameterization, biochemical systems may also display variability in their initial conditions. As [Varah, 1982] suggested for the fully observed case, an additional fine-tuning step could follow gradient matching to estimate the initial conditions in isolation via trajectory matching. A filtering approach [Särkkä and Svensson, 2023] could be employed to infer initial conditions for partially observed systems.

On the output side, our current implementation of GMGTS is based on the assumption that the uncertainty of individual parameters follows a (multivariate) normal distribution across a cell population. Though biochemical network parameters are often non-negative, this assumption suffices if their coefficient of variation is not too large. Note that the normality assumption is imposed in the second stage of GMGTS and GTS (i.e. the EM algorithm), and that the first stage is distribution-agnostic. Postulating a non-normal distribution for the individual parameters is also possible, but this modeling assumption would require more complex deconvolution methods [Efron, 2016] in the second stage of GMGTS (and GTS). An alternative for modeling non-negative parameters with the standard form of GMGTS (and GTS) is to work in the log-parameter space and assume that the parameter logarithms are normally distributed [Dharmarajan et al., 2019] - an approach that amounts to postulating log-normal individual parameter distributions. Although this operation sacrifices the linearity of the ODE system in the original parameters, GMGTS can still be used to avoid the repeated numerical integration of the system.

In summary, GMGTS is a highly efficient and powerful inference method for dynamic nonlinear mixed-effects models that is also highly extensible, serving as the basis for the development of more powerful and general inference methods in the future.

## ACKNOWLEDGEMENTS

Y.B.O. and A.M.-A. were supported by the Netherlands Organization for Scientific Research (NWO) through a VIDI grant to A.M.-A. (project number 016.Vidi.189.116).

## AUTHOR CONTRIBUTIONS

Conceptualization, Y.B.O. and A.M.-A.; Methodology, Y.B.O. and A.M.-A.; Software, Y.B.O; Visualization, Y.B.O.; Formal Analysis, Y.B.O. and A.M.-A.; Writing - original draft, Y.B.O.; Writing - review and editing, Y.B.O. and A.M.-A.; Funding acquisition, Supervision, and Project administration, A.M.-A.

## SUPPLEMENTAL INFORMATION

Supplemental Information contains 3 tables (T1-T3), 17 figures (S1-S17), mathematical derivations, and software implementation details.

## METHODS

### METHOD DETAILS

#### Model

Consider a population of *N* cells, each having an underlying mechanistic model given by a *K*-dimensional ODE system

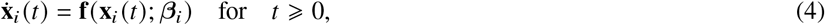

where **f** : ℝ^*K*^ × ℝ^*P*^ → ℝ^*K*^ is common for all cells. The dynamics of each cell are specified by an unknown associated *P*-dimensional parameter vector *β*_*i*_, assumed to be drawn from a normal distribution with mean vector **b** and covariance matrix **D**, so

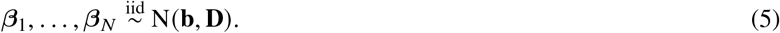

Equations (4) and (5) comprise an *ODE-based mixed-effects* model, where *β*_1_, … , *β*_*N*_ are called *random effect* vectors, **b** is the *fixed effect* vector, and **D** is the *random effects covariance matrix*.

The time series data for each cell *i* are assumed to be a *T*-dimensional vector of noisy samples **x**^*^ (*t*; *β*_*i*_), the solution to (4), i.e.,

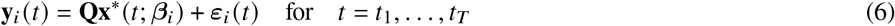

In (6), **Q** is a binary matrix used to select the observed components of 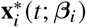 and ***ε***_*i*_ *t* is a vector containing the measurement noise for each state. We consider the measurement dimension to be common across cells, but accommodating different measurement times for each cell is straightforward. Not all states are necessarily observed in practice, so assume without loss of generality only the first *L ⩽ K* states are measured, i.e., **Q** consists of the first *L* rows of the *K*-dimensional identity matrix. The methodology described in the following sections readily extends to (possibly nonlinear) measurement functions **Q** : ℝ^*K*^ → ℝ^*L*^ of the underlying process **x**_*i*_ (*t*) with a differentiable partial inverse, such as a partially invertible linear map.

The measurement errors ***ε***_*i*_ (*t*_*j*_) are assumed to be a combination of additive and multiplicative noise, and hence have components *ε*_*ik*_ (*t*_*j*_) independently distributed as

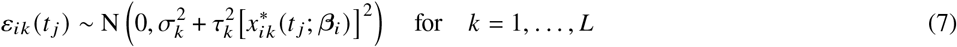

for some *σ*_*k*_, *τ*_*k*_ *⩾* 0 taken to be common across cells, and 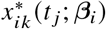 denotes the *k*-th component of 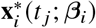.

In the context of gradient matching (discussed in more detail in a later section), we focus on the special case where **f** is linear in the unknown parameters *β*_*i*_. In that case, (4) can be decomposed as

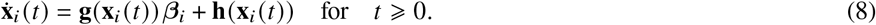

Here, **g** : ℝ^*K*^ → ℝ^*K*^ × ℝ^*P*^ and **h** : ℝ^*K*^ → ℝ^*K*^ are both common across cells and independent of *β*_*i*_, but neither is restricted to be linear in **x**_*i*_ (*t*).

#### Global Two-Stage (GTS) aproach for nonlinear mixed-effects models

The current gold standard for inference in ODE-based mixed-effects models is the *global two-stage* (GTS) approach [Davidian and Giltinan, 2017, Ch. 5]. Individual estimates of each *β*_*i*_ are obtained in the first stage, along with estimates of their associated precisions 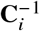. In the second stage, the random effects distribution is inferred from the estimated *β*_*i*_’s, taking into account their covariances **C**_*i*_. Specifically, for the model described by (4)–(7), GTS proceeds as follows:

I. For each *i* = 1, … , *N*, estimate *β*_*i*_ by minimizing the *weighted sum of squares* (WSS)

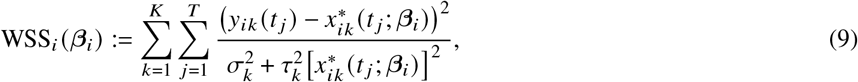

where *y*_*ik*_(*t*_*j*_) denotes the *k*-th component of **y**_*ik*_(*t*_*j*_). In practice, the measurement model parameters *σ*_*k*_, *τ*_*k*_ are unknown and need to be iteratively estimated together with the system parameters. This joint estimation problem is solved with a *feasible weighted least squares* (FWLS) algorithm [Greene, 2003, Ch. 10]. Given an estimate 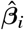 of *β*_*i*_, consistent estimates of the measurement model parameters can be obtained by maximizing the following log-likelihood function for each state:

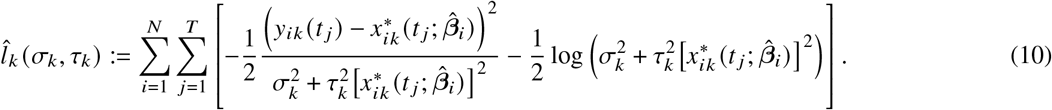 Let 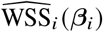 denote WSS_*i*_ (*β*_*i*_) when *σ*_*k*_ and *τ*_*k*_ are substituted by their estimators. Each estimate of *β*_*i*_ may then be obtained by means of FWLS using the following scheme:
  1. Initialize variance parameter, e.g., each 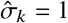 and 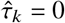.
  2. Update 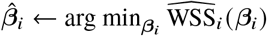 using the current variance parameter estimates 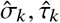 cf. (9).
  3. Update 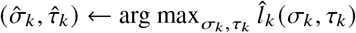 using the current parameter estimate 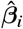 ; cf. (10);
  4. Repeat steps 2 and 3 until convergence.
  Estimates 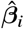 are produced for each cell *i* = 1, … , *N*. Subsequently, each error covariance matrix **C**_*i*_ is estimated using the Hessian of 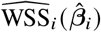 , i.e.,

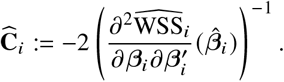
II. Infer the population parameters **b** and **D** via maximum likelihood using an *expectation maximization* (EM) algorithm. In this algorithm, the random effects *β*_*i*_ are considered as latent variables, and their estimates 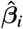 from the first stage are treated as data. As stated in [Davidian and Giltinan, 2017, § 5.3.2], using an asymptotic approximation as the number of measured time points (*T* ) increases, we have

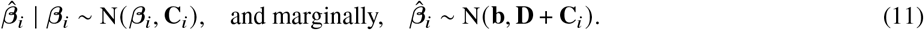

We initialize the estimates 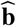 and 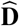 using the sample mean and covariance of 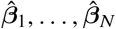 . Treating *β*_*i*_ as latent variables, their estimates 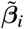 are initialized at their respective first stage estimate 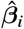 . At the *expectation step* (E-step), we compute the expectation of *β*_*i*_ conditional on 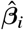 and the current estimates 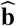 and 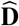 for each cell. This means we update (see SI, § S3 for the derivation)

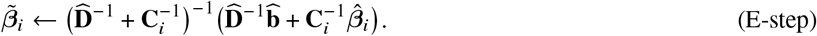

Next, we update 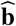 and 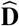 given their current values and each updated 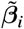 in the *maximization step* (M-step) as

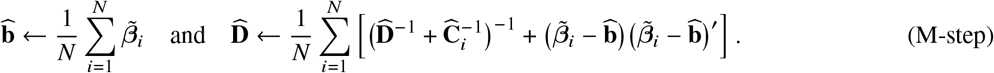

The EM algorithm iterates (E-step) and (M-step) until convergence. The refined estimates 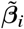 are called *empirical Bayes predictors*, and correspond to modified first-stage estimates that are shrunk toward the population mean. In this work, we disregard these estimates and focus on the estimated population parameters 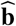 and 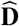 and random effects 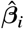 .

##### Remark 1.

Note that each evaluation of 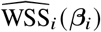 generally requires a numerical solution to system (4) to be computed. The frequent inclusion of nonlinearity in processes characterized by ODEs is the reason the optimization of the cell-specific model parameters is typically nonconvex.

##### Remark 2.

The first stage of the method does not depend on the assumption (5) that random effects follow a multivariate normal distribution in the population. However, the normality assumption becomes important in the second stage, where the population parameters are inferred. Using another distributional assumption for the random effects would necessitate the solution of a deconvolution problem, which is technically challenging. For this reason, the normality assumption (5) is standard in the mixed-effects literature.

#### The Gradient Matching Global Two-Stage (GMGTS) approach

Gradient matching [Varah, 1982] can be applied to any nonlinear dynamical system, but it is particularly efficient for systems that are linear in parameters (8). GMGTS replaces trajectory matching with gradient matching in the first step of the GTS method. Using the uncertainty estimates for the cell-specific estimates produced in the first stage, the second stage remains unmodified. This section provides detailed methodology corresponding to the case of full state information, so *L* = *K*. A similar procedure is required for partially observed systems but some modifications are needed to estimate the hidden states, which are presented in the next section. In brief, the following steps comprise the first stage of the GMGTS method:

I.A Smooth the measurements *y*_*ik*_ (*t*) with a penalized B-spline 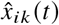 using FWLS for each state *k* = 1, … , *K*, with weights dependent on the estimated measurement error variance. The details are given in SI, § S4.
I.B Use the splines estimated in Step I.A to approximate (8), where the left-hand side is obtained by differentiating the splines, while the right-hand side is produced from substitution of the splines into **g**(·) and **h**(·). Denoting the spline approximations by 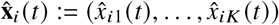 for *t*_1_ *⩽ t ⩽ t*_*T*_ , we introduce these splines in (8) with the goal of finding *β*_*i*_ so that

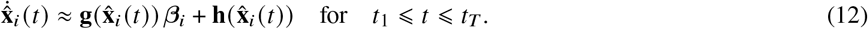

Since *β*_*i*_ is assumed to be independent of *t*, we can stack the matrices containing the spline data in (12) to obtain

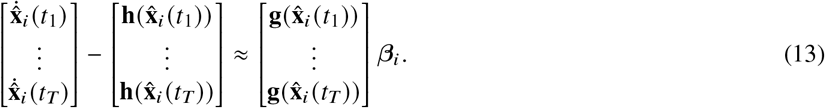

The arrangement (13) facilitates the estimation of *β*_*i*_ by means of convex optimization, such as generalized linear regression. The challenge lies in deriving a suitable error model for the residuals of (13).

The remainder of this section is dedicated to the details of the gradient matching performed in Step I.B. Since the model parameters for different cells are optimized independently at this stage, we drop the subscripts *i* to simplify the notation, writing *β* := *β*_*i*_ and **x** := **x**_*i*_ and following the same convention for estimates or components of these vectors. Denote the stacked matrices of spline approximations 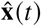, their derivatives 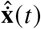, and function evaluations 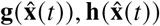 as

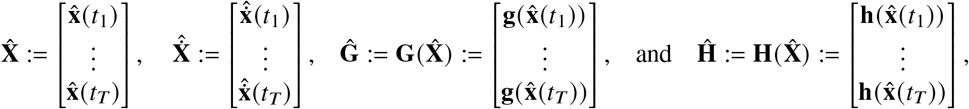

allowing us to transform (13) into a formal model

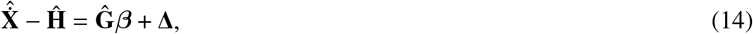

where Δ is the vector of residuals. Equation (14) is a generalized linear regression model with response vector 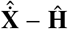 and design matrix 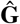 . The main difficulty of this model is the fact that the covariance structure of the residuals is not straightforward to obtain. We address this problem by calculating delta-method approximation **V**(*β*) of the covariance matrix of Δ conditional on *β*, as described in SI, § S5. The respective estimates 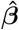 and 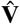 of the model parameters and the residual covariance matrix need to be computed iteratively in a *feasible generalized least squares* (FGLS) algorithm:

1. Initialize 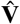 at the *T K*-dimensional identity matrix.
2. Given the current value of 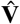, update the model parameter estimate 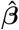 by minimizing the generalized sum of squares criterion

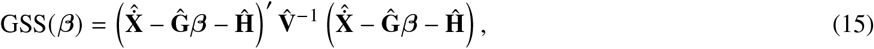 To satisfy parameter bounds, minimize (15) using quadratic programming. In case the optimum is not found on the boundary of the parameter space (see Remark 5), it coincides with the generalized least squares estimate of *β*:

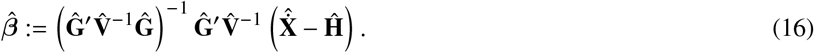
3. Update 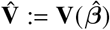 according to (S7) using the updated estimate 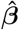.
4. Repeat Step 2 and Step 3 until convergence (see SI, § S8 for the details).

Since the 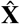 appears on the right-hand side of the generalized linear model, using the Fisher information matrix corresponding to (15) to approximate the uncertainty of the final estimate 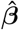 leads to an upward bias in terms of precision. Rather, we estimate its uncertainty using a delta-method approximation as

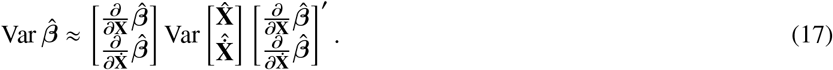

The derivations of 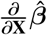 and 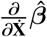 are provided in SI, § S5.

##### Remark 3.

In case of large differences in state magnitudes, 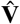 may alternatively be initialized as a diagonal matrix with elements inversely proportional to squared mean state values.

##### Remark 4.

We retained the *t*_1_, … , *t*_*T*_ in this section for notational convenience. However, in case the measured time series are sampled too densely, it is worth considering a smaller set of measurement time points that still adequately capture the dynamics of the system. This choice offers a significant reduction in computational cost since GMGTS performs a large number of operations on large unstructured data matrices. Because the data are smoothed prior to gradient matching, the gradient matching time points can be freely chosen anywhere within the smoothing interval.

##### Remark 5.

It should be noted that estimates on the boundaries of the parameter space are generally undesirable. When this happens for some cells, we halt the FGLS iterations and omit the corresponding estimates from the second stage. In case a large fraction of cell-specific parameters are estimated at the boundary, we can either extend the search space or consider a suitable parameter prior, cf. SI, § S7.

#### GMGTS for partially observed systems

In the case of partial observation, without loss of generality, we assume that the first *L* < *K* states are measured. The overall process of smoothing followed by gradient matching presented for fully observed systems is preserved. However, since gradient matching requires full state information, we need to estimate the hidden states using the ODE solution given the current parameter estimate 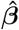. That is, we obtain predictions

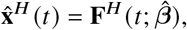

Where 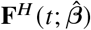 is the solution of the hidden states of the dynamical system. As before, we smooth the *L* observed states with penalized B-splines 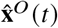. Both vector components can then be combined into full state and gradient estimates

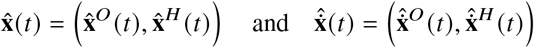

to perform gradient matching as before. Specifically, given the smoothed observed data, we need to approximate the model parameter vector *β* and the hidden states 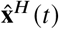 using an iterative procedure:

1. Initialize 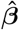 at the same point 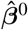 for all cells (e.g., from prior knowledge or by fitting the average of the single-cell measurements) and numerically integrate the system using the initial value of 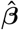. The hidden state predictions are obtained directly from the numerical solution, and the gradient predictions are obtained by substituting the solution into the right-hand side of the ODE. The uncertainty of the initial estimate 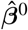 needs to be prescribed as well, see Remark S2. With this uncertainty, initialize 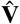 using (S17).
2. Given the full state estimate 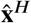 and the error covariance matrix 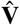 , update 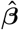 by minimizing (15) as above.
3. Numerically integrate the system using the updated value of 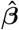 to obtain updated hidden state and gradient predictions.
4. Update 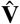 by propagating the uncertainty from the splines via 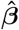 to the hidden state components of 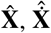 , and subsequently to 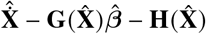 (see SI, § S6).
5. Repeat Steps 2–4 until convergence (see SI, § S8 for the details).

As with full state information, the uncertainty of 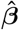 should be approximated using (17) upon convergence. In the partially observed case, this quantity is already calculated during the calculation of 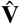 .

#### Implementation of the GMGTS method

The GMGTS method is implemented in MATLAB using IQM Tools with Monolix for numerical integration [van Oppen, 2023]. Due to the fact that smoothing precedes parameter estimation in GMGTS, high-quality smoothing of the measurements is essential for obtaining accurate estimates of population distributions. However, a fully automated method for high-quality time series smoothing cannot exist, and some trial and error is still required on the user side. To facilitate and speed up this process, we implemented an interactive smoothing app to provide immediate visual feedback on the choice of the B-spline knot locations, the interval of curvature penalization, and the penalty multiplier on the smooth trajectory estimates. The smooth estimates obtained via the app are directly fed into the gradient matching algorithm, making it easier to test the fidelity of the individual parameter estimates obtained with different smoothing settings. Our smoothing app, as well as the GMGTS code, are provided in [van Oppen, 2023]. Extensive implementation details are listed in SI, § S8.

#### Preprocessing of dynamical systems in simulation studies

##### Reformulation of the repressilator equations

To reformulate (3) into an ODE system with linear dependence in parameters, we consider a plausible set of elementary reactions that can produce the rational terms in (3) via a quasi-steady-state approximation. In these reactions, a monomer of repressor *n*_1_ (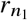, where *n*_1_ = 1, 2 or 3) forms a dimer 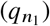 and binds to a free promoter of repressor 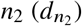 to form a DNA-protein complex 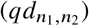. Using a quasi-equilibrium approximation for the dimerization reaction, we can write the dimer concentration as a function of the total repressor concentration 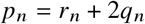. Concretely, using the dissociation constant *K*_*d*_ of the dimerization reaction and the law of mass action to write the equilibrium condition, we get

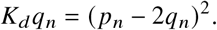

This equation can be solved in terms of *q*_*n*_ to find a function that connects *q*_*n*_ to the total concentration *p*_*n*_. Setting *K*_*d*_ = 4000 µM, the physically meaningful solution of the quadratic equation is

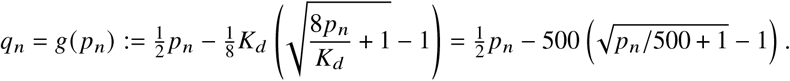

Finally, we rewrite (3) by explicitly modeling the binding of the dimer 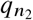 to the promoter 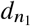 , assuming that the rate constant of the binding reaction is equal to 1 µM^−1^min^−1^ and the rate constant of the unbinding reaction is 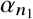 :

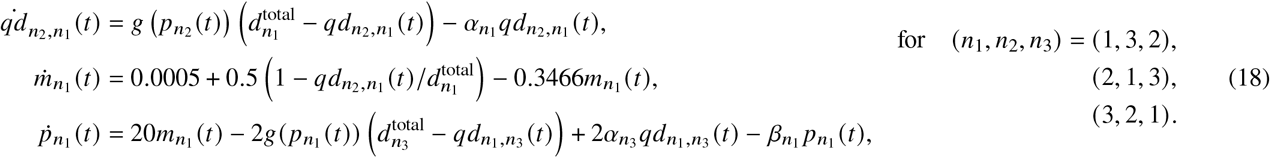

In these equations, 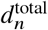 is the total concentration of DNA that repressor *n* can bind. This concentration is set to 1.0µM for each repressor gene, which is much smaller than the concentration of the repressor itself, so that the dynamics of repressor concentration remains roughly the same. Note that the right-hand side of the system is nonlinear in each *p*_*n*_ but linear in each *α*_*n*_, *β*_*n*_. Since the complexes 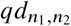 are generally not observed, we treat these states as hidden in the partially observed case.

The system (18) can be used to infer random effects distributions of each *α*_*n*_, *β*_*n*_ using GMGTS. To see how it is equivalent to (3) assume that the DNA-protein complexes reach a fast equilibrium. This means that the first ODE in (18) is at equilibrium, and the DNA-repressor complex varies according to the total amount of repressor, implying

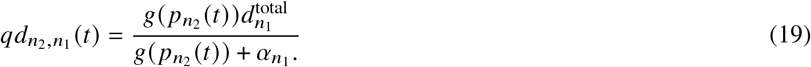

Using the fact that the dimer and monomer also reach a fast equilibrium, we can substitute the dimer abundance with the monomer squared divided by *K*_*d*_. That is,

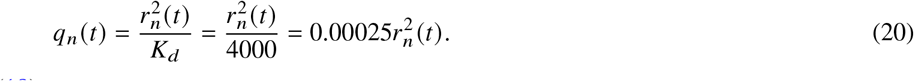

Substituting relations (19) and (20) into (18), we get

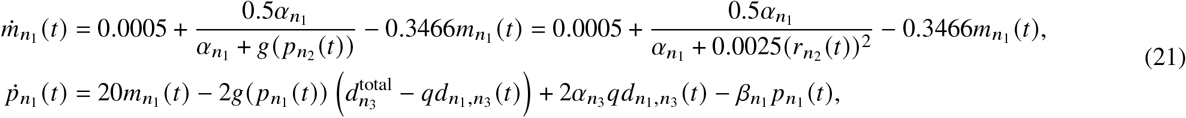

Given the large dissociation constant *K*_*d*_ of the dimerization reaction and the range of repressor concentrations reached for the nominal parameter values, it is safe to assume that 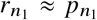 in (21) to arrive at the differential equations for mRNA in (3). Furthermore, given the small concentration of DNA compared to the repressor concentrations, the middle terms in the protein equation have only a very small impact on the protein dynamics. Overall, (21) produce mRNA and protein trajectories that are very close to those of the reduced system, as can be verified by simulation of the two systems. See Figure S4 for data simulated from both systems for a sample of sets of parameters.

##### Bifunctional two-component system dynamics expressed in terms of the total abundances of phosphorylated H and R

As discussed in [Kurdyaeva and Milias-Argeitis, 2021], the dynamics of the bifunctional two-component system is described by a six-dimensional set of ordinary differential equations. Omitting the time argument for clarity, these equations are

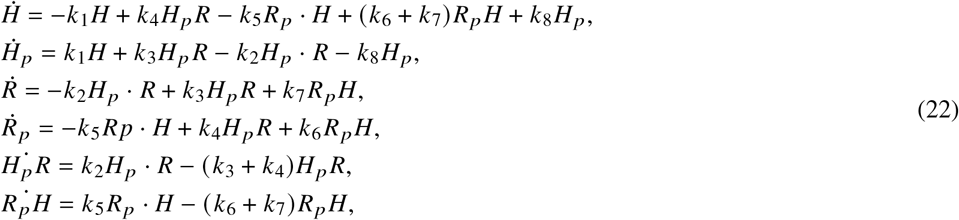

where *H* and *H*_*p*_ denote the unphosphorylated and phosphorylated histidine kinase respectively, and a similar convention is used for the response regulator *R. H*_*p*_ *R* and *R*_*p*_*H* are the complexes formed by the binding of the kinase *H*_*p*_ to *R* and the phosphatase *H* to *R*_*p*_. We will express (22) in terms of quantities that can be observed experimentally, such as the total amounts of phosphorylated *H* and *R*, denoted by

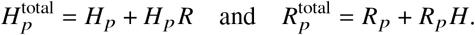

Using that 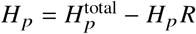 and 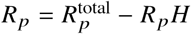, we can rewrite (22) as

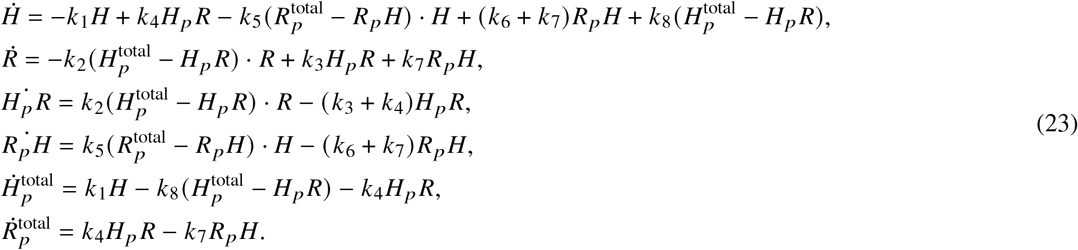

For the example we present in Results, we assumed that only 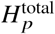 and 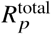 are observable, with all other states being hidden. To prevent a lack of identifiability, we also assumed that the reaction rate constants for one direction of each reaction were fixed, with the rate constant of the other direction being free.

##### Approximation of the delay-differential equations for one- and two-step fluorescent protein maturation with an ODE system

The kinetic models of one- and two-step fluorescent protein maturation used in [Guerra et al., 2022] are described in terms of delay differential equations to account for the delays in transcription and translation. In these models, mRNA transcription is activated by an optogenetic system at time *t* = 0. mRNA (*M*) starts appearing *τ* minutes later, and undergoes degradation at a fixed rate per molecule. Dark (immature) fluorescent protein (*D*) is translated from mRNA with a time delay *d*, and eventually matures into fluorescent protein (*F*). All system components are diluted due to cell growth. These processes are described by the following set of delay differential equations:

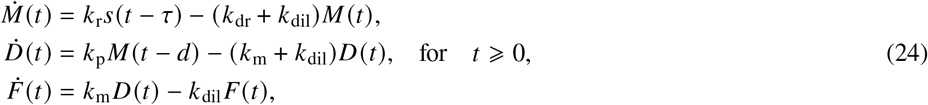

where *s* : ℝ → {0, 1} denotes the Heaviside step function. In the two-step maturation model, dark protein (*D*_1_) goes through a second non-fluorescent form (*D*_2_) before maturing into *F*. The analogous delay differential equations are

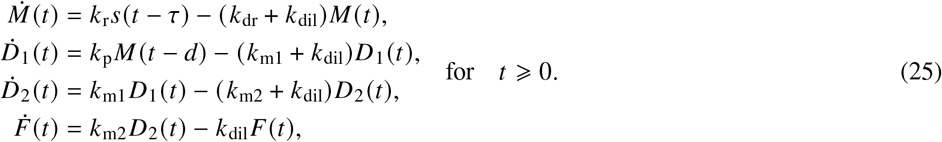

In either system, only the concentration of *F* can be measured by means of fluorescence microscopy, so we assume that states *M* and *D* (or *D*_1_ and *D*_2_) are unobserved. When *F* is measured, the transcriptional delay *τ* can easily be ignored by redefining the time axis of the system, so that time point zero of (24) and (25) coincides with time point *τ* of the measurements. However, the delay of translation requires additional treatment.

Since the mRNA synthesis rate *k*_r_ and the dilution rate *k*_dil_ have already been fixed, additionally fixing the mRNA degradation rate *k*_dr_ effectively turns *M* into an input function to the dark protein equation. The solution of the mRNA differential equation with *M* (0) = 0 is readily verified to be

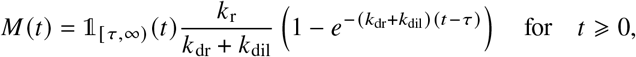

where 𝟙_*A*_ : ℝ → {0, 1} denotes the indicator function on an interval *A*. We can then address the second time delay (in the dark protein equation) by substituting the delayed solution *M* (*t* − *d*) into the differential equation for *D*, yielding (analogously for *D*_1_)

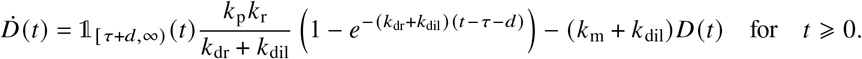

However, because *τ* and *d* only appear in the input function *M* (*t* − *d*), they vanish when we redefine the time axis in terms of *u* := *t* − *τ* − *d*. Note this operation of shifting the time axis means that time point zero of (26) and (27) corresponds with time point *τ* + *d* of the measurements. Accordingly, we obtain a delay-free two-dimensional system

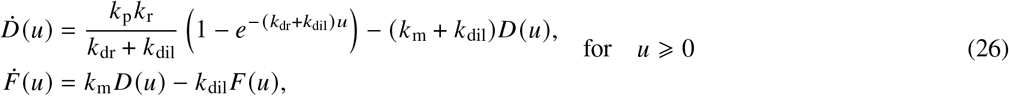

for the one-step maturation model, and a three-dimensional system

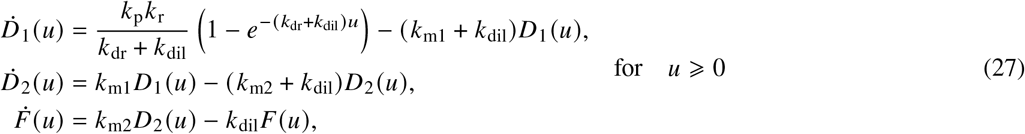

for the two-step maturation model.

##### Preprocessing of experimental data obtained through fluorescence microscopy

Prior to inference, the measurements for each fluorescent protein need to be preprocessed. The protein concentrations are computed from the total fluorescent channel intensity divided by the corresponding volume per cell. By the nature of the optogenetic stimulation, mRNA synthesis starts with a delay of two minutes, and translation is delayed by another four minutes. To simplify the mixed-effects model, this delay is excluded from the ODE system, and instead realized by linearly interpolating and shifting the measurements by six minutes. Subsequently, in the new shifted time grid, only times *t* = 0, 5, 10, … , 100 minutes are considered (*t* = 0, 5, … , 200 minutes for the two-step model). Since some of the fluorescence measurements are contaminated with background luminosity, the corresponding time series were shifted to start at zero. Any measurements that became negative as a result of this modification were truncated at zero. Cells with missing or too few measurements were omitted.

##### Approximation of an equivalent maturation half-time for proteins undergoing two-step maturation

For proteins with a single rate-limiting maturation step, the maturation half-time t_1/2_ and the maturation rate are related through 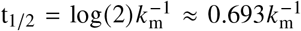. The relationship between maturation rates and the maturation half-time is less straightforward for proteins with two rate-limiting maturation steps, as the half-times of the individual steps only form a lower bound for the actual maturation half-time of the protein [Guerra et al., 2022]. To approximate this half-time analytically, we consider the solution to the initial value problem

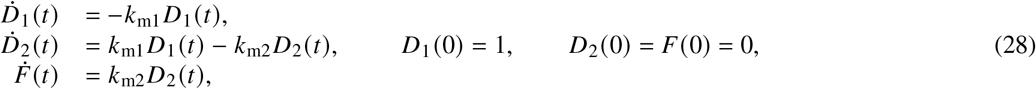

and find the time t_1/2_ at which *F* (t_1/2_) = 0.5. As we have previously shown [Guerra et al., 2022], independent inference of *k*_m1_ and *k*_m2_ is not possible when only measurements of *F* (*t*) are available, because *k*_m1_ and *k*_m2_ are practically non-identifiable. To overcome this problem, we assume that *k*_m_ := *k*_m1_ = *k*_m2_. Under this assumption, the solution the solution *F* (*t*) to (28) is

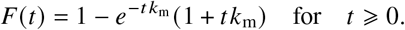

Therefore, t_1/2_ solves

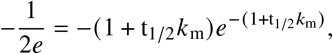

Implying

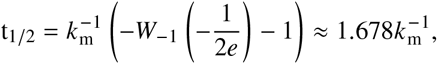

where *W*_−1_ : [−1/ *e*, 0) → ℝ denotes the real part of the the (− 1)-branch of the Lambert *W* function [Corless et al., 1996]. This relation provides a straightforward connection between the maturation half-time and the maturation rates of the two-step model.

##### Approximation of nonnegative maturation half-time distributions

Inference methods based on the classical GTS framework produce normal distribution estimates. When inferring rate distributions, this assumption of normality implies positive probability density at 0, which translates to positive probability density at infinite maturation half-times. Since such half-times are biologically implausible, prior to converting the rates, we approximate their normal distributions with Gamma distributions. To preserve the location of the mean and the variability, we chose to equate the modes and variances of both distributions. Let *μ* > 0 denote the mean (which coincides with the mode) and *σ* > 0 denote the standard deviation of the normal distribution to be approximated. For a Gamma distribution with shape *α* > 0 and rate *β* > 0, we need to solve

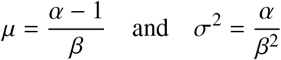

for *α, β*. Combining the equations for *μ* and *σ*^2^ yields a quadratic equation for *β* with positive solution

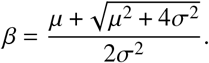

Substitution into the equation for *μ* yields

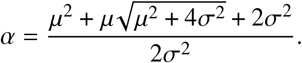

## Supplemental Information

### S1 Supplementary tables and figures

**Table T1:**
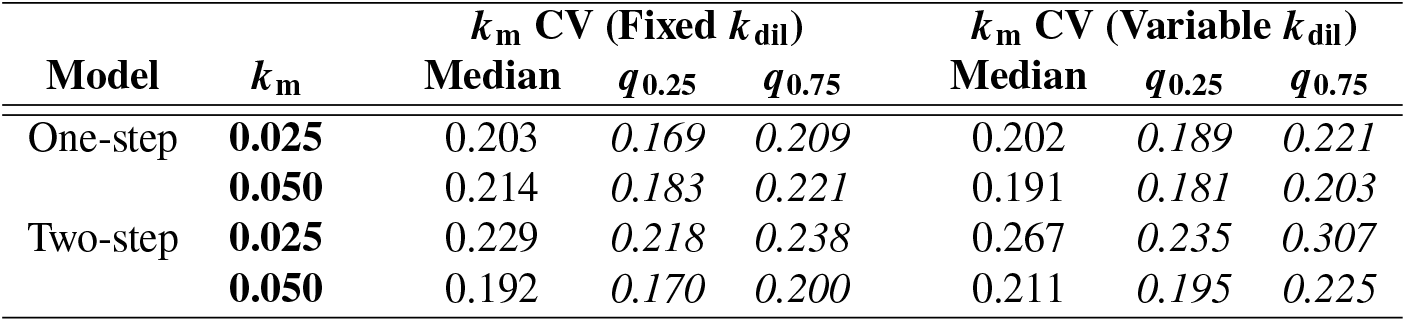
Simulation study on ignoring dilution rate variability in the FP maturation system. Synthetic data were generated under different scenarios (fixed versus variable *k*_dil_ and two different *k*_*m*_ means) and were used to infer the population distribution of *k*_*m*_, assuming multiplicative measurement noise with *τ* = 4%. The variability of the inferred distribution (summarized by the coefficient of variation) was then compared to the variability of the ground-truth (data-generating) distribution of *k*_*m*_. The nominal value *k*_dil_ = 0.004 and corresponding variability in terms of a CV of 0.05 were inferred from [Guerra et al., 2022]. The first column of the table indicates whether the one- or the two-step maturation model was used. The second column lists the *k*_m_ mean used to generate the data; each data-generating distribution of *k*_m_ had a CV of 0.25. Columns 3–5 and 6–8 summarize the CVs of the inferred *k*_*m*_ distributions for 10 independent repetitions of data generation and inference using medians and quartiles. The origin of the slight underestimation of the variability in *k*_m_ is discussed in the caption of Figure S2. Overall, there is no indication that treating the dilution rate as fixed when it is actually variable leads to inflated variability in the maturation rate estimates.

| Model | $k_m$ | $k_m$ CV (Fixed $k_{dil}$) | | | $k_m$ CV (Variable $k_{dil}$) | | |
| --- | --- | --- | --- | --- | --- | --- | --- |
| | | Median | $q_{0.25}$ | $q_{0.75}$ | Median | $q_{0.25}$ | $q_{0.75}$ |
| One-step | **0.025** | 0.203 | 0.169 | 0.209 | 0.202 | 0.189 | 0.221 |
|  | **0.050** | 0.214 | 0.183 | 0.221 | 0.191 | 0.181 | 0.203 |
| Two-step | **0.025** | 0.229 | 0.218 | 0.238 | 0.267 | 0.235 | 0.307 |
|  | **0.050** | 0.192 | 0.170 | 0.200 | 0.211 | 0.195 | 0.225 |

**Table T2:**
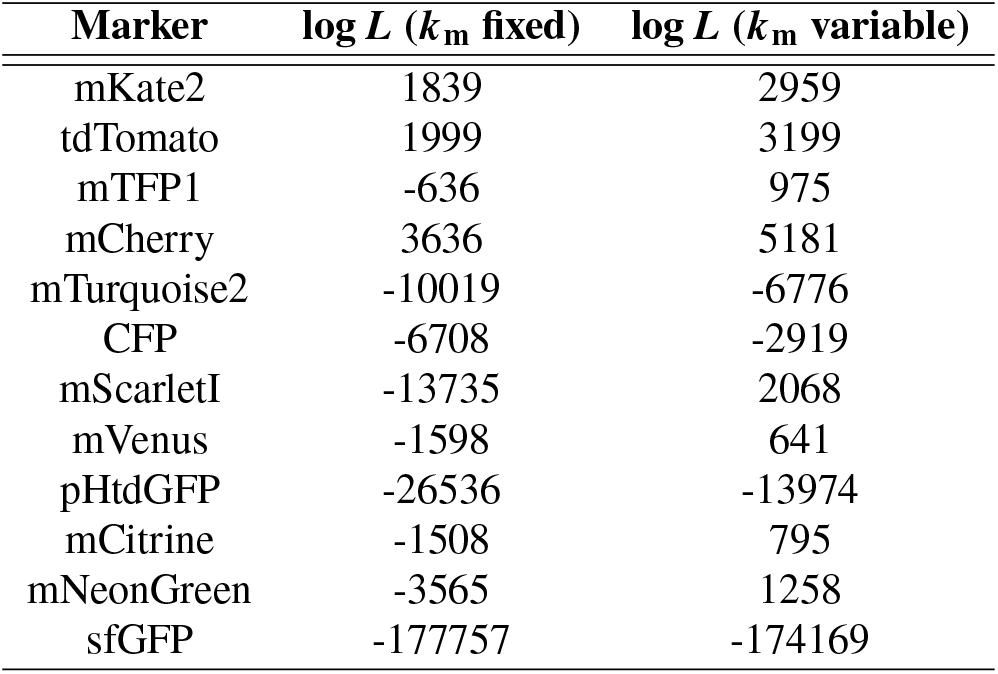
Marginal likelihood values of FP maturation models with fixed vs. variable maturation rate. Middle column: marginal log-likelihoods of a model with variable *k*_p_ and fixed *k*_m_ fixed. Right column: marginal log-likelihood values of a model with variable *k*_p_ and *k*_m_. Likelihoods are many orders of magnitude larger when *k*_m_ is considered variable across a population. Since all likelihood differences are substantial, the differences in degrees of freedom between the two models can be safely neglected; the model with variable maturation rate is considerably more likely given the experimental data.

| Marker | $\log L$ ($k_m$ fixed) | $\log L$ ($k_m$ variable) |
| --- | --- | --- |
| mKate2 | 1839 | 2959 |
| tdTomato | 1999 | 3199 |
| mTFP1 | -636 | 975 |
| mCherry | 3636 | 5181 |
| mTurquoise2 | -10019 | -6776 |
| CFP | -6708 | -2919 |
| mScarletI | -13735 | 2068 |
| mVenus | -1598 | 641 |
| pHtdGFP | -26536 | -13974 |
| mCitrine | -1508 | 795 |
| mNeonGreen | -3565 | 1258 |
| sfGFP | -177757 | -174169 |

**Table T3:** Summary of inferred maturation half time distributions and comparison with previously published results. Second column: maturation half-times (in minutes) estimated by [Guerra et al., 2022] for each fluorescent protein. Third column: mean half-times obtained from the GMGTS method. Remaining columns: summaries of the maturation half-time distributions by their 25% and 75% quartiles, as well as their relative quartile range (RQR). The latter is defined as the interquartile range divided by the mean and multiplied by 0.75 (making it comparable to the coefficient of variation for normal distributions). Faster-maturing proteins tend to have higher relative variability in terms of RQR.

| Marker | $t_{50}$ [Guerra et al., 2022] | Mean $t_{1/2}$ (GMGTS) | $q_{0.25}$ | $q_{0.75}$ | RQR |
| --- | --- | --- | --- | --- | --- |
| mKate2 | 111.9 | 104.2 | 93.1 | 113.6 | 0.148 |
| tdTomato | 104.9 | 89.5 | 76.4 | 100.0 | 0.198 |
| mTFP1 | 79.9 | 76.5 | 68.2 | 83.6 | 0.151 |
| mCherry | 49.4 | 44.0 | 32.0 | 51.7 | 0.336 |
| mTurquoise2 | 46.6 | 41.8 | 34.1 | 47.5 | 0.239 |
| mScarletI | 28.9 | 27.0 | 22.1 | 30.6 | 0.237 |
| CFP | 8.3 | 11.6 | 7.3 | 13.7 | 0.418 |
| mVenus | 13.9 | 10.5 | 7.9 | 12.3 | 0.314 |
| pHtdGFP | 12.4 | 10.1 | 6.8 | 12 | 0.383 |
| mCitrine | 12.0 | 10.1 | 7.5 | 11.8 | 0.320 |
| mNeonGreen | 12.0 | 9.2 | 6.4 | 10.9 | 0.362 |
| sfGFP | 6.9 | 7.0 | 4.3 | 8.4 | 0.427 |

**Figure S1:**
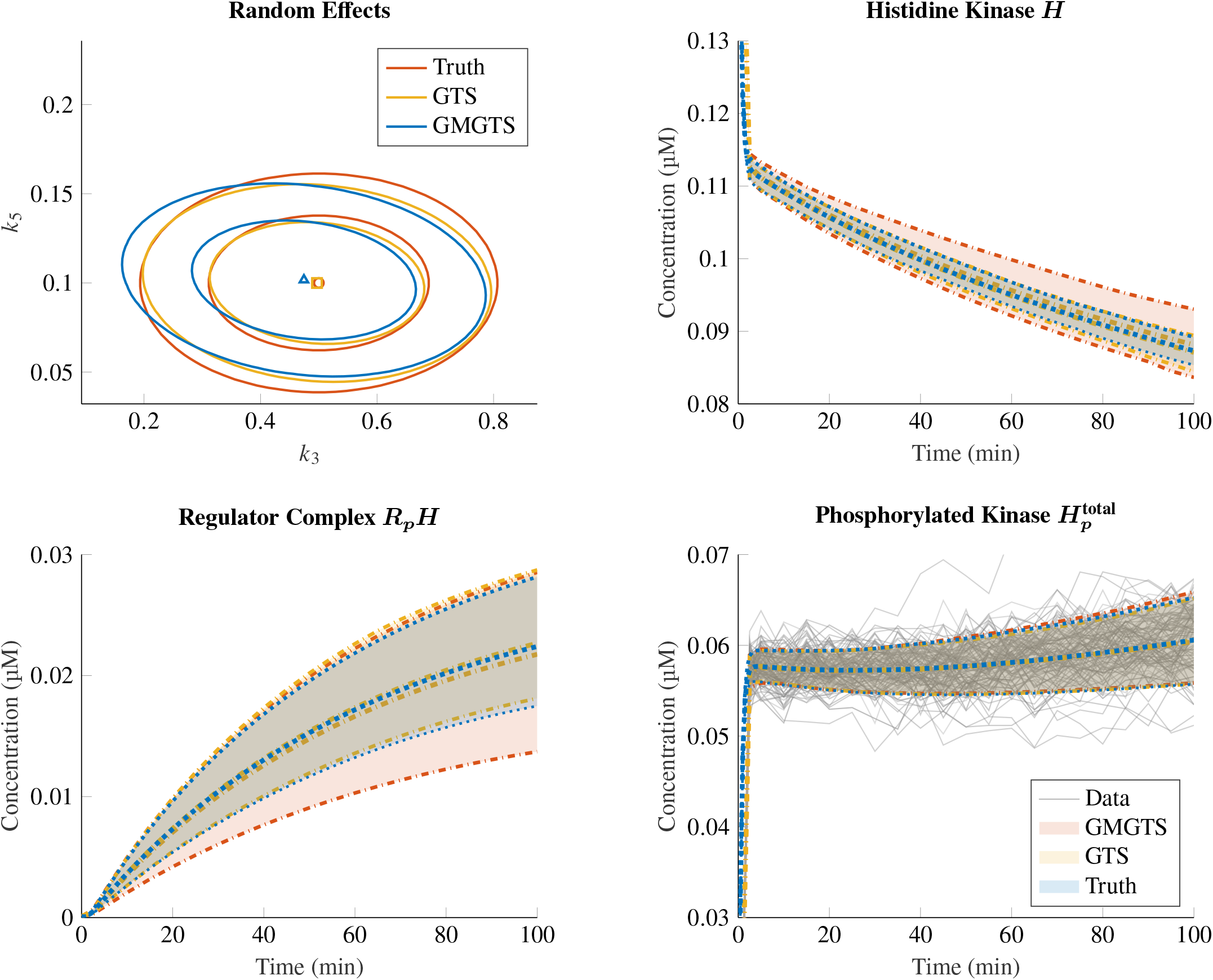
Bifunctional two-component system with partial observation: examples of inference and prediction results. *Top left panel:* example of a (marginal) random effects distribution inferred by the GMGTS and GTS methods, compared to the true data-generating distribution. Distributions are summarized by their mean and 68% and 95% contour levels. *Remaining panels:* predicted distributions of some unobserved and observed states (unphosphorylated histidine kinase *H*, phosphorylated response regulator-kinase complex *R*_*p*_*H*, and total phosphorylated histidine kinase 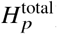), summarized by their mean and 90% confidence intervals of the state distributions. The simulated measurement data for 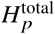 are also included in the bottom-right panel. The (multiplicative) noise level was set at *τ* = 3% and the data were simulated at *T* = 22 time points. The attained Wasserstein distances of the inferred random effects distributions from the ground-truth are 0.074 versus 0.075 for GMGTS and GTS respectively.

**Figure S2:**
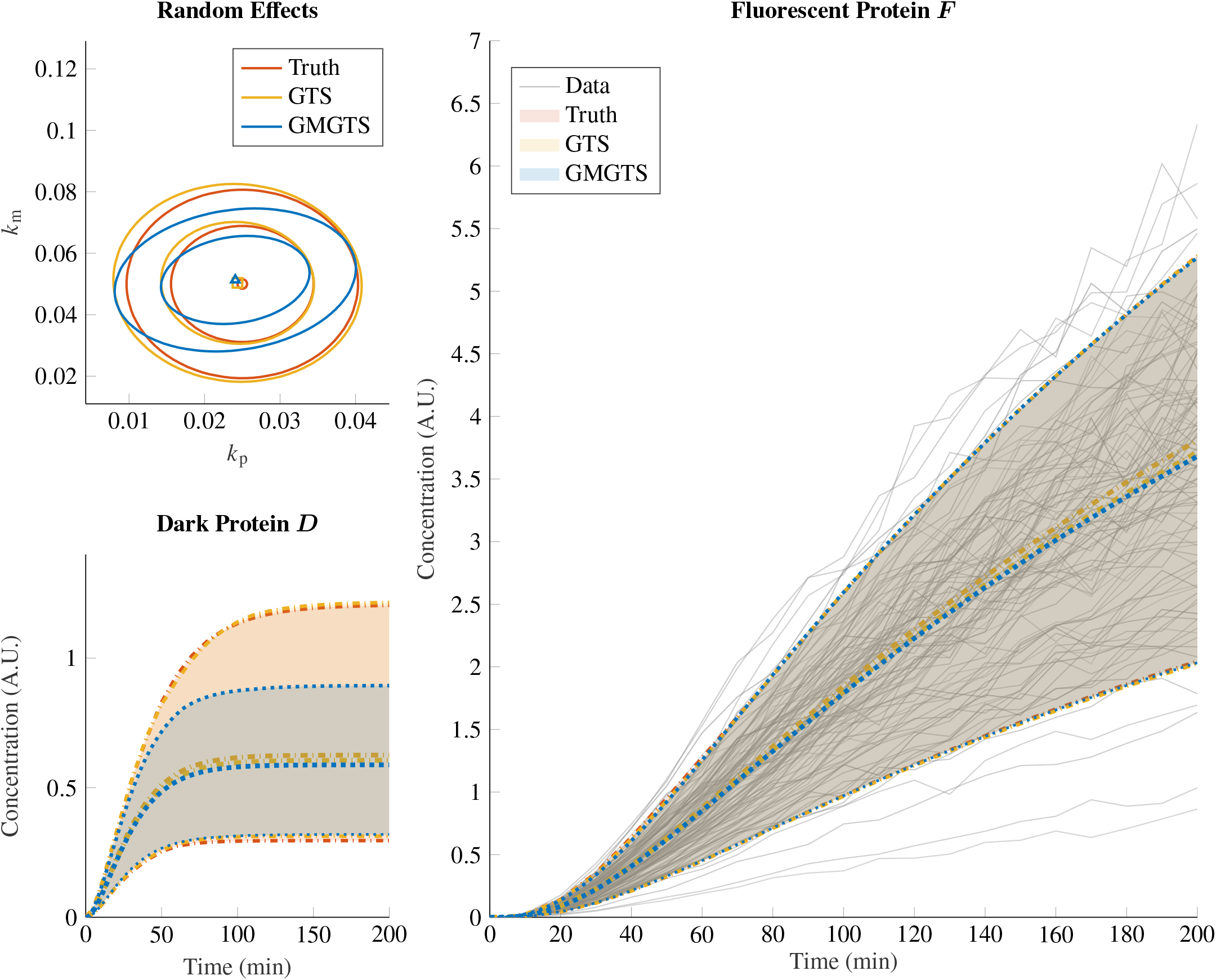
Simulation study of FP maturation dynamics: illustration of inference and prediction results. *Top left panel:* random effects distributions inferred by the GMGTS and GTS methods compared to the true data-generating distribution, summarized by their mean and 68% and 95% contour levels. *Remaining panels:* predicted state distributions (immature and mature FP), summarized by their mean and variability (90% confidence intervals of the state distributions). The simulated measurement data for *F* (mature FP) are also included in the right panel. The (multiplicative) noise level was set at *τ* = 3% and the data were simulated at *T* = 22 time points. The attained Wasserstein distances of the inferred random effects distributions from the ground-truth are 0.0022 versus 0.0010 for GMGTS and GTS respectively. The downward bias in the variability of *k*_*m*_ arises mainly from the relatively long sampling interval of the simulated data (set at 5 min to match that of the experimental data). Sparse sampling increases the uncertainty of the first 10-15 minutes of the smoothed single-cell time series, and this increased uncertainty makes it harder for the individual maturation rate estimates to move away from their common initial value.

**Figure S3:**
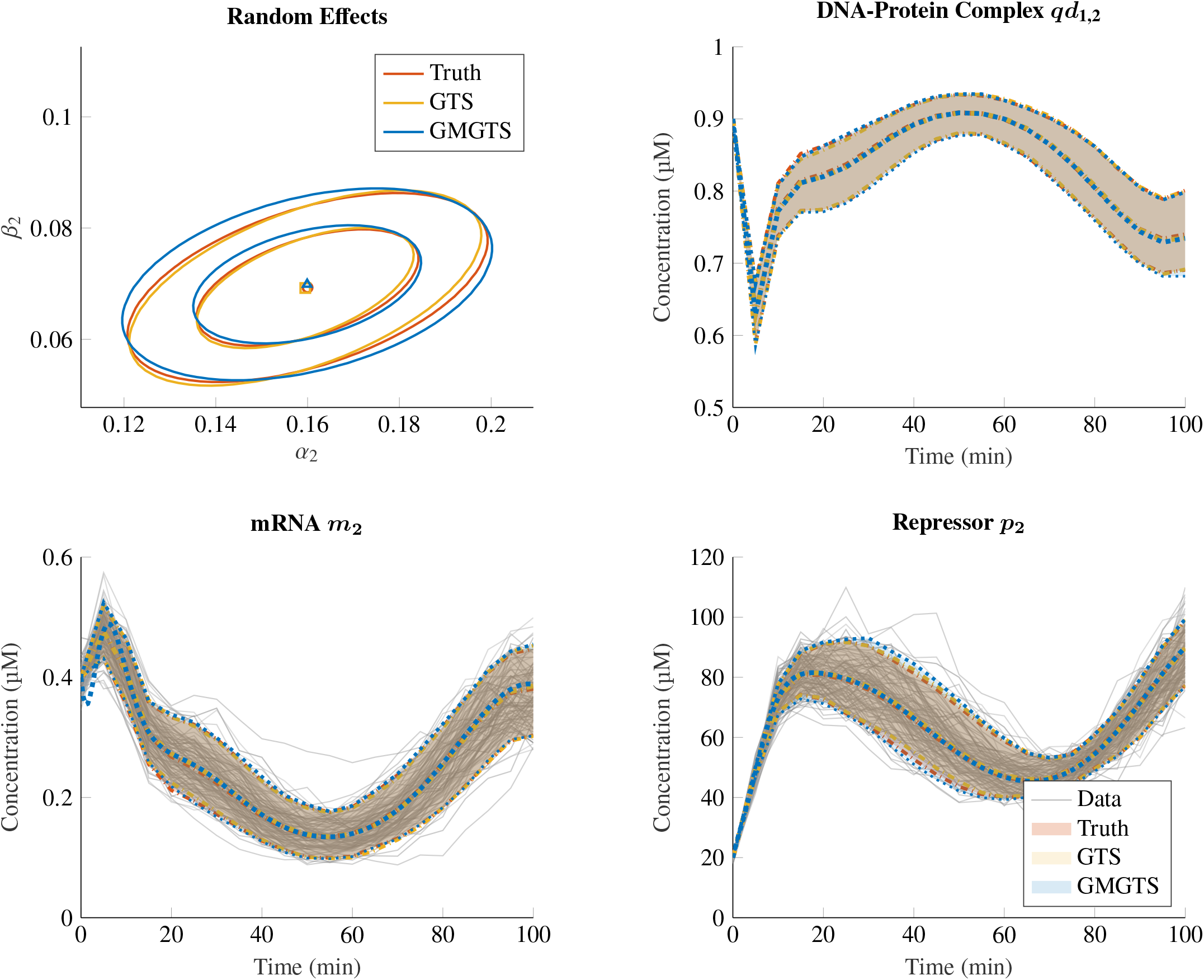
Simulation study of the repressilator: illustration of inference and predictions results. *Top left panel:* selected random effects distributions inferred by the GMGTS and GTS methods compared to the true data-generating distribution, summarized by their mean and 68% and 95% contour levels. *Remaining panels:* predicted state distributions (DNA-protein complex *qd*_1,2_, mRNA *m*_1_, and repressor *p*_1_ concentrations) summarized by their mean and variability (90% confidence intervals of the state distributions). The selected states directly influence the concentration of the first repressor in the system (*p*_1_), and similar results are obtained for the remaining ones. The simulated measurement data for *m*_1_ and *p*_1_ are also included in the bottom panels, where the (multiplicative) noise level was set at *τ* = 5% and the data were simulated at *T* = 21 time points.

**Figure S4:**
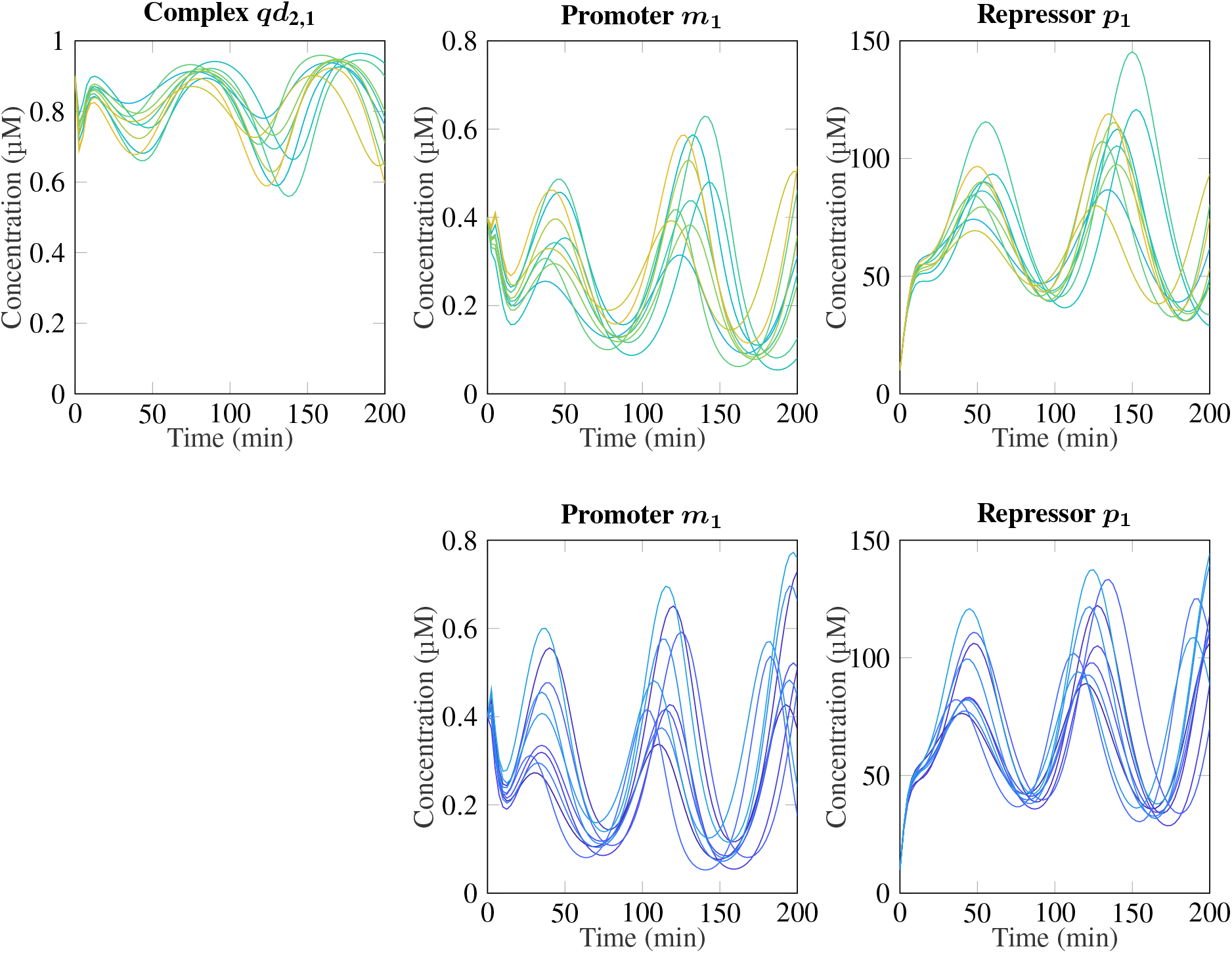
Repressilator trajectories obtained from the expanded and original system equations. The states that directly influence repressor *p*_1_ are plotted for a sample of ten parameter sets *α*_*n*_, *β*_*n*_ (*n* = 1, 2, 3) drawn from the random effects distribution described in Results. Similar trajectories are obtained for the remaining states. *Top panels:* Trajectories simulated from the repressilator system with explicitly modeled protein-DNA interactions (18). *Bottom panels:* Trajectories simulated from the condensed repressilator system obtained from quasi-steady-state approximations and simplifying assumptions. (3).

**Figure S5:**
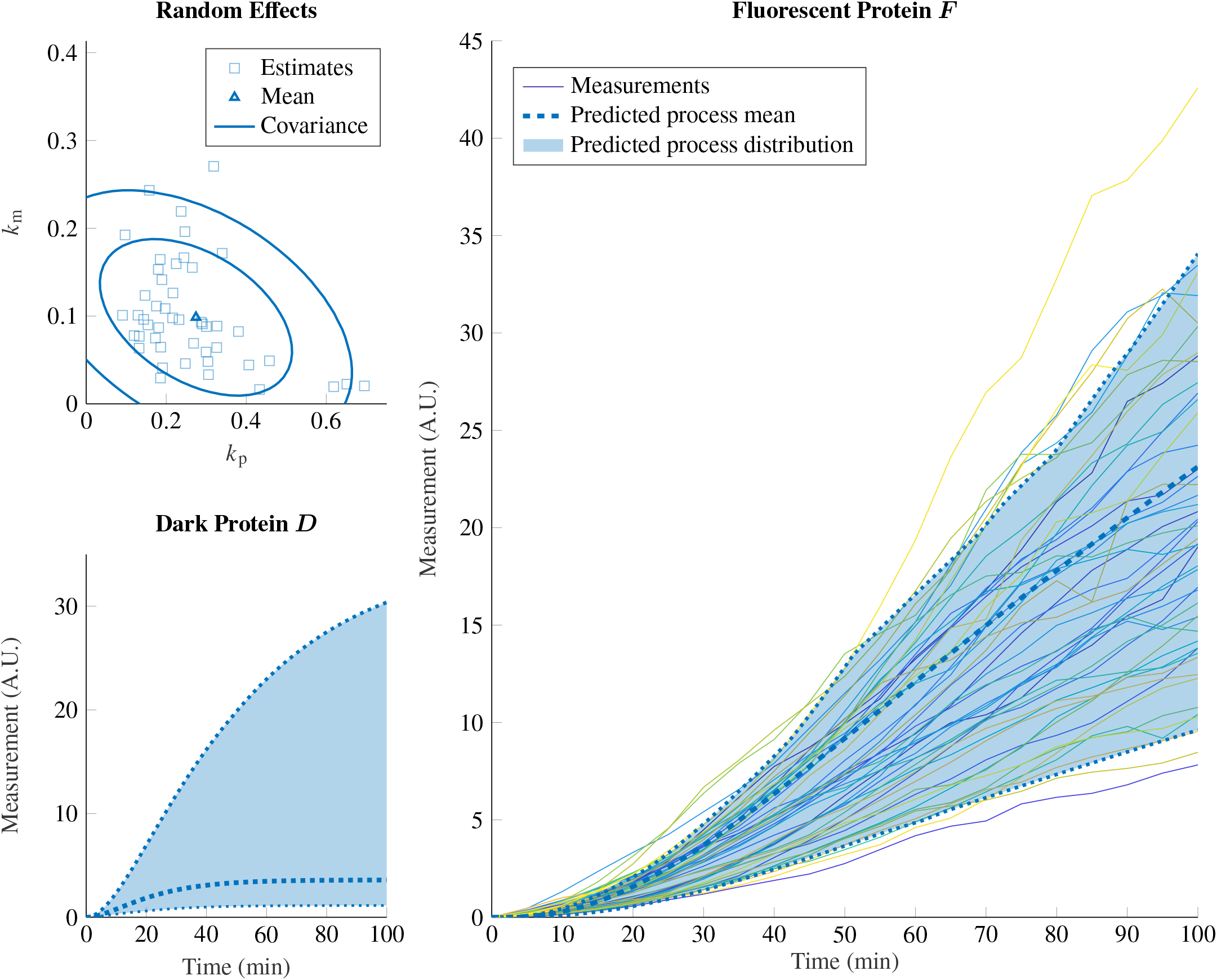
sfGFP: parameter inference and state predictions. *Top left panel:* individual estimates of *k*_p_ and *k*_m_ produced by the first stage of GMGTS (squares) and the associated random effects distribution obtained from the second stage, summarized by its mean and 68% and 95% contour levels. *Remaining panels:* Predicted distributions of immature and mature FP, summarized by their mean and variability (90% confidence intervals of the state distributions). The (preprocessed) experimental data are also included in the right panel.

**Figure S6:**
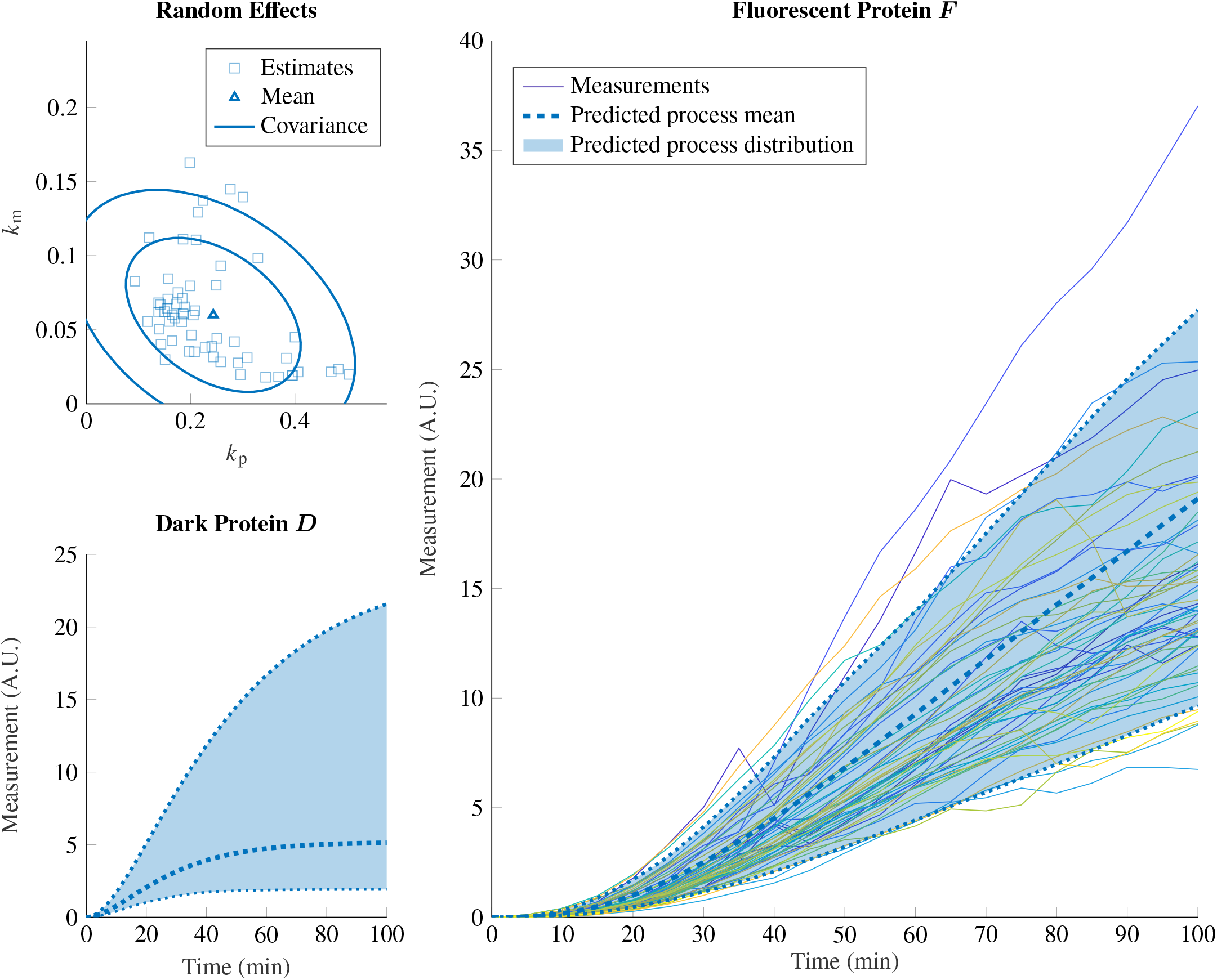
CFP: parameter inference and state predictions. *Top left panel:* individual estimates of *k*_p_ and *k*_m_ produced by the first stage of GMGTS (squares) and the associated random effects distribution obtained from the second stage, summarized by its mean and 68% and 95% contour levels. *Remaining panels:* Predicted distributions of immature and mature FP, summarized by their mean and variability (90% confidence intervals of the state distributions). The (preprocessed) experimental data are also included in the right panel.

**Figure S7:**
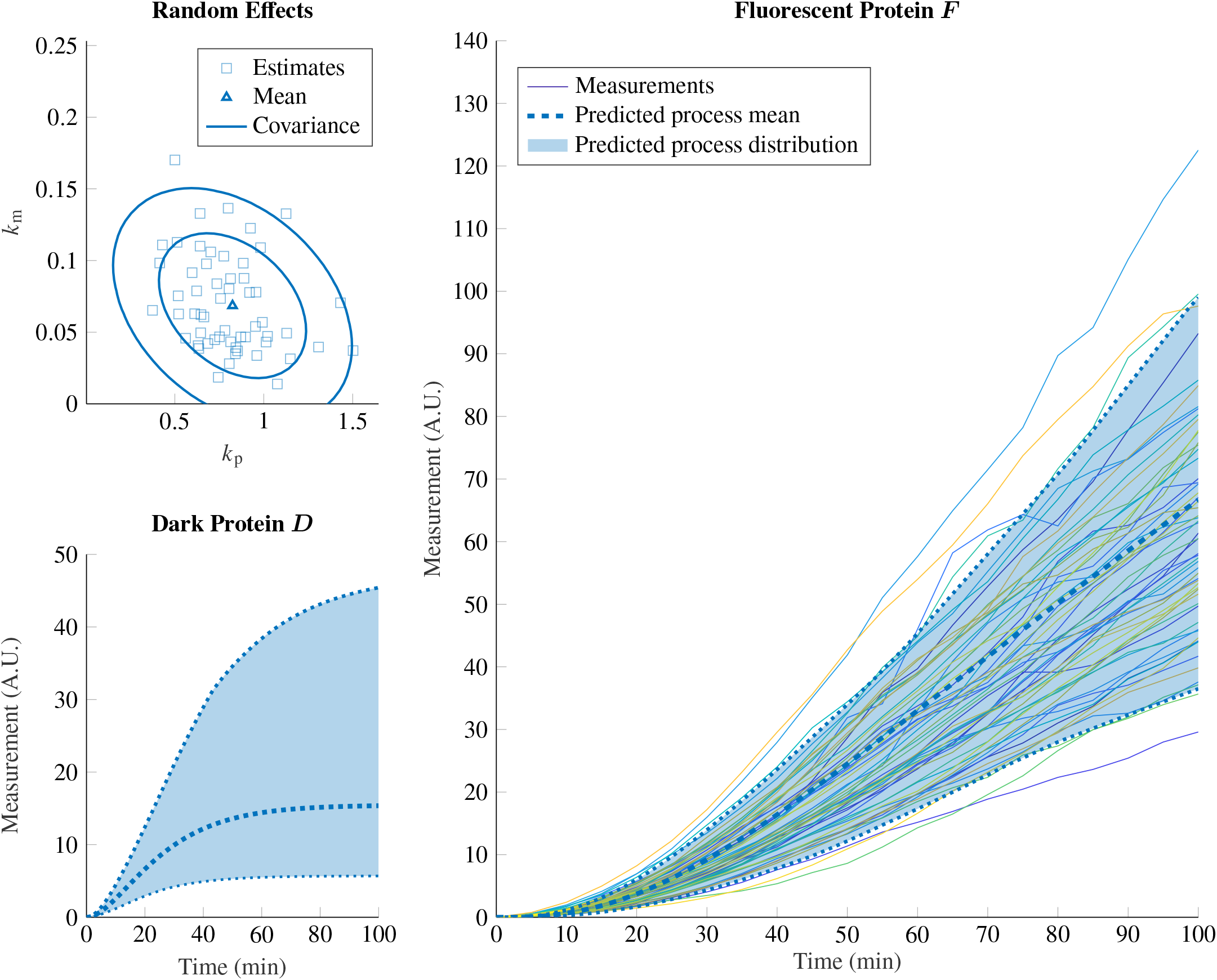
pHtdGFP: parameter inference and state predictions. *Top left panel:* individual estimates of *k*_p_ and *k*_m_ produced by the first stage of GMGTS (squares) and the associated random effects distribution obtained from the second stage, summarized by its mean and 68% and 95% contour levels. *Remaining panels:* Predicted distributions of immature and mature FP, summarized by their mean and variability (90% confidence intervals of the state distributions). The (preprocessed) experimental data are also included in the right panel.

**Figure S8:**
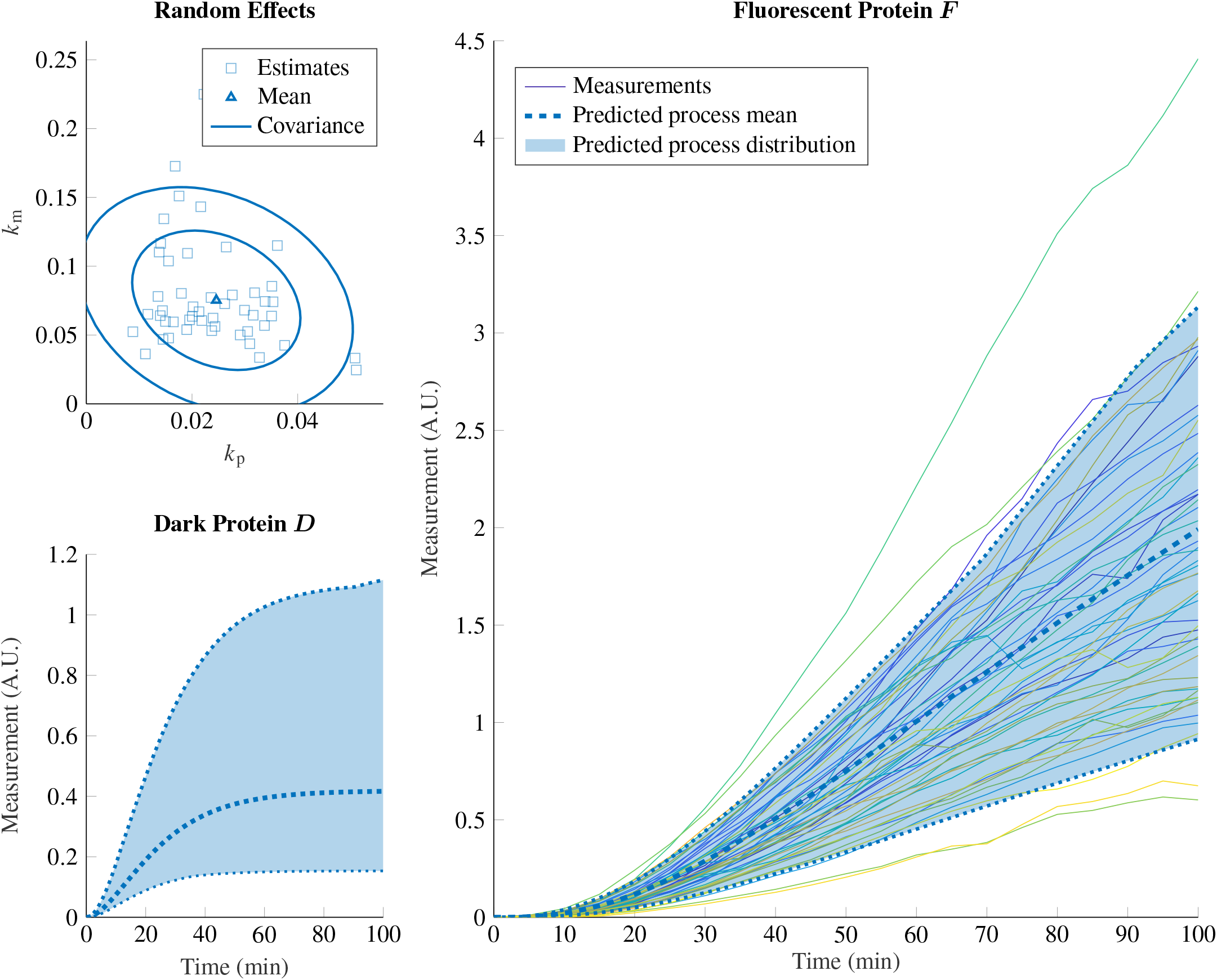
mNeonGreen: parameter inference and state predictions. *Top left panel:* individual estimates of *k*_p_ and *k*_m_ produced by the first stage of GMGTS (squares) and the associated random effects distribution obtained from the second stage, summarized by its mean and 68% and 95% contour levels. *Remaining panels:* Predicted distributions of immature and mature FP, summarized by their mean and variability (90% confidence intervals of the state distributions). The (preprocessed) experimental data are also included in the right panel.

**Figure S9:**
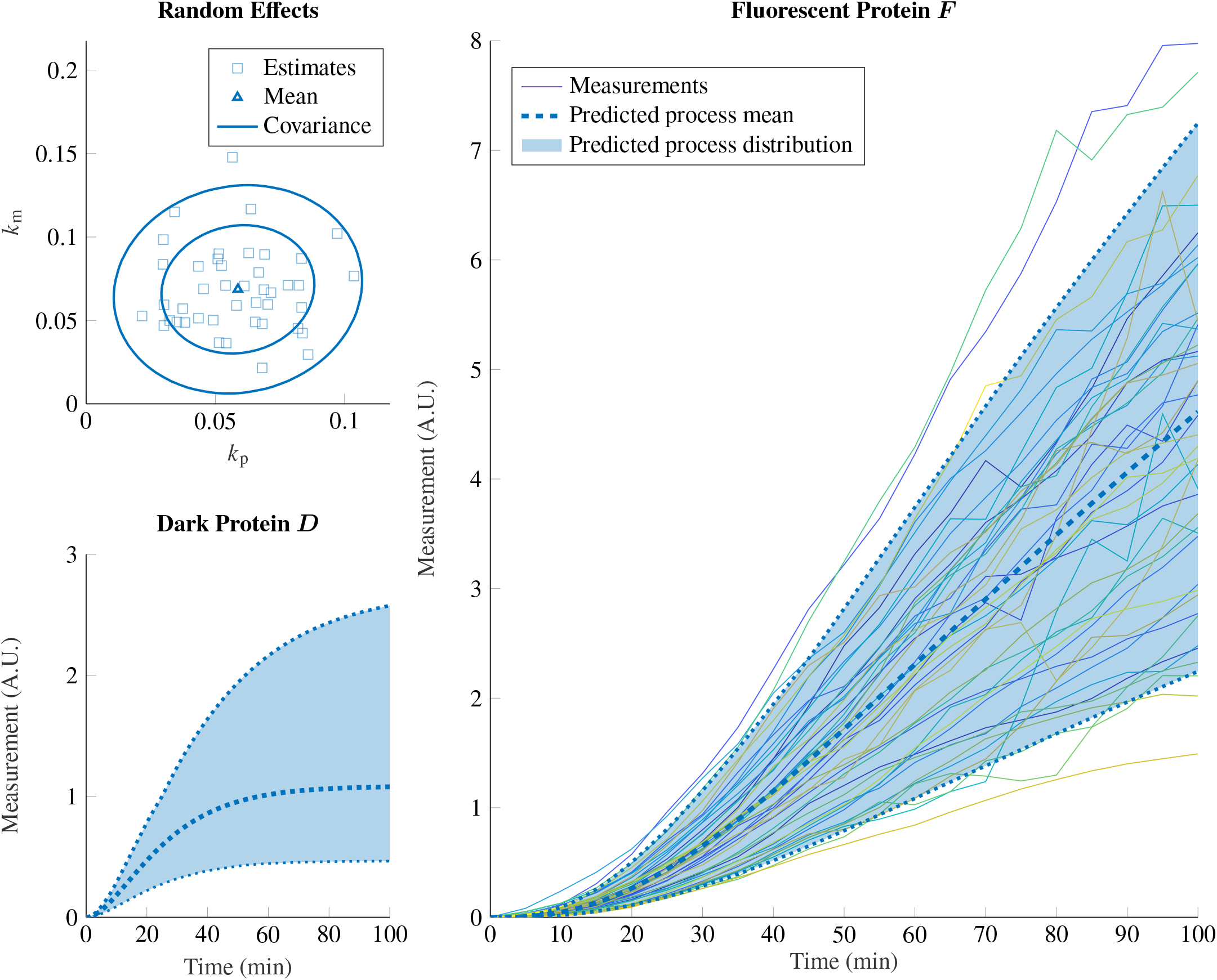
mCitrine: parameter inference and state predictions. *Top left panel:* individual estimates of *k*_p_ and *k*_m_ produced by the first stage of GMGTS (squares) and the associated random effects distribution obtained from the second stage, summarized by its mean and 68% and 95% contour levels. *Remaining panels:* Predicted distributions of immature and mature FP, summarized by their mean and variability (90% confidence intervals of the state distributions). The (preprocessed) experimental data are also included in the right panel.

**Figure S10:**
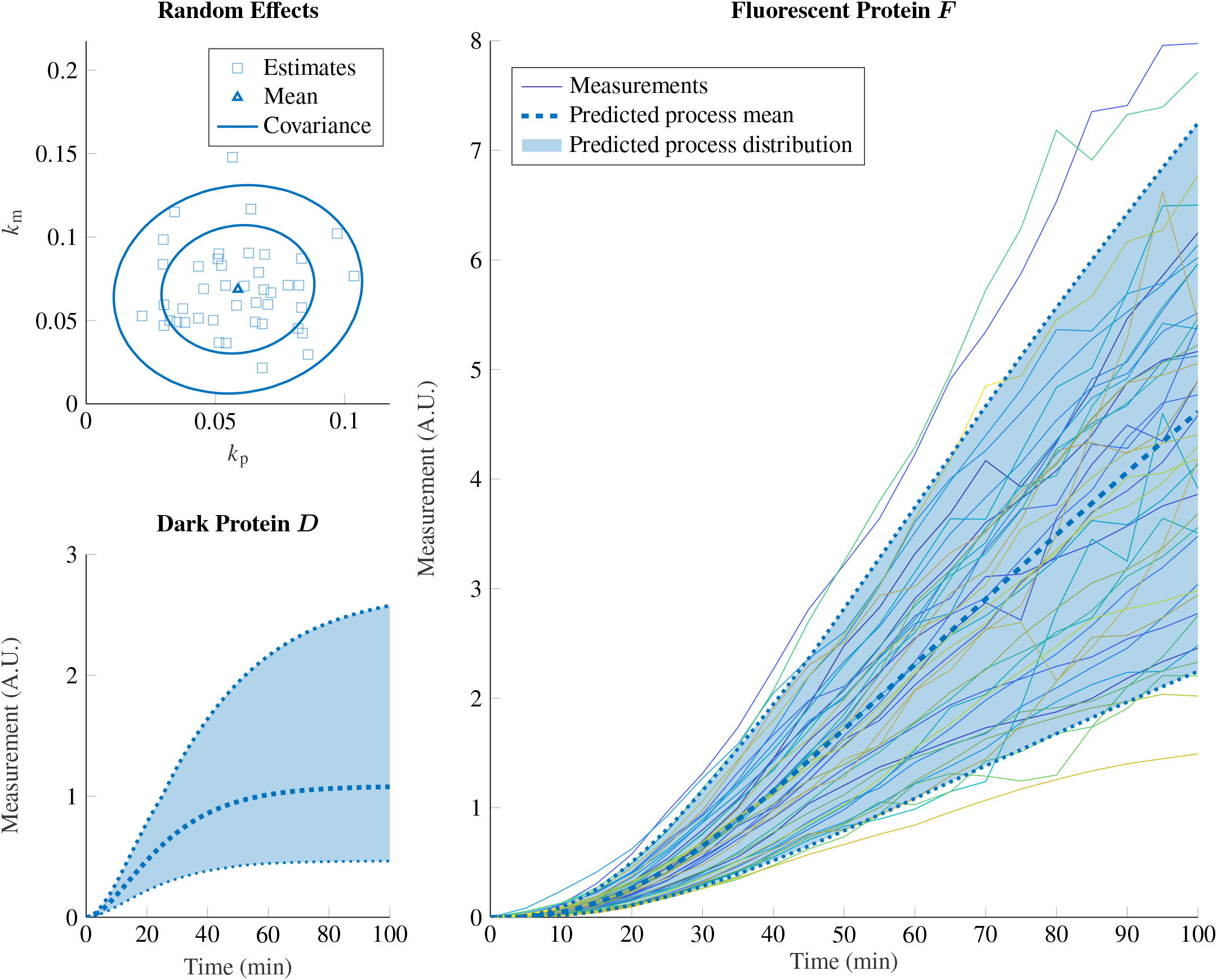
mVenus: parameter inference and state predictions. *Top left panel:* individual estimates of *k*_p_ and *k*_m_ produced by the first stage of GMGTS (squares) and the associated random effects distribution obtained from the second stage, summarized by its mean and 68% and 95% contour levels. *Remaining panels:* Predicted distributions of immature and mature FP, summarized by their mean and variability (90% confidence intervals of the state distributions). The (preprocessed) experimental data are also included in the right panel.

**Figure S11:**
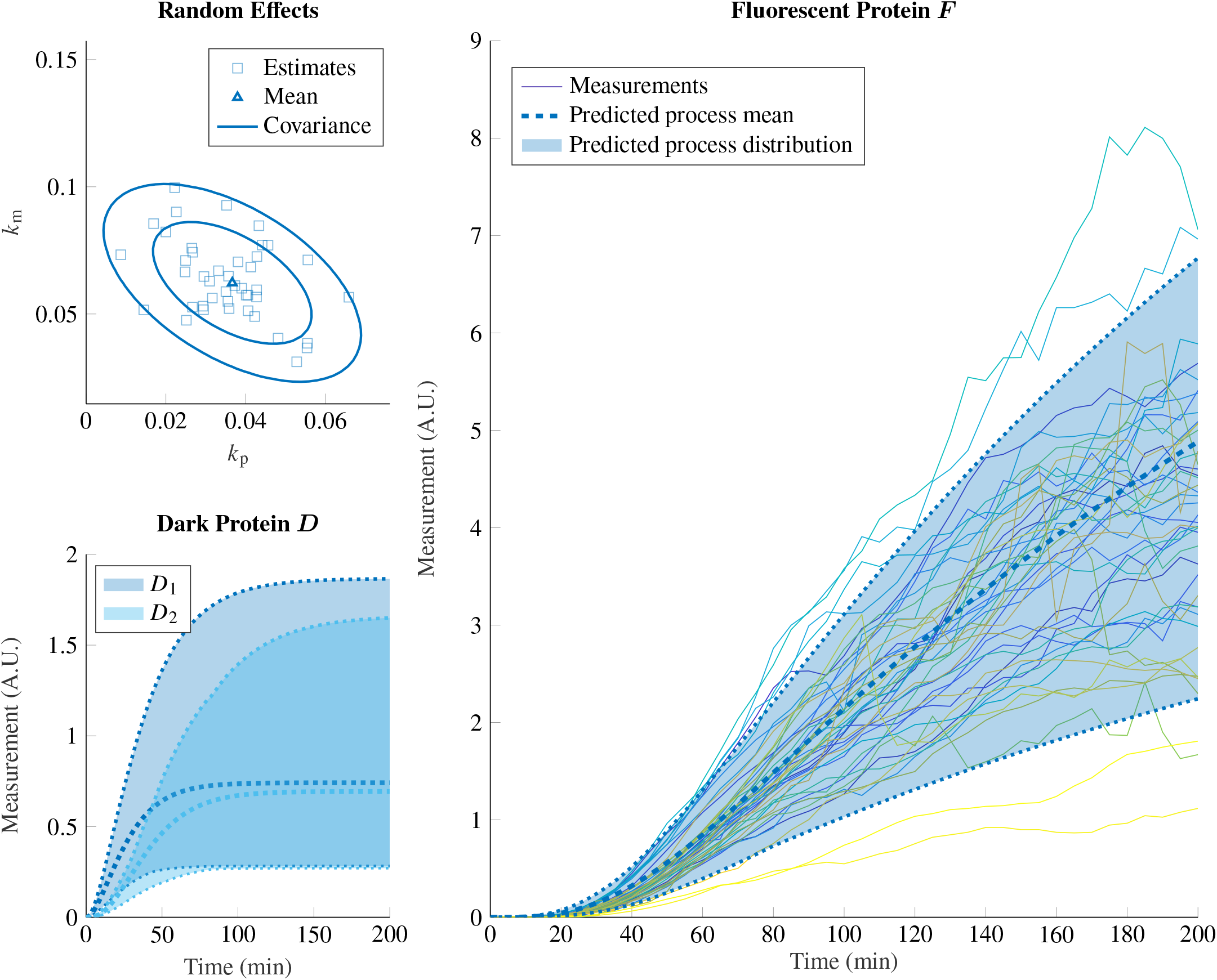
mScarlet-I: parameter inference and state predictions. *Top left panel:* individual estimates of *k*_p_ and *k*_m_ produced by the first stage of GMGTS (squares) and the associated random effects distribution obtained from the second stage, summarized by its mean and 68% and 95% contour levels. *Remaining panels:* Predicted distributions of immature and mature FP, summarized by their mean and variability (90% confidence intervals of the state distributions). The (preprocessed) experimental data are also included in the right panel.

**Figure S12:**
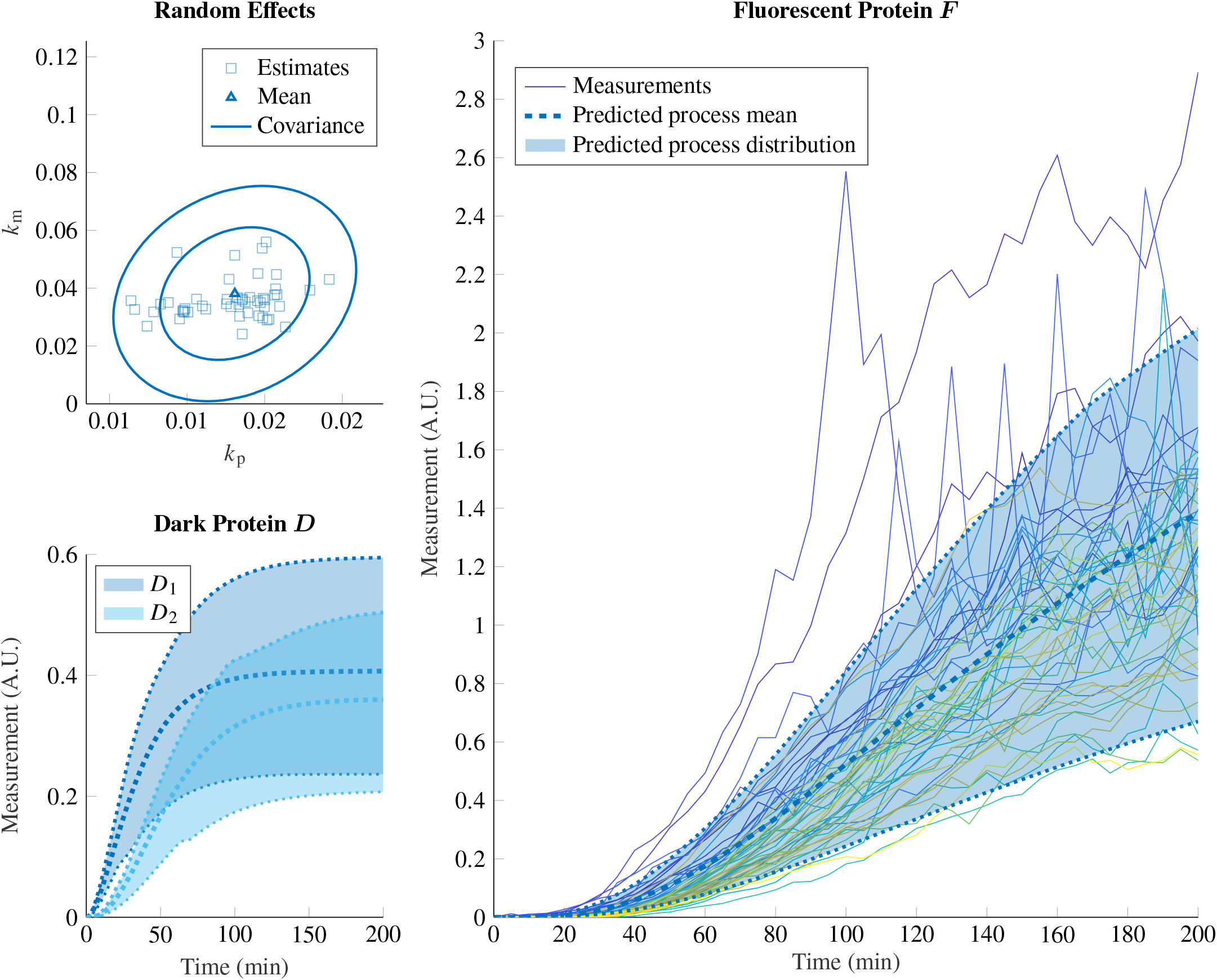
mCherry: parameter inference and state predictions. *Top left panel:* individual estimates of *k*_p_ and *k*_m_ produced by the first stage of GMGTS (squares) and the associated random effects distribution obtained from the second stage, summarized by its mean and 68% and 95% contour levels. *Remaining panels:* Predicted distributions of immature and mature FP, summarized by their mean and variability (90% confidence intervals of the state distributions). The (preprocessed) experimental data are also included in the right panel.

**Figure S13:**
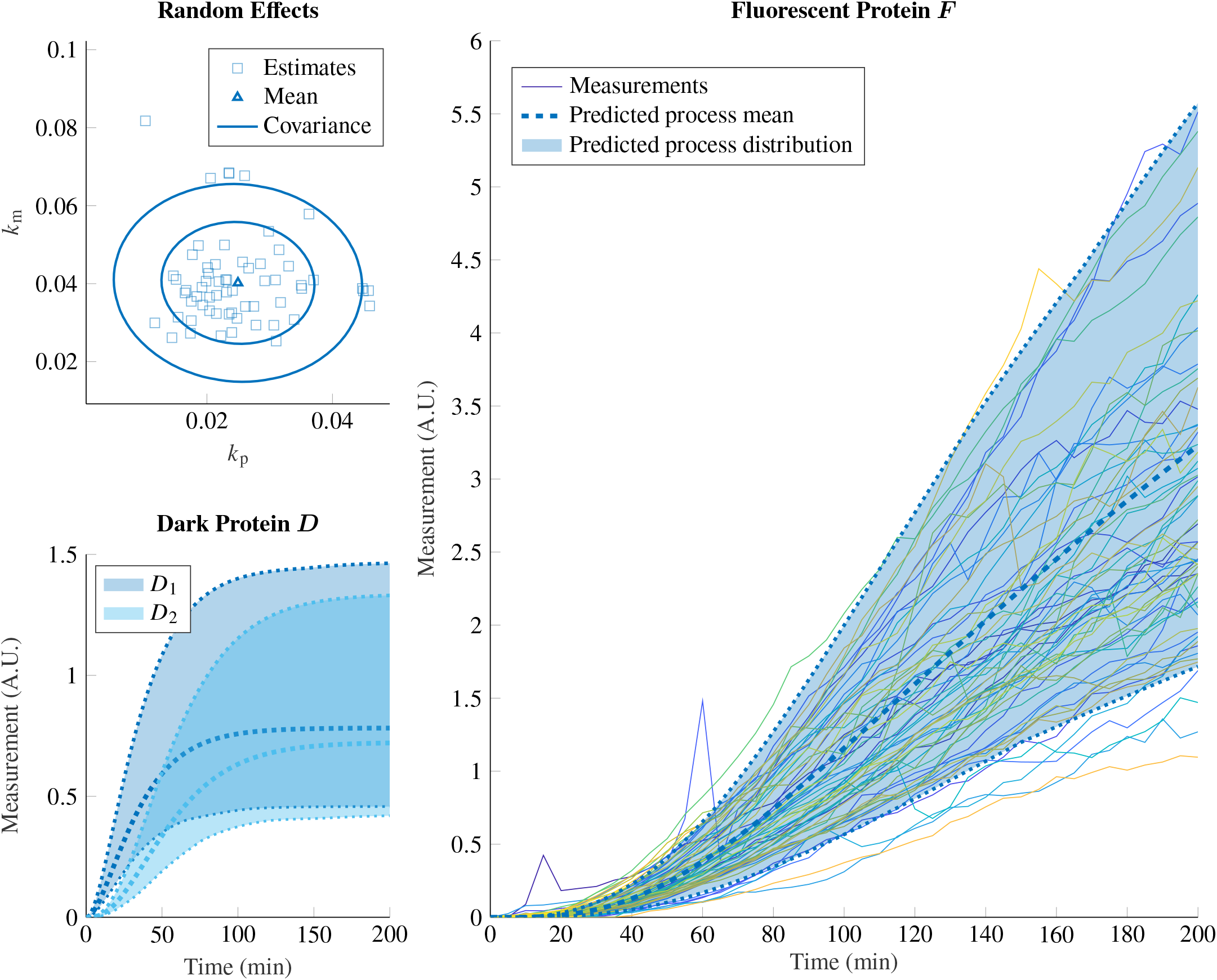
mTurquoise2: parameter inference and state predictions. *Top left panel:* individual estimates of *k*_p_ and *k*_m_ produced by the first stage of GMGTS (squares) and the associated random effects distribution obtained from the second stage, summarized by its mean and 68% and 95% contour levels. *Remaining panels:* Predicted distributions of immature and mature FP, summarized by their mean and variability (90% confidence intervals of the state distributions). The (preprocessed) experimental data are also included in the right panel.

**Figure S14:**
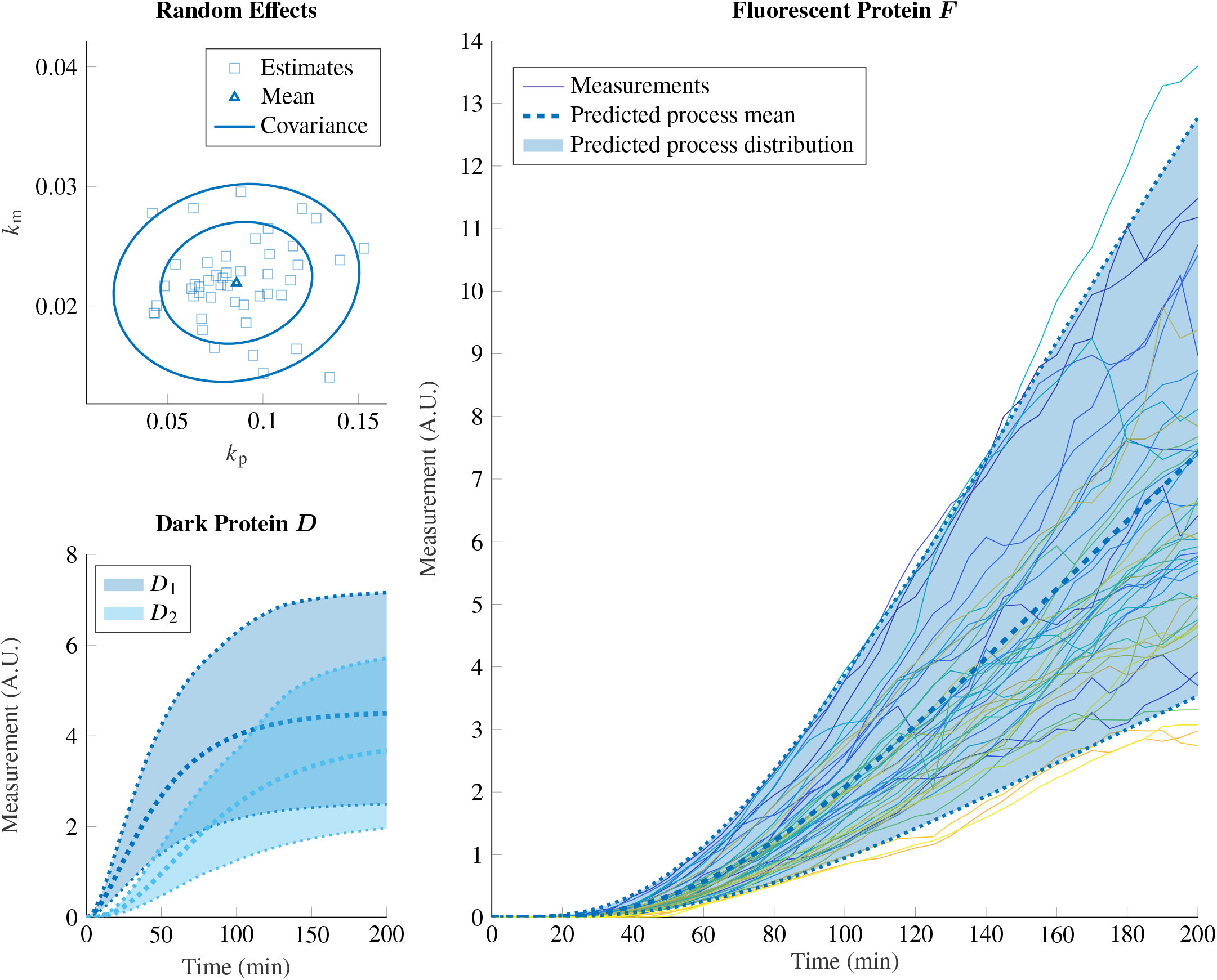
mTFP1: parameter inference and state predictions. *Top left panel:* individual estimates of *k*_p_ and *k*_m_ produced by the first stage of GMGTS (squares) and the associated random effects distribution obtained from the second stage, summarized by its mean and 68% and 95% contour levels. *Remaining panels:* Predicted distributions of immature and mature FP, summarized by their mean and variability (90% confidence intervals of the state distributions). The (preprocessed) experimental data are also included in the right panel.

**Figure S15:**
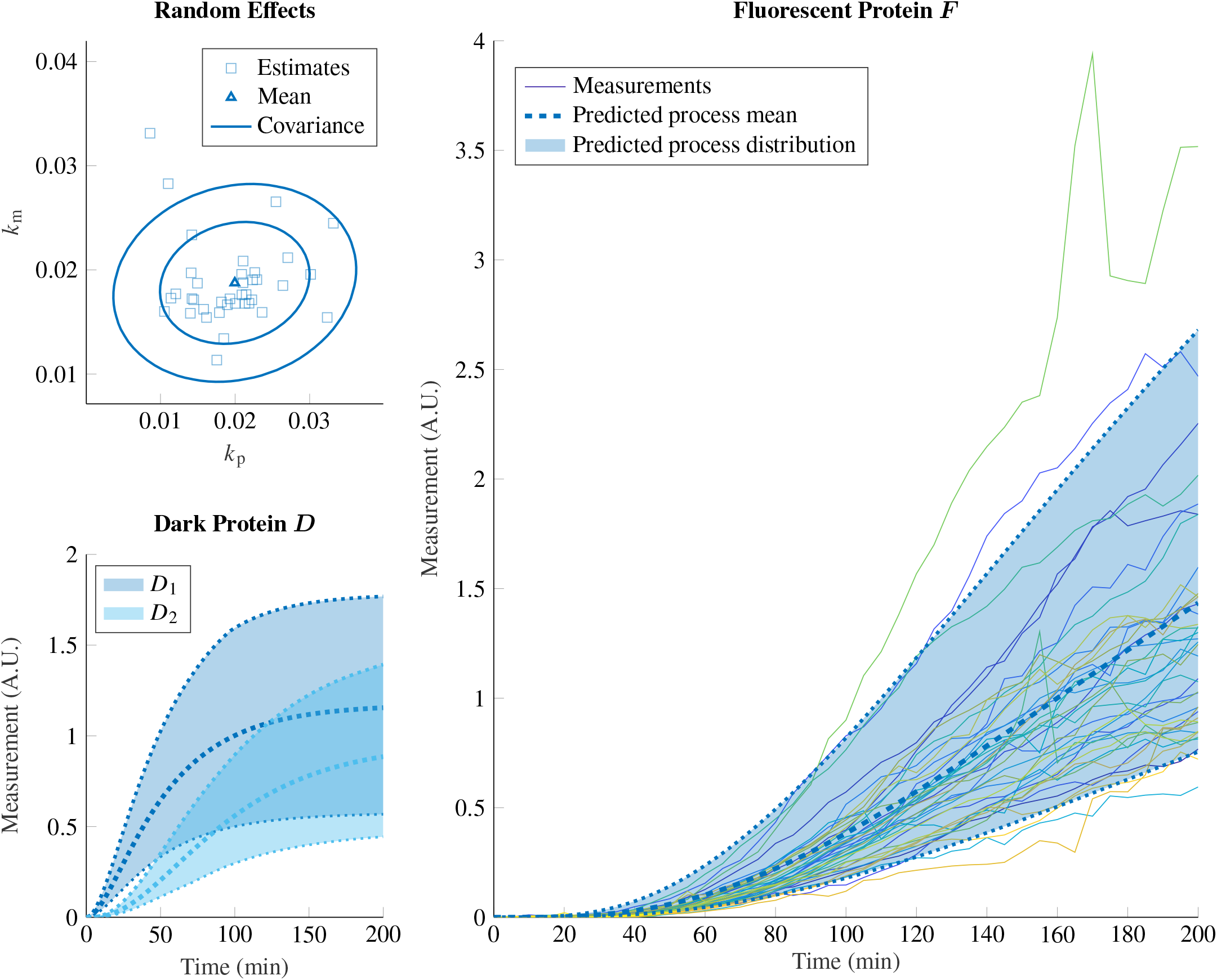
tdTomato: parameter inference and state predictions. *Top left panel:* individual estimates of *k*_p_ and *k*_m_ produced by the first stage of GMGTS (squares) and the associated random effects distribution obtained from the second stage, summarized by its mean and 68% and 95% contour levels. *Remaining panels:* Predicted distributions of immature and mature FP, summarized by their mean and variability (90% confidence intervals of the state distributions). The (preprocessed) experimental data are also included in the right panel.

**Figure S16:**
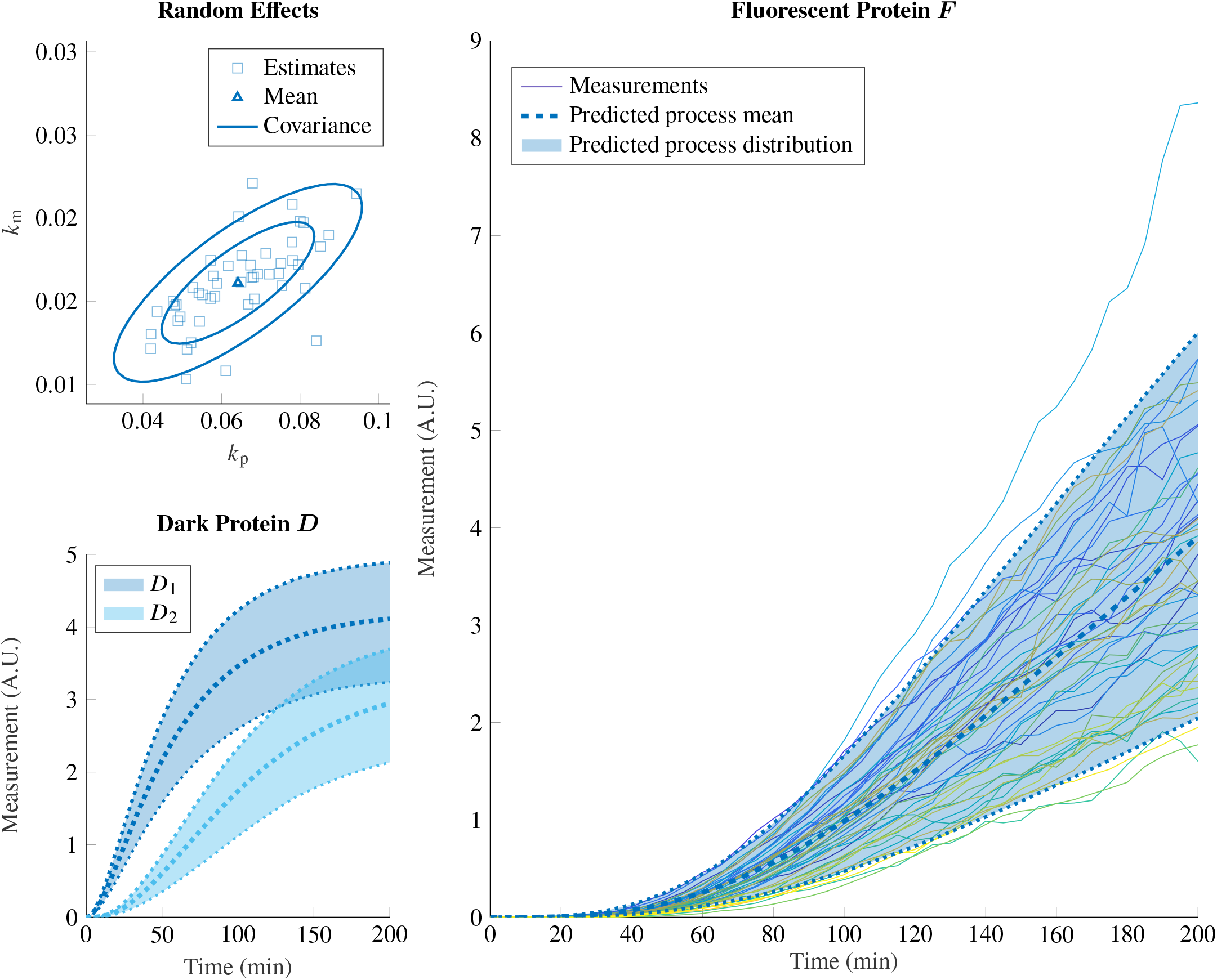
mKate2: parameter inference and state predictions. *Top left panel:* individual estimates of *k*_p_ and *k*_m_ produced by the first stage of GMGTS (squares) and the associated random effects distribution obtained from the second stage, summarized by its mean and 68% and 95% contour levels. *Remaining panels:* Predicted distributions of immature and mature FP, summarized by their mean and variability (90% confidence intervals of the state distributions). The (preprocessed) experimental data are also included in the right panel.

**Figure S17:**
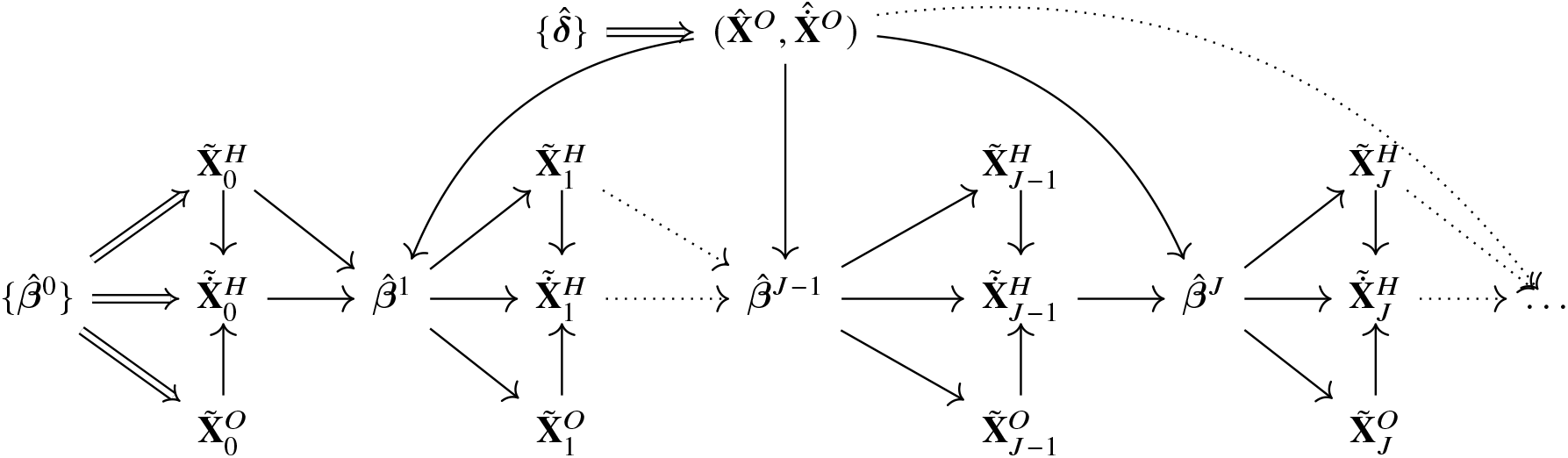
Dependence hierarchy in the iterative optimization scheme for partially observed systems. Uncertainty enters the scheme through the initial parameter estimate 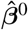 and the spline coefficient estimates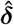. These estimates determine the initial hidden state estimates (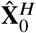 and 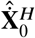) and the smoothed measurements (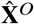and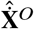), respectively, as well as their associated uncertainties. As soon as we obtain 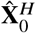 and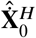 , we use these values together with the smoothed measurements (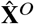 and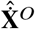 ) to obtain a new parameter estimate 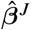 (*J* ≥ 1) via gradient matching. We also calculate the uncertainty of 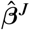 from the covariance of hidden and observed states and their derivatives at step *J* − 1. The new 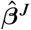 estimate gives rise to new hidden state estimates (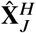 and 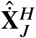) with their corresponding uncertainty, and the cycle repeats.

### S2 Notation

This section briefly lists some national conventions used in the supplemental mathematical derivations. Numerous references are also made to the main paper; to be able to clearly discern referenced sections, equations, and remarks, all items appearing in this document are numbered starting with ‘S’.

- Vectors and matrices are written in boldface. They can be defined in terms of scalars or smaller vectors/matrices by enclosing these quantities in square brackets. For matrices 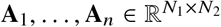 (where possibly *N*_1_ = *N*_2_ = 1), we define a “stacked” matrix using the shorthand notation of a comma-separated bracketed list; that is,

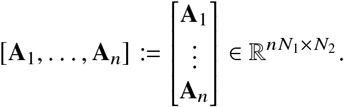 In particular, we let [*a*_1_, … , *a*_*n*_] denote an *n*-dimensional column vector with elements *a*_1_, … , *a*_*n*_. Note that if the elements of a block matrix (or vector) are *not* separated by commas, then no stacking operation is implied.
- Vectors are assumed to be columns unless noted otherwise.
- For vectors **v** and matrices **A, v**′ and **A**′ denote the respective transposes.
- Whenever a quantity **A** is a function of estimates 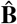, **Ĉ** of (scalar, vector, or matrix) quantities **B, C**, we simplify the notation by omitting the hat from the partial derivatives of **A**. That is, we write

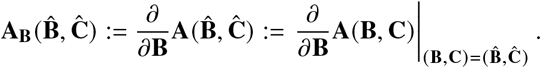 Additionally, wherever it is clear from the context, **A**(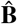 , **Ĉ** ) may be abbreviated as **Â** . Using this notation, we write

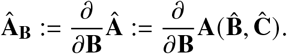
- The (*n* × *n*)-dimensional identity matrix is denoted by **I**_*n*_ and the respective (*n* × *m*)-dimensional matrices of zeros is denoted by **0**_*n*×*m*_; we also write **0**_*n*_ := **0**_*n*×*n*_.

### S3 Derivation of the second stage EM algorithm

In the second step of the GTS method, an EM algorithm is used to estimate the population parameters **b** and **D**, treating the first-stage individual estimates 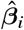 as data and the unknown *β*_*i*_ as latent variables. Though this EM algorithm has been frequently mentioned in the NLME literature over the years, we have been unable to find the calculations behind the EM steps outlined in Methods. We therefore provide a derivation of these steps below.

The EM algorithm starts with the complete-data likelihood function of **b, D**, denoted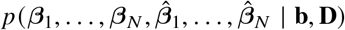. In the E-step, we compute the expected value 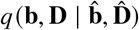 of the complete-data log-likelihood with respect to the conditional distribution of *β*_1_, … , *β*_*N*_ given 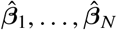 and the current estimates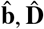. Symbolically,

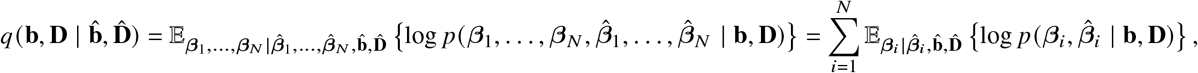

where we used the conditional independence of individual estimates to factorize the likelihood.

At the M-step, we maximize 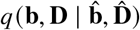 with respect to **b** and **D** to obtain updated estimates 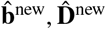.That is,

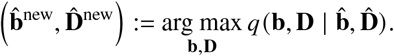

To derive the conditional distribution of each *β*_*i*_ given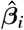, we use Bayes’ theorem together with (11) to write

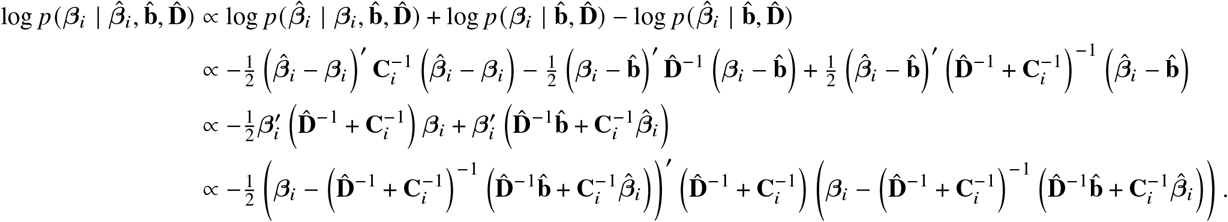

Therefore, the conditional distribution of *β*_*i*_ given 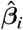 is normal with

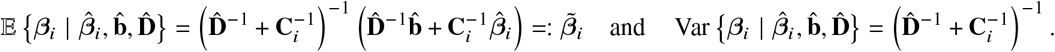

To calculate the expectation of the complete-data log-likelihood with respect to this conditional distribution, we make use of the fact that, for a normally distributed random vector **z** and a symmetric matrix **A** of conforming dimension, 𝔼{**z**′**Az**} = 𝔼{**z**′}**A** 𝔼{**z**} + tr(**A**Var **z**) [Kendrick, 2002, App. B.1]. Therefore, we obtain

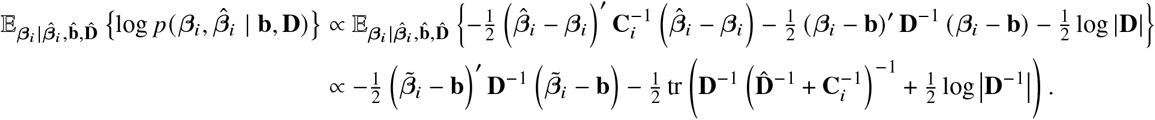

Consequently,

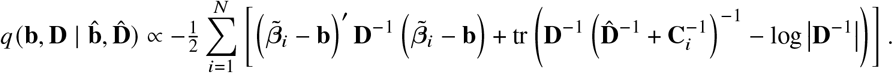

To derive the M-step update, note 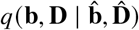 has partial derivatives [Petersen and Pedersen, 2008, Ch. 2]

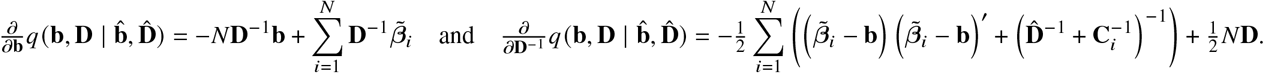

The first-order conditions 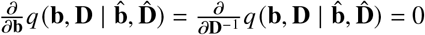 imply

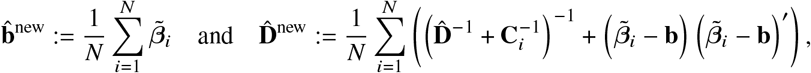

and the second-order conditions are readily verified to confirm 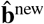 and 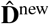 maximize 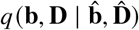 with respect to **b** and **D**.

### S4 Penalized B-spline smoothing

As in Methods, we drop the subscripts *i* for readability, writing **x** := **x**_*i*_ and **y** := **y**_*i*_ (see (8) and (6)) for each statement that applies to every individual independently. Denote the number of observed states by *L*, so that *L* ⩽ *K*, and assume without loss of generality that the first *L* states are observed. For the *i*-th cell, we estimate each observed component of *x*_*k*_ (·) for *k* = 1, … , *L* using a basis expansion of an *M*-dimensional B-spline basis {*B*_1,*v*+1_(·), … , *B*_*M*,*v*+1_(·)} of degree *v* with *M* − *v* − 1 internal knots *k*_1_, … , *k*_*M*−*v*−1_ ∈ (*t*_1_, *t*_*T*_ ). Denote

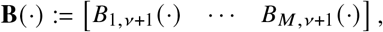

and let ***δ***_*k*_ be an *M*-dimensional vector of coefficients for each *k* = 1, … , *L*. We can then approximate

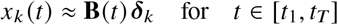

for each *k* = 1, … , *L*. Collecting all basis elements in the block matrix **N**(*t*) = **I**_*L*_ ⊗ **B**(*t*) and the coefficient vectors in ***δ*** := [***δ***_1_, … , ***δ***_*L*_], we can approximate the observed states **x**^*O*^ (*t*) (= **x**(*t*) when *L* = *K*) as

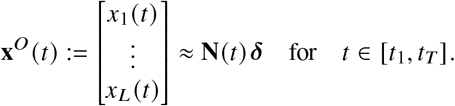

Plugging in the measurement time points in **x**^*O*^ (*t*) and **N**(*t*), we obtain the matrices **X**^*O*^ := [**x**^*O*^ (*t*_1_), … , **x**^*O*^ (*t*_*T*_ )] and **Z** := [**N**(*t*_1_), … , **N**(*t*_*T*_ )], which are related via

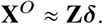

This approximation suggests the following (weighted) linear regression framework for estimating the spline coefficients: let **Y** = [**y**(*t*_1_), … , **y**(*t*_*T*_ )] be the combined (*T L*)-dimensional vector of observed data. Then,

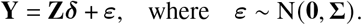

The diagonal error covariance matrix Σ is determined by (7), i.e.

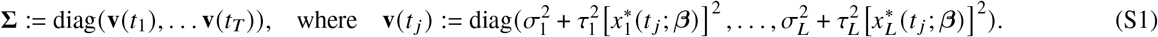

To arrive at a penalized least-squares estimate of the spline coefficients, we consider a penalty multiplier λ_*k*_ > 0 for each state, and denote Λ = diag (λ_1_, … , λ_*L*_ ) ⊗ **I**_*M*_. We then estimate the combined coefficient vector ***δ*** by minimizing the penalized sum of squared differences

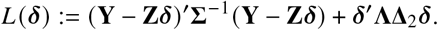

In the above formula, the choice of roughness penalty determines the structure of the matrix Δ_2_. Concretely, to enforce spline smoothness, we consider two alternative options for Δ_2_:

1. Penalize the second derivative of the resulting splines, in which case 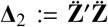 where 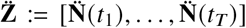 and 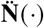 contains the second derivatives of the B-spline basis functions contained in **N**(·);
2. Penalize the second-order differences of the spline coefficients, by letting ***D***_2_ be the ( *M* − 2) × *M* matrix given by

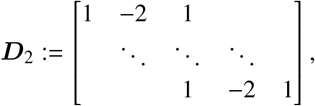

and letting Δ_2_ := **I**_*L*_ ⊗ ***D***^′^ ***D***_2_ [Eilers and Marx, 1996].

We use the second option in our test since it has a higher tendency to produce straight lines. However, the edge effects it produces are more pronounced, so the first option is more suitable if high accuracy near the measurement limits is desired. Instead of penalizing the roughness of entire trajectories, the roughness of the splines over certain subintervals may be penalized instead by premultiplying 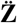 or ***D***_2_ with a suitable diagonal matrix before computing Δ_2_. This approach is beneficial when smoothing single-cell data with fast and slow dynamics, where a single roughness penalty may result in over-/under-smoothing of trajectory segments. In particular, this flexibility was necessary to correctly smooth the data simulated from the bifunctional two-component signaling cascade (cf. Results).

The smoothing parameters λ_1_, … , λ_*L*_ are estimated by minimizing (see Remark S1) the generalized cross-validation error

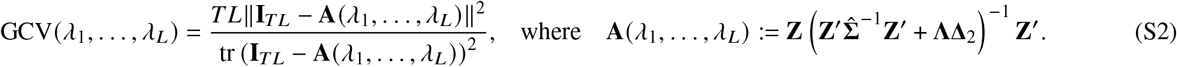

Since Σ (the measurement noise covariance) is generally unknown, it has to be estimated in a feasible weighted least squares (WLS) scheme. Given Λ and an estimate 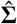 of Σ, the WLS estimate of ***δ*** becomes

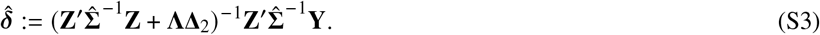

The log-likelihood of each (σ_*k*_, τ_*k*_) is given by (10) with 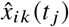 replacing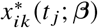. To find the corresponding maximum likelihood estimate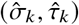, we initialize it at the non-negative least squares estimate

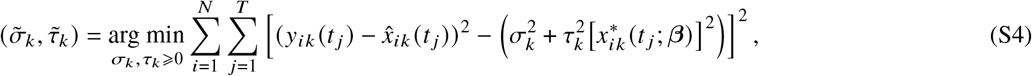

and subsequently optimize the log-likelihood of (σ_*k*_ , τ_*k*_) using an interior-point algorithm. After alternating the estimation of ***δ*** and Σ until convergence, we approximate

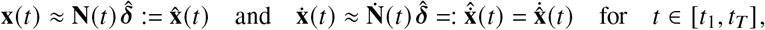

where 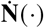 contains the time derivatives of the B-spline basis functions contained in **N** (·) .

#### Remark S1.

Since measured time series from different cells usually do not display high variability in terms of curvature, it often suffices to minimize the GCV for a small set of randomly chosen cells. From the set of minimized state-specific penalty multipliers, state-specific averages may then be computed and used for all cells.

### S5 First-stage uncertainty estimates for fully observed systems

Up to this point, state values across measurement time points have been organized in block vectors and block matrices using *T* blocks of *K* or *L* states (for fully and partially observed systems, respectively). However, for this section, we will use block matrices with *K* or *L* blocks of *T* time points. This rearrangement aids clarity since we will frequently need to distinguish between hidden and observed states. Concretely, we define

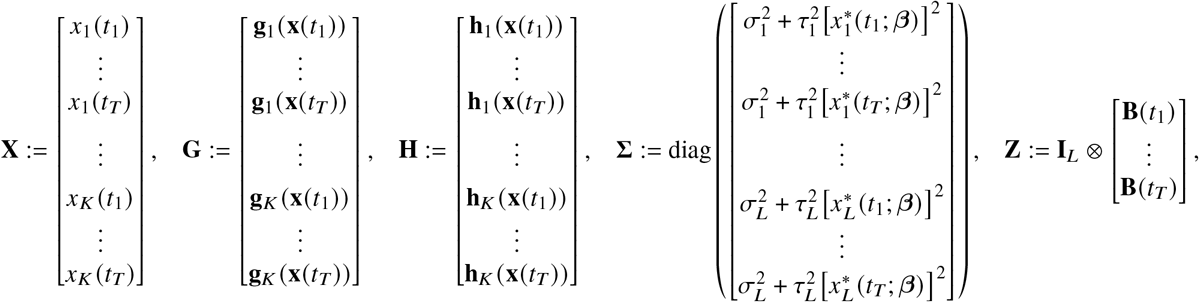

and consider analogous rearrangements for the estimates and the derivatives of the quantities defined above. Similarly to above, ***δ***, 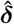, and Λ are organized into *L* blocks of *M* basis functions (as in SI, § S4). Note that the GLS estimate expression (16) is invariant to the applied reordering.

To infer the parameters of the generalized linear model (14), the covariance matrix of Δ needs to be estimated alongside the parameter vector at each iteration of the generalized least squares algorithm. For this calculation, we use the definitions of **Z**, 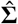, and Λ, and Δ_2_ provided in SI, § S4, but reordered as stated above. Denote by Ω the variance of the spline coefficient vector, which is obtained from (S3):

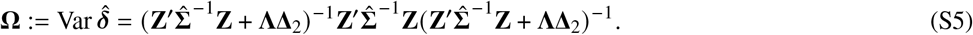

Then, approximately, the error between the smoothed and actual states and their derivatives is given by [Zhou et al., 1998]

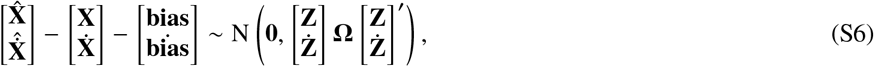

where 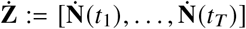 and N (μ, Σ) denotes a multivariate normal distribution with mean μ and covariance Σ. We ignore the bias since there is no way to reliably estimate it (since it depends on derivatives of the unknown ODE solution), but it vanishes as the number of time points increases [Wu et al., 2012]. Treating § 5.5] together with (S6), we obtain

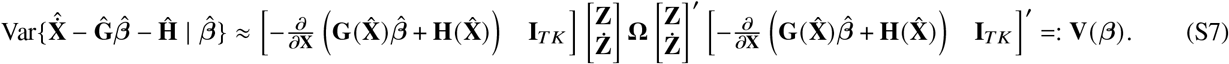

From the definitions at the start of this section, one can see that **V (***β)* is organized into *K* × *K* square blocks of size *T* × *T* . Upon convergence of the FGLS algorithm presented in Methods, we use the covariance matrix of the splines and their gradients to obtain an estimate of the uncertainty of 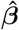 . To this end, the delta-method approximation (17) provides better accuracy compared to the inverse Fisher information matrix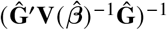. This is because the splines introduce uncertainty to the design matrix 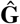that is not accounted for in the Fisher information matrix.

Combining (17) and (S6) we get

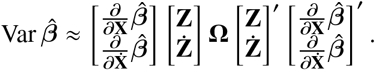

Recall that when the GSS criterion is not maximized at the parameter boundary (cf. Remark 5), 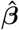 is given by (16), where 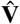 is obtained using (S7) with the previous estimate of *β*. To estimate the uncertainty of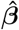, we need to calculate its partial derivatives with respect to the state and gradient estimates. From (16), it is straightforward to see that

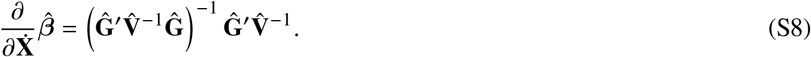

To compute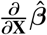, we apply the product rule to (16), i.e.,

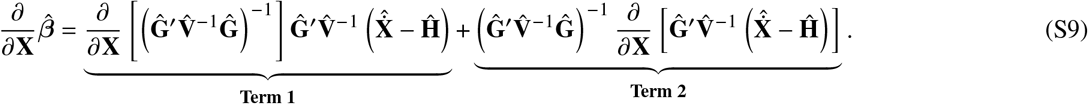

[**Calculation of Term 1**] We start from the fact that [Petersen and Pedersen, 2008, § 2.2]

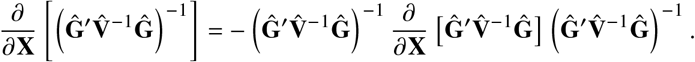

Recall 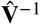 is organized into *K* × *K* blocks of dimension *T* × *T* . Let 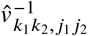 denote the element corresponding to the *j*_1_, *j*_2_-th time point within the *k*_1_, *k*_2_-th block of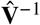. Moreover, denote the *k*-th row of **g**(·) by **g**_*k*_(·) , and the respective partial derivatives of **g**(·) and **g**(·) with respect to *x* by 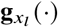 and 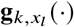. Then

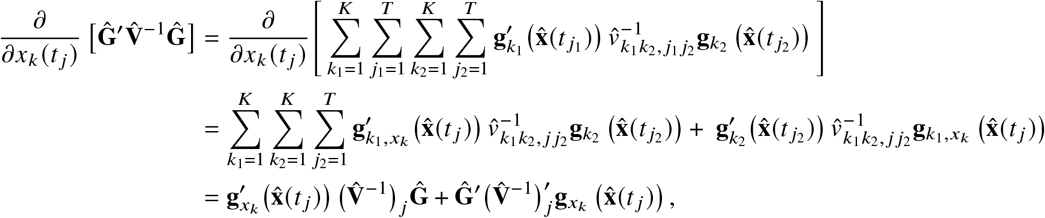

where 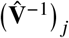 is the submatrix of 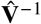 with only the *j* -th row of each *T* × *T*-dimensional block. Denote

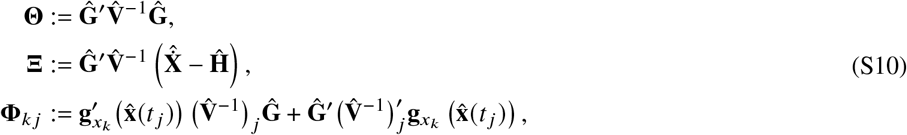

then

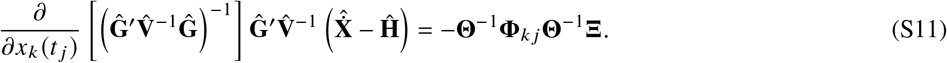

Horizontally stacking (S11) across time points and states, we arrive at

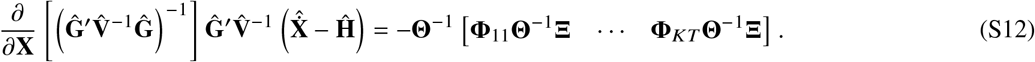

[**Calculation of Term 2**] Similarly,

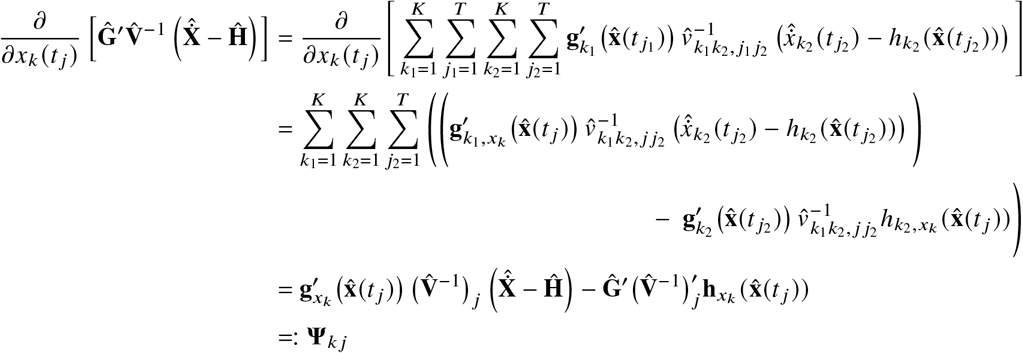

This means

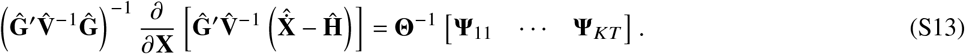

Combining (S12) and (S13), we obtain

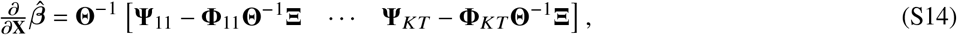

and thus an explicit formula to approximate (17).

### S6 First-stage uncertainty estimate derivation for partially observed systems

Without loss of generality, we assume the last *K* − *L* states are hidden, so **X**^*O*^ simply consists of the first *L* blocks of **X**. Analogously, the vector **X**^*H*^ containing the time point blocks for the hidden states consists of the last *K* − *L* blocks of **X**. The same holds for estimates, gradients, and estimates of gradients of **X**^*O*^ and **X**^*H*^ . The goal of this section is to reconcile the residual covariance expression (S7) with the treatment of partially observed systems. To this end, uncertainty needs to be propagated in accordance with the iterative scheme presented in Methods. The flow of information, and thus of uncertainty, in the algorithm is depicted in Figure S17.

Our iterative estimation scheme starts with smoothed observed states 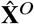 (with gradients 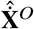) from the estimated spline coefficients 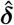 and an initial estimate 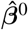 of the cell-specific parameters. The spline coefficients and initial cell-specific parameter estimates come with their estimated (or guessed) covariances Var 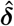, cf. (S5), and Var 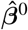. Therefore, the smoothed observed states and their derivatives have their own covariance matrix

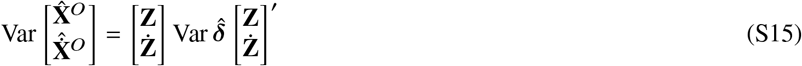

that remains fixed throughout iterations.

Following the definition of the initial cell-specific parameter estimates 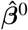 , we integrate the system (8) to obtain the vectors of observed and hidden states, 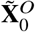 and 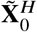 . With respect to notation, from this point on, we use a hat over a symbol to indicate estimated quantities, while a tilde exclusively corresponds to state predictions produced by numerical integration of the system (8) using the current parameter estimate. Since the only way to obtain estimates of the hidden states and their derivatives is via numerical integration, we use the tilde and hat notations interchangeably for those states, i.e.,

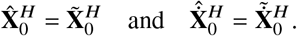

For the observed states, however, spline smoothing estimates 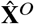 and 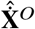 are available, so in general,

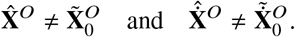

We can combine the estimated hidden and observed states and their gradients into

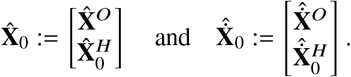

As discussed above, the covariance matrix describing the uncertainty in 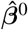 needs to be initialized at the start of the iterations, and before any state, gradient or residual covariance is calculated. We assume the initial parameter estimate 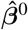 approximates the fixed effect vector **b**, and thus initialize its uncertainty with a guess of the random effect covariance matrix; see Remark S2 at the end of this section.

To compute the equivalent of (S7) for the partial observation case, we first need an estimate of Var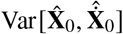. Note that the hidden part of 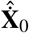 depends on 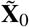 and 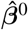. Let ***F* (***β*) denote the vector of solution values of (8) given the cell-specific parameter vector *β*, that is,

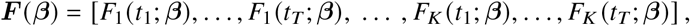

where *F*_*k*_ (*t* _*j*_ ; ***θ***) denotes the *k*-th component of the ODE solution at time *t* _*j*_ . Denote the respective observed and hidden parts by ***F*** ^*O*^ (*β*) (first *L* blocks of ***F*** (*β*)) and ***F*** ^*H*^ (*β*) (last *K* − *L* blocks of ***F*** (*β*)). Finally, let 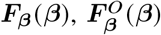, and 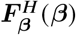 be the corresponding partial derivatives with respect to *β*, which may be obtained by numerically solving the sensitivity equations of (8) or through finite difference approximations. Then, using the delta method, the full covariance matrix of 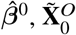 , and 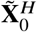 is given by

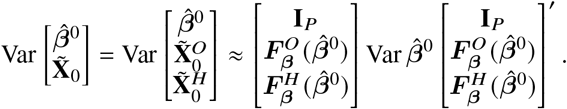

We can now compute Var 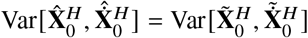 from the right-hand side of (8) and the preceding covariance matrix as

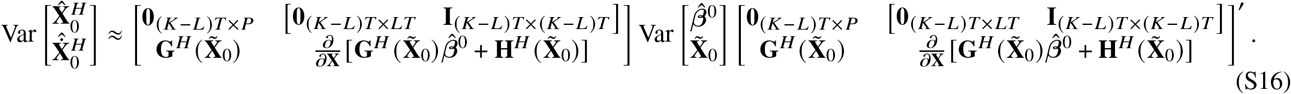

Now that we have derived approximate covariances for the hidden states and their gradients, we can obtain a covariance matrix estimate for the full set of states and gradients. Since 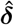 and 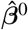 were estimated independently,

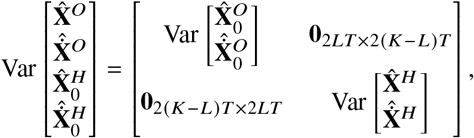

Reorganizing row and column blocks, we obtain an approximation for Var 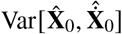, allowing us to compute 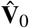 analogously to (S7) (specifically, according to (S17) presented below, with *J* = 0).

From this point onwards, we consider expressions at a general iteration *J* ⩾ 0. Given an (approximate) covariance matrix Var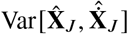, we use another delta method approximation to obtain 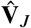as

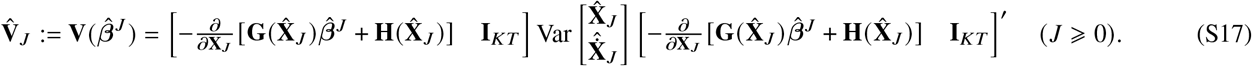

Approximating Var 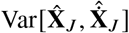 is more involved for *J* ⩾ 1, however, since estimates and predictions of states and gradients are used to obtain 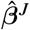 in a gradient matching step, as presented in Methods. Specifically, 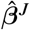 is given by

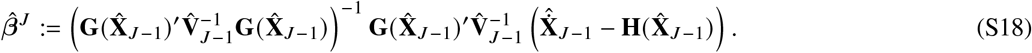

Analogously to (16), this relation holds when the estimate 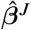 is not found on the parameter boundary (see Remark 5).

The updated parameter estimate is subsequently used to produce new hidden state and gradient predictions. As illustrated in Figure S17, this fact implies that uncertainty propagates from 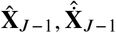 to 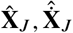 through 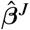 . Therefore, we may consider 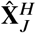 and 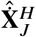 as functions of 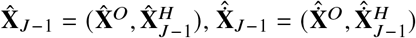 and 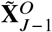 . Then, the relevant sensitivities for 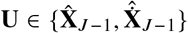 follow from the chain rule as

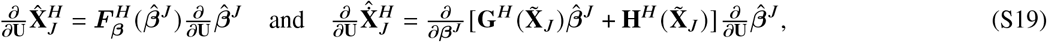

where 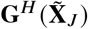 and 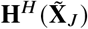 consist of the last *K* − *L* blocks of 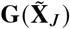 and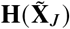. The partial derivatives of 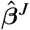 with respect to the gradient and state estimates derive from (S8) and (S14), respectively. Putting everything together, we find

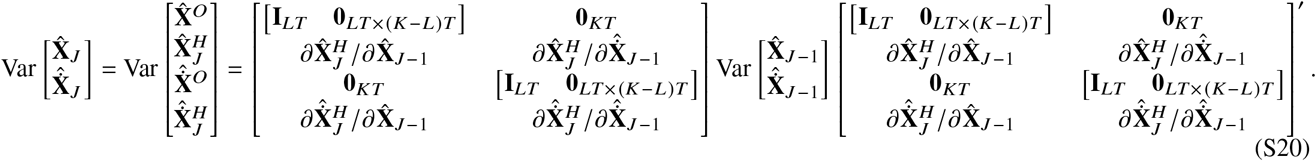

As with full state information, upon convergence of the iterative algorithm presented in Methods, we can compute the variance of 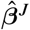 using

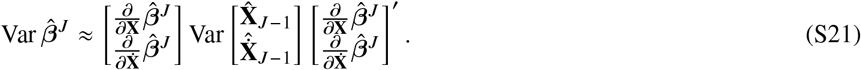

#### Remark S2.

When the initial parameter estimate 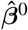 is produced by fitting the mean trajectory across cells, it is more reasonable to consider 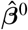 as a (biased) estimate of the fixed effects vector **b** (see SI, § S8), but not as approximating any individual parameter vector *β*. Assuming that 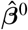 approximates **b**, it is thus best to approximate the covariance of 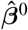 with an initial estimate of the random effects covariance matrix, rather than with the (inverse) Hessian matrix of the objective function that was minimized to obtain 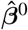. This is because the precision implied by the Hessian does not in general correspond to the precision of 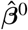 as an estimate of any cell-specific parameter vector. To see why this the case, assuming that 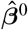 approximates **b**, we get

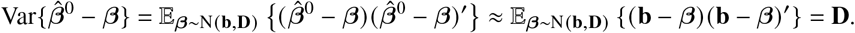

In case it is not possible to obtain reasonable orders of magnitude for the **D** a priori, it may be necessary to try a couple of different values if subsequent individual parameter estimates 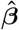 stay too close to 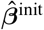.

### S7 Including prior information and modifications for maximum a posteriori estimates

In some situations, gradient matching will tend produce practically infeasible estimates, which thereby end up on the parameter boundary. When this happens, the uncertainty computations in SI, § S5–S6 become inaccurate, which may result in severe inaccuracy when inferring the random effects distribution in the second stage of the GMGTS method. In such cases, besides setting parameter boundaries, information on reasonable parameter values should be incorporated by including a prior parameter distribution.

Suppose a common prior N(*β*^*^, **C**^*^) for *β*_1_, … , *β*_*N*_ is included, then the GSS becomes

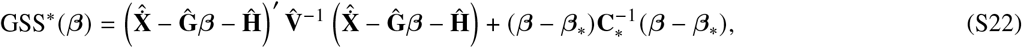

Where (15) was minimized at the maximum likelihood estimate, the estimate that maximizes GSS^*^ *β* is referred to the *maximum a posteriori* (MAP) estimate. It is straightforward to verify the MAP estimate for each *β* becomes

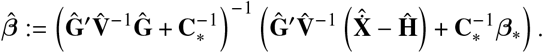

Since *β*^*^ and **C**^*^ do not depend on any state or gradient estimates, very few adjustments are needed to the uncertainty computations in SI, § S5–6. It immediately follows that

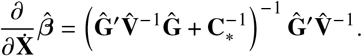

In the computation of 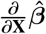, only undifferentiated factors change. Specifically, Θ and Ξ in (S10) need to be modified, i.e.,

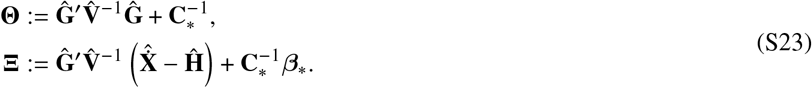

### S8 Implementation details

This section outlines the software implementation details of the methodology presented in Methods and SI, § 4–6. Note that cell subscript indices were omitted in some of the referenced equations for ease of reading; they are included here wherever applicable. The notes can serve as guidelines for custom implementation efforts; for a user-ready implementation in Matlab, see [van Oppen, 2023].

#### Input processing

We used the Symbolic Math Toolbox of Matlab to implement rudimentary preprocessing of a user-supplied IQM Tools model file containing differential equations, initial values, and nominal parameter values. From this file, our preprocessing code extracts the right-hand side **g** and **h** functions from (8), as well as their partial derivatives with respect to the system states. This preprocessing is necessary to avoid the error-prone process of manually defining the different matrix-valued functions required by GMGTS. Since symbolic computation is slow, the derived symbolic functions are vectorized to handle multiple cells and time points and converted to regular functions. Doing so generates a small amount of overhead, so function calls are made for all cells at all time points rather than single cells or time points.

#### Smoothing

All measurement noise parameters are initialized at σ_*k*_ = 1 and τ_*k*_ = 0; that is, we start by assuming homoskedastic errors. The spline basis and smoothness penalty matrices are precomputed for computational efficiency. Then, the smoothing proceeds iteratively:

1. Update the penalty multipliers λ_*k*_ by minimizing the GCV error (S2). As noted in Remark S1, in most practical cases, it suffices to optimize each λ_*k*_ for a small set of randomly selected cells.
2. Update the spline coefficient vector estimate 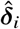 using (S3) and compute the corresponding estimated state values 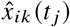.
3. Optimize the log-likelihood (10) with 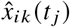 in place of 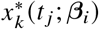 to update each 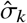 and 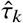, initializing at the non-negative least squares estimate (S4). In our Matlab implementation, we used the interior point algorithm provided by fmincon for the log-likelihood maximization.
4. Repeat steps 1–3 until convergence.

When the procedure ends, use 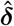 to compute each 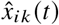 and 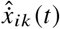 on a minimal set of time points *t* that still capture the dynamics of the system. Also compute (S1) using 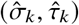 and with 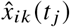 in place of 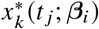.

#### Individual estimates with full observation

Initialize the residual covariance matrices Var Δ_*i*_ as *T K* × *T K* identity matrices, starting the GLS iterations with homoskedastic errors. Then,

1. Update each 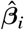 by minimizing the GSS criterion in (15) using quadratic programming to conform with parameter bounds. Note the smoothed states and gradients at the first and last time point should be omitted as they often carry more bias than other points.
2. Compute Var 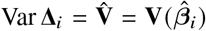 using (S7). Generally speaking, this matrix will be rank-deficient due to the small number of knots versus time points. Wherever we need its inverse, we supply its Moore-Penrose pseudoinverse [Horn and Johnson, 2012] instead. For numerical stability, we compute the pseudoinverse using the singular value decomposition of 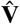, ignoring singular values smaller than a set tolerance. When there are large magnitude differences between states or gradients, 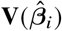 needs to be scaled accordingly before computing its pseudoinverse, and the pseudoinverse needs to be scaled back. Failing to rescale 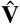 may cause singular values corresponding to states and gradients of smaller magnitude to be ignored when computing the pseudoinverse.
3. Repeat steps 1 and 2 until convergence. Since the random effects distribution rather than the individual random effects is the object of interest in our method, it suffices to use a high cell-specific convergence tolerance (for instance, less than 0.01 relative change in the magnitude of the parameter vector) and to terminate after a set number (e.g., 5–10) iterations or when the majority (e.g., 90%) of the cell estimates have converged. While cell-specific parameter estimates can in principle be computed sequentially, our preprocessing step makes it significantly more efficient to make ODE-related function calls in batches. In this way, we simultaneously update all individual parameter estimates that have not converged at each step.

#### Individual estimates with partial observation

For the iterative estimation scheme described above, we initialize all cell-specific parameter estimates 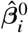 at the same value. To do this, we minimize the discrepancy between the ODE model prediction for a single parameter value and the mean trajectory data across cells. That is, we estimate

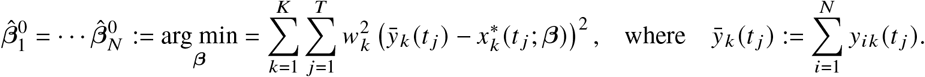

To avoid unwanted additional sensitivity with respect to the smoothing, the state-specific weights *w*_*k*_ in the sum of squares above are not derived from estimates of the cell-specific measurement errors. Instead, they may simply be taken to all equal 1 or be inversely proportional to the state-specific measurement averages across cells and time points.

To perform the minimization, we use the interior point algorithm provided by fmincon in Matlab, multistarted at 5–20 points randomly sampled within the parameter bounds. As noted before, Var 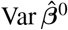 should be set using the initial random effects covariance matrix. We have found that, in the absence of prior knowledge, a conservative estimate using a diagonal matrix with a CV of *c* = 0.05 or *c* = 0.1 suffices. Specifically, in this case we initialize

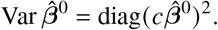

As discussed above, contrary to the case of full observation, we first produce estimates of the hidden states by integrating the ODE system for each cell using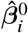. We then estimate the initial residual covariance matrix 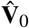 using (S17), store the intermediate estimate of Var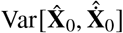, and at each iteration *J* = 1, 2, …,

1. As with full state information, compute each updated 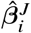 by minimizing the GSS criterion in (15).
2. Integrate the system using the current estimates 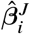 to compute updated hidden state and gradient estimates 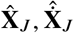.
3. Compute the updated 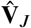 using (S17). To do so, first compute the updated estimate of Var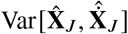 using (S14), (S19), (S20), and the saved value of Var 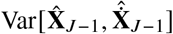 from the previous iteration. When using maximum a posteriori estimates in Step 1, take note of the modifications listed in (S23).
4. Analogously to the case of full observation, repeat steps 1–3 until convergence or terminations. Upon termination, approximate Var 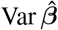 using (S21).

#### Global Two-Stage settings and implementation

To facilitate performance comparisons, we aimed to implement the GTS approach as efficiently as possible and used the same estimation settings as for the GMGTS method wherever applicable. The overall structures of the two methods are very similar. For GMGTS, an iteration in the first stage amounts to a gradient matching step using linear regression (and integrating the system in case of partial observation) followed by estimation of the residual covariance matrix. For the GTS method, an iteration comprises an application of a non-convex optimization routine followed by estimation of the noise variance parameters. The second stages of the two methods are identical.

Similar to GMGTS with partial observation, the first stage individual parameter estimates of GTS are initialized by matching the average of the single-cell measurements using multi-started optimization, yielding a common initial estimate for all cells. At each subsequent iteration, the optimization routine that updates individual parameter estimates is initialized at the previous estimate. To terminate the optimization of each individual parameter estimate, an optimality tolerance of 1E-6 was used. Finally, a tolerance of 1E-3 was used for estimating measurement noise parameters via optimization, in a manner analogous to the one described at the end of SI, § S4. Wherever numerical optimization was needed, we used the interior-point method provided by the fmincon function in Matlab.

Common settings between methods were the number of iterations per stage (5 for the first and 10 for the second stage), the relative convergence tolerances of the first and second stage iterations (2E-3), the number of multi-start points (10) for initialization of the first stage iterations, the parameter search spaces, and the prior (whenever included).

Finally, analogously to [Dharmarajan et al., 2019], we approximated the uncertainty of each first-stage estimate with the inverse of the Fisher information matrix, ignoring second-order terms. This approximation is based on ODE solution sensitivities with respect to the individual parameter estimate. For the calculation of these sensitivities, finite-difference approximations turned out to be more computationally efficient than solving the forward sensitivity equations in IQM Tools and Monolix.

## REFERENCES

[Albeck et al., 2008] Albeck, J. G., Burke, J. M., Aldridge, B. B., Zhang, M., Lauffenburger, D. A., and Sorger, P. K. (2008). Quantitative analysis of pathways controlling extrinsic apoptosis in single cells. Molecular Cell, 30(1):11–25. 1

[Aldridge et al., 2006] Aldridge, B. B., Burke, J. M., Lauffenburger, D. A., and Sorger, P. K. (2006). Physicochemical modelling of cell signalling pathways. Nature Cell Biology, 8(11):1195–1203. 1

[Almquist et al., 2015] Almquist, J., Bendrioua, L., Adiels, C. B., Goksör, M., Hohmann, S., and Jirstrand, M. (2015). A nonlinear mixed effects approach for modeling the cell-to-cell variability of Mig1 dynamics in yeast. PLoS ONE, 10(4):e0124050. 1

[Asmussen and Glynn, 2007] Asmussen, S. and Glynn, P. W. (2007). Stochastic simulation: algorithms and analysis, volume 57. Springer. 3

[Balleza et al., 2018] Balleza, E., Kim, J. M., and Cluzel, P. (2018). Systematic characterization of maturation time of fluorescent proteins in living cells. Nature Methods, 15(1):47–51. 8

[Barber and Wang, 2014] Barber, D. and Wang, Y. (2014). Gaussian processes for Bayesian estimation in ordinary differential equations. In International Conference on Machine Learning, pages 1485–1493. PMLR. 11

[Battich et al., 2015] Battich, N., Stoeger, T., and Pelkmans, L. (2015). Control of transcript variability in single mammalian cells. Cell, 163(7):1596–1610. 1

[Bijman et al., 2021] Bijman, E. Y., Kaltenbach, H.-M., and Stelling, J. (2021). Experimental analysis and modeling of single-cell time-course data. Current Opinion in Systems Biology, 28:100359. 1

[Butcher and Tabor, 2022] Butcher, R. J. and Tabor, J. J. (2022). Real-time detection of response regulator phosphorylation dynamics in live bacteria. Proceedings of the National Academy of Sciences, 119(35):e2201204119. 5

[Calderhead et al., 2009] Calderhead, B., Girolami, M., and Lawrence, N. D. (2009). Accelerating Bayesian inference over nonlinear differential equations with Gaussian processes. In Advances in Neural Information Processing Systems, pages 217–224. Citeseer. 2, 11

[Corless et al., 1996] Corless, R. M., Gonnet, G. H., Hare, D. E., Jeffrey, D. J., and Knuth, D. E. (1996). On the Lambert W function. Advances in Computational mathematics, 5:329–359. 21

[Davidian and Giltinan, 2003] Davidian, M. and Giltinan, D. M. (2003). Nonlinear models for repeated measurement data: an overview and update. Journal of Agricultural, Biological, and Environmental Statistics, 8:387–419. 1

[Davidian and Giltinan, 2017] Davidian, M. and Giltinan, D. M. (2017). Nonlinear Models for Repeated Measurement Data. Routledge. 1, 3, 10, 15, 16

[Dharmarajan et al., 2019] Dharmarajan, L., Kaltenbach, H.-M., Rudolf, F., and Stelling, J. (2019). A simple and flexible computational framework for inferring sources of heterogeneity from single-cell dynamics. Cell Systems, 8(1):15–26. 1, 3, 11

[Dondelinger et al., 2013] Dondelinger, F., Husmeier, D., Rogers, S., and Filippone, M. (2013). ODE parameter inference using adaptive gradient matching with Gaussian processes. In Artificial Intelligence and Statistics, pages 216–228. PMLR. 2, 11

[Duvall and Childers, 2020] Duvall, S. W. and Childers, W. S. (2020). Design of a histidine kinase fret sensor to detect complex signal integration within living bacteria. ACS Sensors, 5(6):1589–1596. 5

[Efron, 2016] Efron, B. (2016). Empirical Bayes deconvolution estimates. Biometrika, 103(1):1–20. 11

[Elowitz and Leibler, 2000] Elowitz, M. B. and Leibler, S. (2000). A synthetic oscillatory network of transcriptional regulators. Nature, 403(6767):335–338. 6

[Foreman and Wollman, 2020] Foreman, R. and Wollman, R. (2020). Mammalian gene expression variability is explained by underlying cell state. Molecular Systems Biology, 16(2):e9146. 1

[Fröhlich et al., 2018] Fröhlich, F., Reiser, A., Fink, L., Woscheé, D., Ligon, T., Theis, F. J., Rädler, J. O., and Hasenauer, J. (2018). Multi-experiment nonlinear mixed effect modeling of single-cell translation kinetics after transfection. NPJ Systems Biology and Applications, 4(1):42. 1

[Greene, 2003] Greene, W. (2003). Econometric analysis, 5th ed. 15

[Guantes et al., 2015] Guantes, R., Rastrojo, A., Neves, R., Lima, A., Aguado, B., and Iborra, F. J. (2015). Global variability in gene expression and alternative splicing is modulated by mitochondrial content. Genome Research, 25(5):633–644. 1

[Guerra et al., 2022] Guerra, P., Vuillemenot, L.-A., Rae, B., Ladyhina, V., and Milias-Argeitis, A. (2022). Systematic in vivo characterization of fluorescent protein maturation in budding yeast. ACS Synthetic Biology, 11(3):1129–1141. 2, 5, 8, 9, 10, 19, 20

[Huang, 2009] Huang, S. (2009). Non-genetic heterogeneity of cells in development: more than just noise. Development, 136(23):3853–3862. 1

[Karlsson et al., 2015] Karlsson, M., Janzén, D. L., Durrieu, L., Colman-Lerner, A., Kjellsson, M. C., and Cedersund, G. (2015). Nonlinear mixed-effects modelling for single cell estimation: when, why, and how to use it. BMC Systems Biology, 9:1–15. 1

[Keren et al., 2015] Keren, L., Van Dijk, D., Weingarten-Gabbay, S., Davidi, D., Jona, G., Weinberger, A., Milo, R., and Segal, E. (2015). Noise in gene expression is coupled to growth rate. Genome Research, 25(12):1893–1902. 1

[Kramer et al., 2022] Kramer, B. A., Sarabia del Castillo, J., and Pelkmans, L. (2022). Multimodal perception links cellular state to decision-making in single cells. Science, 377(6606):642–648. 1

[Kuhn and Lavielle, 2005] Kuhn, E. and Lavielle, M. (2005). Maximum likelihood estimation in nonlinear mixed effects models. Computational Statistics & Data Analysis, 49(4):1020–1038. 1

[Kurdyaeva and Milias-Argeitis, 2021] Kurdyaeva, T. and Milias-Argeitis, A. (2021). Uncertainty propagation for deterministic models of biochemical networks using moment equations and the extended kalman filter. Journal of the Royal Society Interface, 18(181):20210331. 5, 19

[Levien et al., 2021] Levien, E., Min, J., Kondev, J., and Amir, A. (2021). Non-genetic variability in microbial populations: survival strategy or nuisance? Reports on Progress in Physics, 84(11):116601. 1

[Lindstrom and Bates, 1990] Lindstrom, M. J. and Bates, D. M. (1990). Nonlinear mixed effects models for repeated measures data. Biometrics, pages 673–687. 1

[Llamosi et al., 2016] Llamosi, A., Gonzalez-Vargas, A. M., Versari, C., Cinquemani, E., Ferrari-Trecate, G., Hersen, P., and Batt, G. (2016). What population reveals about individual cell identity: single-cell parameter estimation of models of gene expression in yeast. PLoS Computational Biology, 12(2):e1004706. 1, 3

[Locke and Elowitz, 2009] Locke, J. C. and Elowitz, M. B. (2009). Using movies to analyse gene circuit dynamics in single cells. Nature Reviews Microbiology, 7(5):383–392. 1

[Loos and Hasenauer, 2019] Loos, C. and Hasenauer, J. (2019). Mathematical modeling of variability in intracellular signaling. Current Opinion in Systems Biology, 16:17–24. 1

[Macdonald and Husmeier, 2015] Macdonald, B. and Husmeier, D. (2015). Gradient matching methods for computational inference in mechanistic models for systems biology: a review and comparative analysis. Frontiers in Bioengineering and Biotechnology, 3:180. 2, 3

[Muzzey and Van Oudenaarden, 2009] Muzzey, D. and Van Oudenaarden, A. (2009). Quantitative time-lapse fluorescence microscopy in single cells. Annual Review of Cell and Developmental, 25:301–327. 1

[Neymotin et al., 2016] Neymotin, B., Ettorre, V., and Gresham, D. (2016). Multiple transcript properties related to translation affect mRNA degradation rates in Saccharomyces cerevisiae. G3: Genes, Genomes, Genetics, 6(11):3475–3483. 5, 9

[Paulsson, 2005] Paulsson, J. (2005). Models of stochastic gene expression. Physics of Life Reviews, 2(2):157–175. 1

[Pinheiro et al., 2022] Pinheiro, S., Pandey, S., and Pelet, S. (2022). Cellular heterogeneity: Yeast-side story. Fungal Biology Reviews, 39:34–45. 1

[Raj and Van Oudenaarden, 2008] Raj, A. and Van Oudenaarden, A. (2008). Nature, nurture, or chance: stochastic gene expression and its consequences. Cell, 135(2):216–226. 1

[Ramsay et al., 2007] Ramsay, J. O., Hooker, G., Campbell, D., and Cao, J. (2007). Parameter estimation for differential equations: a generalized smoothing approach. Journal of the Royal Statistical Society Series B: Statistical Methodology, 69(5):741–796. 2, 11

[Raser and O’shea, 2005] Raser, J. M. and O’shea, E. K. (2005). Noise in gene expression: origins, consequences, and control. Science, 309(5743):2010–2013. 1

[Särkkä and Svensson, 2023] Särkkä, S. and Svensson, L. (2023). Bayesian filtering and smoothing, volume 17. Cambridge University Press. 11

[Sherman et al., 2015] Sherman, M. S., Lorenz, K., Lanier, M. H., and Cohen, B. A. (2015). Cell-to-cell variability in the propensity to transcribe explains correlated fluctuations in gene expression. Cell Systems, 1(5):315–325. 1

[Spencer et al., 2009] Spencer, S. L., Gaudet, S., Albeck, J. G., Burke, J. M., and Sorger, P. K. (2009). Non-genetic origins of cell-to-cell variability in trail-induced apoptosis. Nature, 459(7245):428–432. 1

[Spiller et al., 2010] Spiller, D. G., Wood, C. D., Rand, D. A., and White, M. R. (2010). Measurement of single-cell dynamics. Nature, 465(7299):736–745. 1

[van Oppen, 2023] van Oppen, Y. (2023). GMGTS Matlab implementation. https://github.com/yulanvanoppen/GMGTS. 11, 18

[Varah, 1982] Varah, J. M. (1982). A spline least squares method for numerical parameter estimation in differential equations. SIAM Journal on Scientific and Statistical Computing, 3(1):28–46. 2, 3, 11, 16

[Villani, 2009] Villani, C. (2009). The Wasserstein distances. In: Optimal Transport, pages 93–111. Springer Berlin Heidelberg, Berlin, Heidelberg. 6

[Waldherr et al., 2009] Waldherr, S., Hasenauer, J., and Allgöwer, F. (2009). Estimation of biochemical network parameter distributions in cell populations. IFAC Proceedings Volumes, 42(10):1265–1270. 1

[Wu et al., 2020] Wu, J., Han, X., Zhai, H., Yang, T., and Lin, Y. (2020). Evidence for rate-dependent filtering of global extrinsic noise by biochemical reactions in mammalian cells. Molecular Systems Biology, 16(5):e9335. 2, 9

[Zopf et al., 2013] Zopf, C., Quinn, K., Zeidman, J., and Maheshri, N. (2013). Cell-cycle dependence of transcription dominates noise in gene expression. PLoS Computational Biology, 9(7):e1003161. 1

## References

[Berger and Casella, 2001] Berger, R. L. and Casella, G. (2001). Statistical inference. Duxbury.

[Dharmarajan et al., 2019] Dharmarajan, L., Kaltenbach, H.-M., Rudolf, F., and Stelling, J. (2019). A simple and flexible computational framework for inferring sources of heterogeneity from single-cell dynamics. Cell Systems, 8(1):15–26. 1, 3, 11

[Eilers and Marx, 1996] Eilers, P. H. and Marx, B. D. (1996). Flexible smoothing with b-splines and penalties. Statistical Science, 11(2):89–121.

[Horn and Johnson, 2012] Horn, R. A. and Johnson, C. R. (2012). Matrix Analysis. Cambridge University Press.

[Kendrick, 2002] Kendrick, D. A. (2002). Stochastic Control for Economic Models. The University of Texas.

[Petersen and Pedersen, 2008] Petersen, K. B. and Pedersen, M. S. (2008). The matrix cookbook. Technical University of Denmark, 7(15):510.

[Wu et al., 2012] Wu, H., Xue, H., and Kumar, A. (2012). Numerical discretization-based estimation methods for ordinary differential equation models via penalized spline smoothing with applications in biomedical research. Biometrics, 68(2):344–352.

[Zhou et al., 1998] Zhou, S., Shen, X., and Wolfe, D. A. (1998). Local asymptotics for regression splines and confidence regions. The Annals of Statistics, 26(5):1760–1782.

